# Cumulative impacts across Australia’s Great Barrier Reef: A mechanistic evaluation

**DOI:** 10.1101/2020.12.01.406413

**Authors:** Yves-Marie Bozec, Karlo Hock, Robert A. B. Mason, Mark E. Baird, Carolina Castro-Sanguino, Scott A. Condie, Marji Puotinen, Angus Thompson, Peter J. Mumby

## Abstract

Cumulative impacts assessments on marine ecosystems have been hindered by the difficulty of collecting environmental data and identifying drivers of community dynamics beyond local scales. On coral reefs, an additional challenge is to disentangle the relative influence of multiple drivers that operate at different stages of coral ontogeny. We integrated coral life history, population dynamics and spatially-explicit environmental drivers to assess the relative and cumulative impacts of multiple stressors across 2,300 km of the world’s largest coral reef ecosystem, Australia’s Great Barrier Reef (GBR). Using literature data, we characterized relationships between coral life history processes (reproduction, larval dispersal, recruitment, growth and mortality) and environmental variables. We then simulated coral demographics and stressor impacts at the organism (coral colony) level on >3,800 individual reefs linked by larval connectivity, and exposed to temporally- and spatially-realistic regimes of acute (crown-of-thorns starfish outbreaks, cyclones and mass coral bleaching) and chronic (water quality) stressors. Model simulations produced a credible reconstruction of recent (2008–2020) coral trajectories consistent with monitoring observations, while estimating the impacts of each stressor at reef and regional scales. Overall, corals declined by one third across the GBR, from an average ∼29% to ∼19% hard coral cover. By 2020, less than 20% of the GBR had coral cover higher than 30%. Global annual rates of coral mortality were driven by bleaching (48%) ahead of cyclones (41%) and starfish predation (11%). Beyond the reconstructed status and trends, the model enabled the emergence of complex interactions that compound the effects of multiple stressors while promoting a mechanistic understanding of coral cover dynamics. Drivers of coral cover growth were identified; notably, water quality (suspended sediments) was estimated to delay recovery for at least 25% of inshore reefs. Standardized rates of coral loss and recovery allowed the integration of all cumulative impacts to determine the equilibrium cover for each reef. This metric, combined with maps of impacts, recovery potential, water quality thresholds and reef state metrics, facilitates strategic spatial planning and resilience-based management across the GBR.

## INTRODUCTION

The increasing threats faced by marine ecosystems compels us to better understand the cumulative impacts of multiple pressures on species and habitats. Yet, progress towards assessment of multiple stressors has been hindered by the difficulty of characterizing biological responses across ecological scales (Crain et al. 2008, Hodgson and Halpern 2019). Responses to a particular stressor can be complex (e.g., indirect, nonlinear), variable in space and time, and compounded with other stressors or ecological processes (Paine et al. 1998, Darling and Côté 2008). Moreover, one stressor can affect specific life-stages or demographic processes that make interactions with other stressors difficult to detect. Integrated approaches to cumulative impact assessment are required to better predict the ecosystem-level effects of multiple stressors and provide enhanced guidance for the strategic planning and spatial prioritization of management interventions (Halpern and Fujita 2013, Hodgson and Halpern 2019).

The impacts of multiple stressors can be particularly difficult to predict in biogenic habitats (e.g. coral reefs, kelp forests) where acute and chronic pressures simultaneously affect the reproduction, growth and mortality of habitat forming species (Harborne et al. 2017, Filbee-Dexter and Wernberg 2018). This is especially challenging on coral reefs which are deteriorating worldwide due to the compounded effects of natural disturbances with accelerating anthropogenic pressures (Hoegh-Guldberg et al. 2007, Hughes et al. 2017). Whereas extensive coral loss can be easily attributed to acute stressors such as tropical storms, coral bleaching and outbreaks of coral predators (Hughes and Connell 1999, De’ath et al. 2012), identifying the causes of hindered coral recovery is more difficult (Graham et al. 2011, 2015, Osborne et al. 2017, Ortiz et al. 2018). Slow regeneration of coral populations can result from the dysfunction of a range of early-life processes, including reproduction, larval dispersal, settlement and post-settlement growth and mortality (Hughes and Connell 1999, Hughes et al. 2011). The underlying causes can be multiple – such as macroalgal overgrowth, excess sediment and nutrient from land run-off, light reduction in turbid waters (Hughes et al. 2003, Fabricius 2005, Mumby and Steneck 2008, Jones et al. 2015, Evans et al. 2020) – and attribution can be difficult without surveying the relevant life-history stages. Moreover, response to stressors vary among coral species (Loya et al. 2001, Darling et al. 2013) and can lead to complex interactions whose outcomes are difficult to predict (Ban et al. 2014, Bozec and Mumby 2015). As the focus of modern reef management is on promoting local coral recovery in the face of less manageable drivers (e.g., anthropogenic climate warming), cumulative impacts assessments on coral reefs must integrate all stressors across the coral life-cycle.

Australia’s Great Barrier Reef (GBR) exemplifies the challenge of evaluating cumulative pressures on coral reefs, despite being widely considered one of the best studied, monitored and managed reef systems in the world (GBRMPA 2019, although see Brodie and Waterhouse 2012). The GBR Marine Park stretches over 2,300 km across an area of >344,000 km^2^, which means that only a limited fraction of coral reefs can be monitored. Over the past three decades, average coral condition across the GBR has declined in response to the combined impacts of cyclones, outbreaks of the coral-eating crown-of-thorns starfish (*Acanthaster* spp.; CoTS), temperature-induced bleaching and poor water quality (Osborne et al. 2011, De’ath et al. 2012, Hughes et al. 2017, Schaffelke et al. 2017). Much of the research on cumulative impacts on the GBR has used time-series of coral cover to evaluate the rate and drivers of coral loss (Thompson and Dolman 2010, Osborne et al. 2011, Sweatman et al. 2011, De’ath et al. 2012, Cheal et al. 2017). Until recently, coral loss was mostly related to tropical storms and CoTS outbreaks (De’ath et al. 2012), with occasional yet significant impacts of coral bleaching (Berkelmans et al. 2004, Hughes et al. 2017). The two consecutive bleaching events in 2016 and 2017, that caused extensive coral mortality on the northern two-thirds of the GBR (Hughes et al. 2017, 2018, GBRMPA 2019), extend the relative impact and sphere of influence across the GBR. Anthropogenic climate warming and the reducing time interval between severe bleaching events are now considered a major threat for the GBR, hindering its ability to recover from other disturbances and maintain key reef functions (Schaffelke et al. 2017, GBRMPA 2019).

Compared to drivers of coral loss, pressures on coral recovery across the GBR are less well established. While run-off of fine sediments, nutrients and pesticides combine to affect water quality on inshore reefs (Brodie and Waterhouse 2012, Schaffelke et al. 2017, Waterhouse et al. 2017), their demographic impacts on corals remain hard to quantify, likely involving interrelated factors such as a reduction in juvenile densities, increased susceptibility to disease, macroalgal growth and enhanced survival of CoTS larvae (Fabricius and De’ath 2004, Brodie et al. 2005, Fabricius et al. 2010, Thompson et al. 2014). Analyses of monitoring data have related reductions in the rate of coral cover growth with exposure to river plumes (Ortiz et al. 2018, MacNeil et al. 2019) but the underlying mechanisms remain unclear. A number of physiological responses to water quality parameters have been be established experimentally (Fabricius 2005) but quantifying the ecological effects of these responses (e.g., on coral cover) is difficult.

To address the challenges of capturing the impacts of multiple stressors across the GBR, several studies have taken a modeling approach whereby coral loss and recovery are integrated into statistical and/or simulation models of coral cover change (reviewed in Bozec and Mumby 2020, see also Vercelloni et al. 2017, Condie et al. 2018, Lam et al. 2018, Mellin et al. 2019). In these studies, coral population dynamics have been modeled as temporal changes in coral cover, most likely because this is the primary variable that is surveyed in monitoring programs. Although coral cover is a common metric of reef health, it does not resolve the demographic structure of corals, i.e. the relative composition of different stages or sizes. This is an important caveat because demographic changes are not necessarily reflected in changes in coral cover (Done 1995), so that impacts on a critical process (e.g. recruitment failure) may not be represented explicitly. Failure to identify which mechanisms (among partial or whole-colony mortality, recruitment or colony growth, Hughes and Tanner 2000) are implicated in coral cover change limits our ability to predict coral trajectories (Edmunds and Riegl 2020). Moreover, stress-induced coral mortality is often size-specific, and which size classes are affected will have important implications for the following rate of recovery. Accurate hindcast and forecast predictions of coral cover require a mechanistic approach by which the processes of coral gains (recruitment, colony growth) and losses (partial and whole-colony mortality) are considered explicitly at the colony level to account for size-specific and density-dependent responses.

We developed a mechanistic model of coral metapopulations to assess the cumulative impacts of recent multiple stressors that have affected the GBR. The model simulates the fate of individual coral colonies across > 3,800 individual reefs connected by larval dispersal while capturing some effects of water quality (suspended sediments and chlorophyll) on the early-life demographics of coral and CoTS. A reconstruction of recent (2008–2020) coral trajectories across the GBR was performed from (1) the integration of mechanistic data into empirical relationships that underlie the demography of corals and CoTS; (2) the calibration of stress-induced coral mortality and recovery with observations from the GBR; (3) the simulation of coral dynamics under spatially- and temporally-realistic regimes of larval connectivity, water quality, CoTS outbreaks, cyclones and mass coral bleaching; (4) the validation of these trajectories with independent coral cover observations. We then combined statistical and simulation-based approaches to evaluate the relative contribution in space and time of each driver to the reconstructed reef response. Specifically, we asked: (1) what are the individual and combined effects of acute stressors (cyclones, CoTS and bleaching) in terms of proportional coral loss across the GBR? (2) what is the relative importance of water quality and connectivity on recovery dynamics at both local and regional scales? (3) what is the reefs’ ability to sustain healthy levels of coral cover with their the current regime of acute and chronic disturbances and how does this vary in space? Finally, we develop a metric (reef *equilibrial cover*) that integrates the cumulative pressures operating on coral growth and stress-induced mortality to quantify reef resilience across the entire GBR. With this mechanistic evaluation of cumulative impacts and resilience we attempt to elucidate the main drivers of coral reef decline and provide guidance for reef monitoring and targeted management to help sustain a healthy GBR.

## MATERIAL AND METHODS

### Model general description

ReefMod (Mumby et al. 2007) is an agent-based model that simulates the settlement, growth, mortality of circular coral colonies and patches of algae over a horizontal grid lattice. With a six-month time-step, the model tracks the individual size (area in cm^2^) of coral colonies and algal patches affected by demographic processes, ecological interactions and acute disturbances (e.g., storms, bleaching) characteristic of a mid-depth (∼5–10 m) reef environment. The model has been successfully tested against *in situ* coral dynamics both in the Caribbean (Mumby et al. 2007, Bozec et al. 2015) and the Pacific (Ortiz et al. 2014, Bozec and Mumby 2019).

We developed the model further to integrate coral metapopulation dynamics across a spatially-explicit representation of the multiple reef environments of the GBR (ReefMod-GBR, Fig. 1A, Appendix S1). We refined a previous parameterization of coral demographics (Ortiz et al. 2014) with recent empirical data based on three groups of acroporids (arborescent, plating and corymbose) and three non-acroporid groups, including pocilloporids, encrusting and massive corals (Appendix S2: Table S1). The model was extended with explicit mechanisms driving the early-life dynamics of corals: fecundity, larval dispersal, density-dependent settlement, juvenile growth and background (chronic) mortality, mediated by water quality and transient coral rubble. In addition, a cohort model of CoTS was developed to simulate the impact of starfish outbreaks on coral populations. Processes of coral recovery and stress-induced mortality were calibrated with regional data, leading to a realistic modeling of the key processes driving coral populations on the GBR (Fig. 1B-C). ReefMod-GBR is implemented using the MATLAB programming language.

**Fig. 1.**
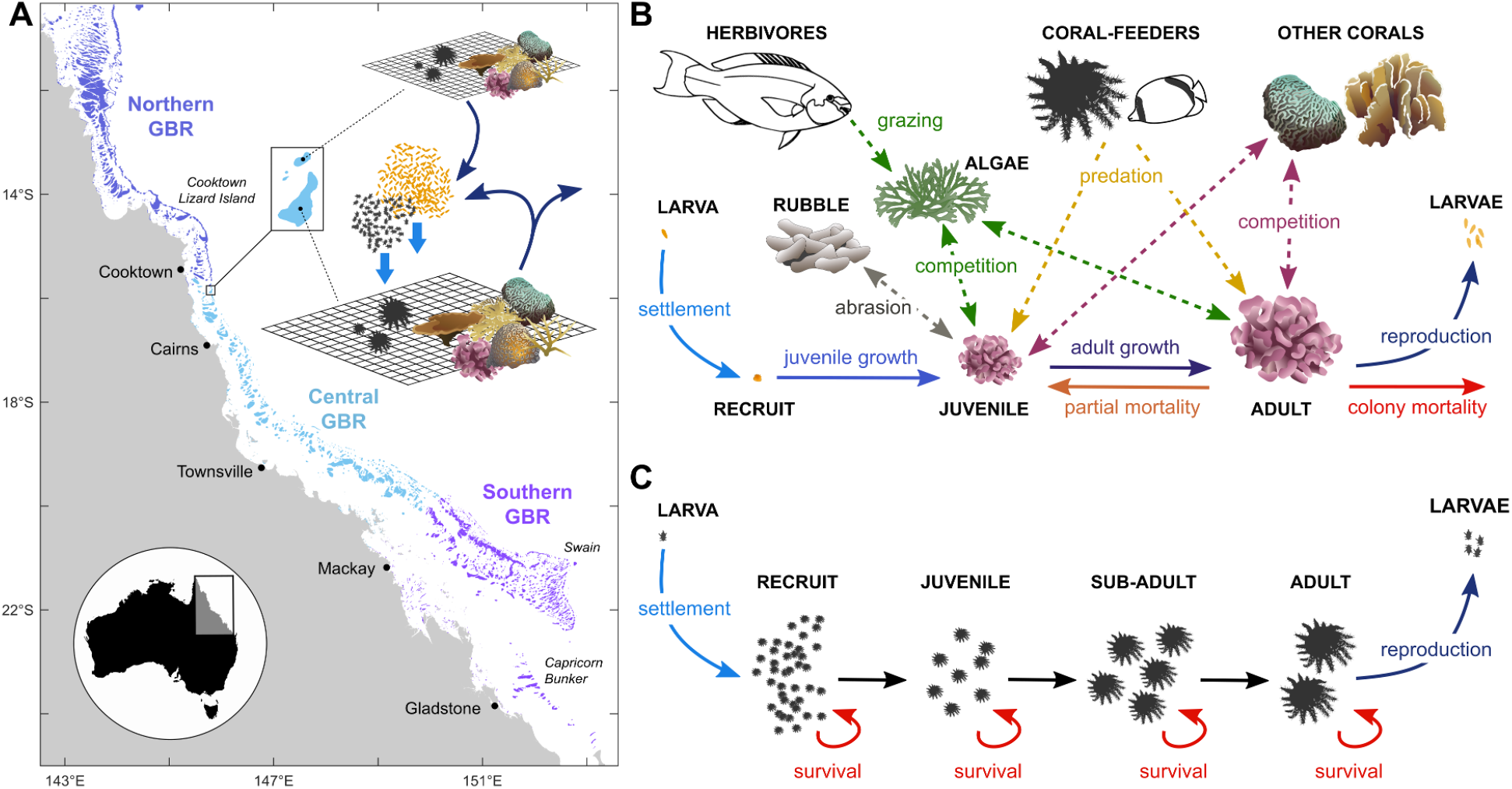
Schematic representation of the reef ecosystem model applied to the Great Barrier Reef (ReefMod-GBR). (**A**) Each of 3,806 individual reefs is represented by a 20 m × 20 m horizontal space virtually colonized by coral colonies belonging to six morphological groups. (**B**) Demographic processes (solid arrows) and ecological interactions (dashed arrows) affecting coral colonies individually. (**C**) Modeling of CoTS cohorts subject to size-specific survival during their life. For both corals and CoTS, settlement occurs from a pool of larvae that results from the retention of locally-produced offspring (self-supply) and the incoming of larvae from connected reef populations (external supply).

#### Model domain and spatial context

For simulating coral dynamics along ∼2,300 km length of the GBR, we used a discretization of the GBR consisting of 3,806 individual reef patches (Hock et al. 2017) across the northern, central and southern sections of the GBR Marine Park (Fig. 1A). A grid lattice of 20 × 20 cells, each representing 1 m^2^ of the reef substratum, was assigned to every reef patch (hereafter referred as a *reef*) identified by a convex polygon in the indicative reef (0–10 m) outline (GBRMPA 2007). Because larval dispersal and environmental forcing are not consistently available at intra-reef scales, each grid lattice represents a mean-field approximation of the ecological dynamics occurring within the environment of a defined reef polygon. This environment is characterized by historical events of tropical storm and heat stress, and a reconstructed regime of water quality during austral summer (wet season, from November to April) and winter (dry season, May to October). Within-reef variability of coral demographics is implicitly included through stochastic coral recruitment and mortality, but also temporally through probabilistic storm and heat stress events. Uncertainty in coral and CoTS trajectories is captured by running a minimum of 40 stochastic simulations. As a result, the model is spatially explicit in three ways: 1) by simulating individual coral colonies on a representative reefscape; 2) by linking coral demographics to their ambient stress regime; 3) by connecting reefs in a directed network that represents larval exchanges for both corals and CoTS.

#### Larval production and transport

Broadcast coral spawning on the GBR extends from October to December (Babcock et al. 1986). Following Hall and Hughes (1996), coral fecundity is a function of colony size and expressed as the total volume of reproductive outputs (Appendix S1) using species-specific parameters (Appendix S2: Table S1). Colony size at sexual maturity was fixed to 123–134 cm^2^ for the three acroporid groups and 31– 38 cm^2^ for the other groups, based on threshold sizes above which 100% of colonies were found reproductive (Hall and Hughes 1996). The number of offspring released by each coral group during the reproductive season is estimated by summing the total volume of reproductive outputs over all gravid colonies, assuming an average egg volume of 0.1 mm^3^ (*Acropora hyacinthus*, Hall and Hughes 1996).

The CoTS spawning period on the GBR extends from December to February (Babcock and Mundy 1992, Brodie et al. 2017). CoTS fecundity expressed as number of eggs is a function of wet weight (Kettle and Lucas 1987) derived from the representative mean size (diameter) of each age class of CoTS. The resulting fecundity-at-age prediction is multiplied by the density of the corresponding age class to calculate the total number of offspring produced on a grid lattice. Starfish become sexually mature when they are 2 years old (Lucas 1984).

During a spawning season, the number of coral and CoTS offspring produced on each grid lattice is multiplied by the area of the associated reef polygon to upscale reproductive outputs to the expected population sizes. Larval dispersal is then processed from source to sink reefs using transition probabilities (Hock et al. 2017, 2019) derived from particle tracking simulations generated by a three-dimensional hydrodynamic model of the GBR (Herzfeld et al. 2016). These probabilities of larval connectivity are combined with the number of larvae produced to estimate larval supply on every sink reef. Matrices of larval connectivity were determined for designated spawning times for both corals and CoTS over the 6 years for which the hydrodynamic models were available: wet seasons 2010-11, 2011-12, 2012-13, 2014-15, 2015-16, and 2016-17.

We note that local retention predicted by the connectivity matrices is extremely low for corals, as the relative proportion of coral larvae retained on a source reef is <0.01 for more than 95% of the 3,806 reefs. However, the empirical rates of larval retention for corals and CoTS across the GBR remain largely unknown. In a study of coral recruitment around a relatively isolated reef of the central GBR, Sammarco and Andrews (1989) observed that 70% of the coral spats collected within a 5 km radius were found within 300 m of the reef. Assuming that ∼40% of the produced larvae survive and become competent for settlement 8–10 days after spawning (Connolly and Baird 2010), a rate of 0.28 was considered as a minimum retention for both corals and CoTS and added to values predicted by dispersal simulations.

#### Larval supply and recruitment

For a given reef, the total number of incoming coral and CoTS larvae (i.e., from external supply and retention) is divided by the area of the reef to estimate a pool of larvae *L* (larva/m^2^) available for settlement. Assuming density-dependence in early (< 6 month) post-settlement survivorship, we first estimate a density potential for settlers (*D*_*settlers*_, settler/m^2^) as a Beverton-Holt (B-H) function (e.g., Haddon 2011) of the available larval pool (*L*):

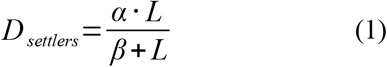

where *α* (settler/m^2^) is the maximum achievable density of settlers for a 100% free space and *β* (larva/ m^2^) is the stock of larvae required to produce half the maximum settlement. For CoTS, the actual density of 6-month-old recruits is obtained by reducing *D*_*settlers*_ to a 3% survived fraction due to intense predation (Keesing and Halford 1992, Okaji 1996). For corals, the actual number of 6-month-old recruits for each coral group is generated in each cell separately following a Poisson distribution with recruitment event rate λ (recruit/m^2^) calculated as:

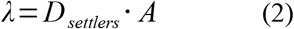

where *A* is the proportional space covered by cropped algal turf on a given cell, i.e., the substratum that is suitable for coral recruitment (Kuffner et al. 2006). This assumes that the probability of coral recruitment is directly proportional to available space (Connell 1997). Corals cannot recruit on sand patches, which are randomly distributed across the grid lattice at initialization (Appendix S1).

Recruitment parameters *α* and *β* were determined by calibration against GBR observations from offshore (mid- and outer-shelf) reefs. For corals (calibration for CoTS is presented thereafter), we simulated coral recovery on hypothetical reefs (see details in Appendix S1) and adjusted the two parameters with the double constraint of reproducing the recovery dynamics observed after extensive coral loss (Emslie et al. 2008, Fig. 2A) while generating realistic densities of coral juveniles (Trapon et al. 2013, Fig. 2B). Densities patterns of coral juveniles varied predictably along the recovery curve: first, by increasing as self-supply of larvae is enhanced by more abundant sexually-mature corals; second, by decreasing with the progressive reduction of settlement space. Recovery dynamics will likely vary with external supply, water quality and changes in coral community structure.

**Fig. 2.**
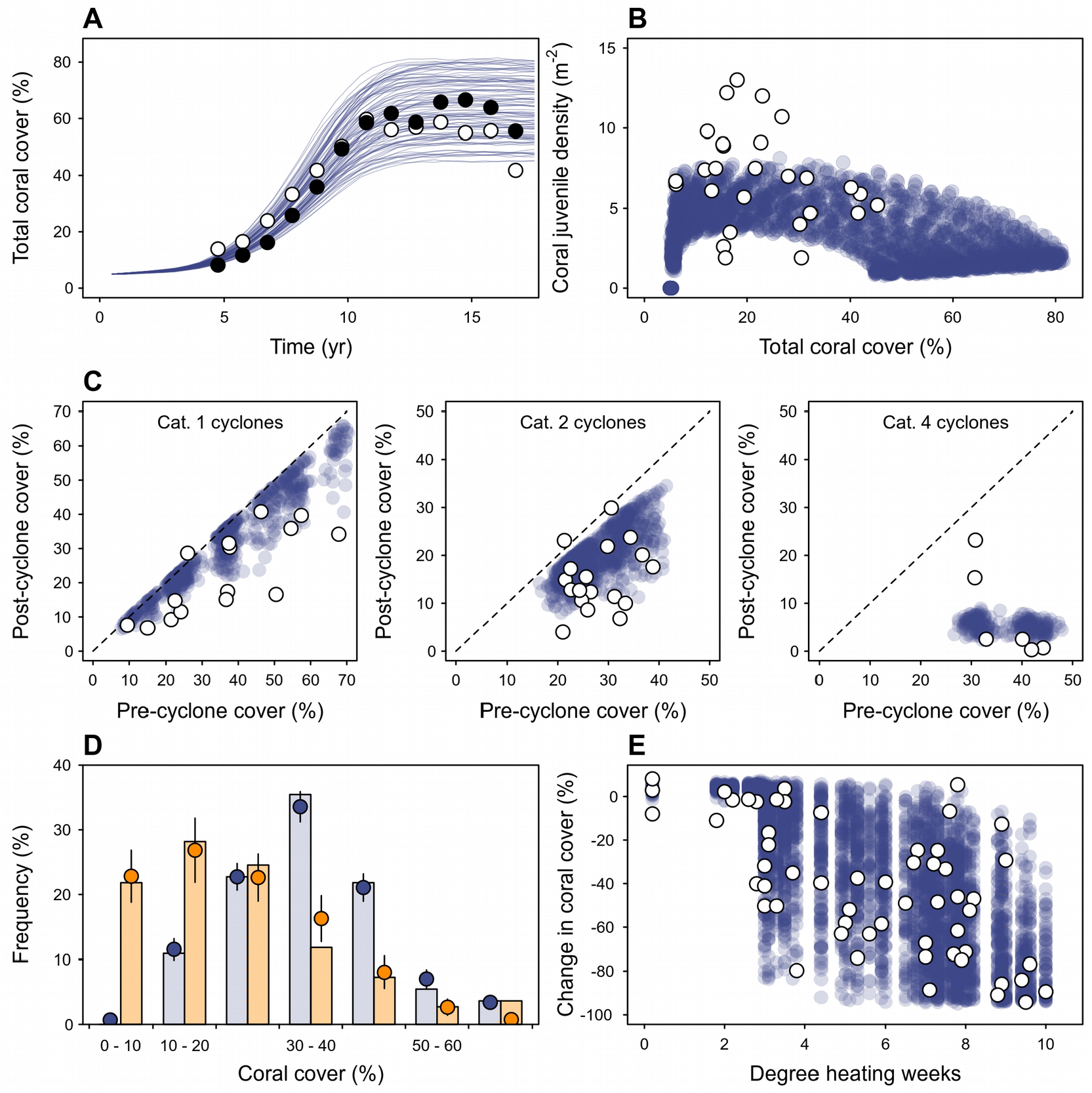
Calibration of ReefMod-GBR. (**A**) Mean coral recovery trajectories (overlaid lines) for hypothetical reefs (n = 40) after calibration of coral recruitment parameters with observed recovery on the outer-shelf of the northern (black dots) and southern (white dots) GBR (Emslie et al. 2008). (**B**) Resulting density of coral juveniles along the recovery trajectories compared with observations (Trapon et al. 2013) on the mid-shelf GBR (white dots). Juveniles defined here as corals < 5 cm excluding 6-mo old recruits (∼1 cm) for comparison. (**C**) Calibration of storm damages on AIMS LTMP sites (white dots: observations; blue dots: simulations, n = 40 stochastic runs) for the expected storm intensities (category 1, 2 and 4). Dotted lines indicate equality between pre- and post-disturbance coral cover (i.e., no change). (**D**) Frequency distributions of coral cover on 63 individual reefs before and after bleaching as measured (Hughes et al. 2018) across the GBR (blue and orange bars, respectively) and as simulated (blue and orange dots, respectively, n = 40 stochastic runs) after calibration of long-term bleaching mortality. (**E**) Corresponding changes in coral cover in response to heat stress (degree heating weeks, DHW) as observed (white dots, Hughes et al. 2018) and simulated (blue dots, n = 40 stochastic runs of the 63 reefs). A minimum 3 DHW was assumed for bleaching mortality to occur.

#### Early post-recruitment coral demographics

Six-month-old coral recruits have a fixed size of 1 cm^2^ and become juveniles at the next step if allowed to grow. Coral juveniles are defined by colony diameters below 4 cm. Their growth rate is set to 1 cm/y radial extension (Doropoulos et al. 2015, 2016) until they reach 13 cm^2^ (i.e., ∼4 cm diameter, 2 years old corals if no partial mortality has occurred) above which they acquire species-specific growth rate (Appendix S2: Table S1). With this parameterization, the maximum diameter of 3-year-old corymbose/small branching acroporids is 10.1 cm, which falls within the range of diameters (7.8– 13.7 cm) observed for *Acropora millepora* at this age (Baria et al. 2012).

Background whole-colony mortality of coral juveniles is set to 0.2 per year as recorded for *Acropora* spp. at Heron Island (Doropoulos et al. 2015). Corals above 13 cm^2^ have escaped the most severe post-settlement bottlenecks (Doropoulos et al. 2016) and are subject to group- and size-specific rates of partial and whole-colony mortality (Appendix S1, Appendix S2: Table S1).

#### Effects of suspended sediments on early coral demographics

River run-off expose coral reefs to loads of sediments that are transient in space and time (Schaffelke et al. 2012, Waterhouse et al. 2017). These dynamics were captured from retrospective (2010–18) spatial predictions of suspended sediments using the eReefs coupled hydrodynamic-biogeochemical model (Herzfeld et al. 2016, Baird et al. 2017). eReefs simulates the vertical mixing and horizontal transport of fine sediments across the entire GBR, including sediments entering the system through river catchments (Margvelashvili et al. 2018). We used the 4 km resolution model (GBR4) with the most recent catchment forcing (model configuration GBR4_H2p0_B3p1_Cq3b). Daily predictions of suspended sediment concentrations (*SSC*) were obtained by summing variables describing the transport and re-suspension of small-sized particles: *Mud* (mineral and carbonate, representative size 30 μm with a sinking rate of 17m with a sinking rate of 17 m/ d), which represent re-suspending particles from the deposited sediments, and *FineSed* (30 μm with a sinking rate of 17m, sinking rate 17 m/d) and *Dust* (1 μm with a sinking rate of 17m, sinking rate 1 m/d), which come from river catchments.

Suspended sediments influence many aspects of coral biology (Jones et al. 2015) but are only considered here at the early-life stages of broadcast spawning corals. Using published experimental data (Humanes et al. 2017a, 2017b), we modeled dose-response curves between *SSC* (mg/L) and the success rate of various early-life processes of corals: gamete fertilization, embryo development and subsequent larval settlement, recruit survival and juvenile growth (Appendix S3: Figs. S1A-C). Experiments and fitting procedures are detailed in Appendix S1.

Spawning corals release combined egg-sperm bundles that immediately ascend to the surface where fertilization and embryo development take place (Richmond 1997, Jones et al. 2015). To capture sediment exposure at these early (< 36 h) developmental stages, we extracted near-surface (−0.5 m) eReefs predictions of *SSC* at the assumed dates of mass coral spawning of six reproductive seasons (2011–2016). For each 4 km pixel, *SSC* was averaged over three days following field-established dates of *Acropora* spp. spawning (Hock et al. 2019) in the northern, central and southern GBR, then averaged among consecutive (split) spawning events (Appendix S3: Fig. S2). The resulting *SSC* values were assigned to the nearest reef polygon and used to predict, for each spawning season, the success of coral fertilization (Appendix S1: Eq. S10) and embryo development (manifested as subsequent larval settlement, Appendix S1: Eq. S11) which we combined to obtain an overall rate of reproduction success (Appendix S3: Figs. S1D, S3). The resulting rate can be multiplied by the number of coral offspring released before dispersal to simulate sediment-driven reductions in coral reproduction.

Daily predictions of *SSC* at 6-m depth from 2010 to 2018 (Appendix S3: Fig. S4) were used to predict the survivorship of *Acropora* recruits (Appendix S1: Eq. S12) and the growth potential of all juveniles (Appendix S1: Eq. S13). Recruit survivorship was expanded to a 6-month period by multiplying the daily survival rates over each summer (Appendix S3: Figs. S1E, S5). Juvenile growth potential was predicted from the *SSC* values averaged over each season (Appendix S3: Fig. S1F, S6, S7).

#### Impacts of cyclones on corals

Cyclone-generated waves cause coral dislodgement and fragmentation. While the wave power needed to dislodge colonies of various sizes and shapes has been estimated (Madin et al. 2014), a measure of wave power at the scale of individual colonies is often unavailable. Indeed, work is underway to estimate coral loss from the duration of local exposure to cyclone-generated sea states capable of damaging reefs, as this can more readily be reconstructed than wave power. In the meantime, we approximated storm-induced colony mortality as a function of colony size and storm intensity defined on the Saffir-Simpson scale (1–5) (Mumby et al. 2007, Edwards et al. 2011). Briefly, the probability of whole-colony mortality for the most severe storm (category 5) is assumed to be a quadratic function of colony size (Massel and Done 1993, Appendix S1): small colonies avoid dislodgement due to their low drag, intermediate-sized corals have greater drag and are light enough to be dislodged, whereas large colonies are heavy enough to prevent dislodgement. A Gaussian-distributed noise *ε* ∼ *N*(*μ* = 0, *σ* = 0.1) adds variability to mortality predictions. For storm categories 1–4, these predictions are lowered by 95%, 88%, 75% and 43%, respectively (Edwards et al. 2011, Appendix S1). Coral colonies larger than 250 cm^2^ suffer partial mortality (i.e., fragmentation): the proportional area lost by a colony follows a normal distribution *N*(*μ* = 0.3, *σ* = 0.2) for a category 5 storm (Mumby et al. 2007), while the aforementioned adjustments are applied for other storm category impacts. Finally, scouring by sand during a cyclone causes 80% colony mortality in recruit and juvenile corals (Mumby 1999).

Because the above parameterization was initially derived for Caribbean reefs (Mumby et al. 2007, 2014, Edwards et al. 2011, Bozec et al. 2015), cyclone-driven mortalities were calibrated with GBR observations of storm damages using the benthic survey database of the Australian Institute of Marine Science (AIMS) Long-Term Monitoring Program (LTMP). We extracted coral cover data on reefs surveyed within one year of storm damages and estimated for each reef the expected cyclone intensity using the Database of Past Tropical Cyclone Tracks of the Australian Bureau of Meteorology (BoM) (see details in Appendix S1). The magnitude of partial- and whole-colony mortality was tuned until a reasonable match between the simulated and observed coral cover changes was found for the expected cyclone categories (Fig. 2C).

#### Mass coral bleaching

Widespread coral bleaching is assumed to be driven by thermal stress (Berkelmans 2002, Hughes et al. 2017, 2018). We used the Degree Heating Week (DHW) as a metric of the accumulated heat stress to predict bleaching-induced coral mortality (Eakin et al. 2010, Heron et al. 2016). In an extensive survey of shallow (2 m depth) corals across the GBR during the 2016 marine heatwave, Hughes et al. (2018) recorded initial coral mortality (i.e., at the peak of the bleaching event) on reefs exposed to satellite-derived DHW (Liu et al. 2017). A simple linear regression model (*R*^2^ = 0.49, *n* = 61) can be fit to the observed per capita rate of initial mortality, *M*_*BleachInit*_ (%), as a function of local thermal stress (Appendix S3: Fig. S8):

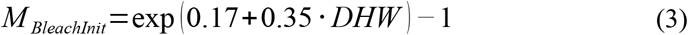

*M*_*BleachInit*_ was used as the incidence rate for both partial and whole-colony mortality caused by bleaching, assuming they are correlated in their response to thermal stress. The resulting mortality incidences were further adjusted to each coral group (Appendix S2: Table S1) following reported species susceptibilities (Hughes et al. 2018). For a coral affected by partial mortality due to bleaching, the extent of tissue lost (Baird and Marshall 2002) was set to 40% of the colony area for small massive/submassive (observations on *Platygyra daedalea*), 20% for large massive corals (*Porites lobota*), and a minimal 5% for the three acroporid groups (*A. hyacinthus* and *A. millepora*) extended to pocilloporids due to morphological similarities.

Because Eq. 3 only captured initial mortality of the 2016 GBR heatwave, coral response over an entire bleaching event (i.e., including post-bleaching mortality) was determined by calibration with coral cover changes reported in the following 8 months (Hughes et al. 2018). We initialized hypothetical reefs with the observed pre-bleaching values of coral cover (Fig. 2D) and simulated heat stress using the DHW values recorded in 2016 (Appendix S1). The overall magnitude of the resulting bleaching mortalities (i.e., *M*_*BleachInit*_) was progressively increased until the predicted coral cover changes matched the observations (Fig. 2D, E).

#### Crown-of-thorns starfish outbreak dynamics

Outbreak dynamics of the crown-of-thorns starfish (*Acanthaster* spp, CoTS) were simulated using a simple cohort model where starfish density is structured in 6-month age classes. The model integrates nutrient-limited larval survivorship and age-specific mortality which are key for predicting outbreak dynamics (Birkeland and Lucas 1990, Pratchett et al. 2014).

Because the survival of pelagic-feeding CoTS larvae is strongly dependent on phytoplankton availability (Okaji 1996, Wolfe et al. 2017), high nutrients following terrestrial run-off, especially after intense river flood events, may have the potential to trigger population outbreaks (Brodie et al. 2005, Fabricius et al. 2010). A daily survival rate (*SURV*) of CoTS larvae can be estimated from the concentration of chlorophyll *a* (Chl *a*, μm with a sinking rate of 17g/L), a proxy of phytoplankton abundance (Fabricius et al. 2010, Appendix S3: Fig. S9):

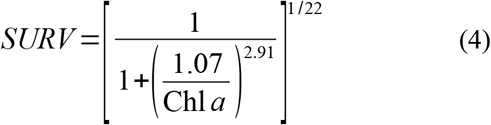

We extracted subsurface (0–3 m) daily concentrations of total chlorophyll *a* predicted by eReefs during eight consecutive spawning seasons (Dec. 2010–Feb. 2018, Appendix S3: Fig. S10). For each 4 km pixel, the average daily survival (geometric mean) over a spawning season was extended to 22 days (duration of the developmental period, Fabricius et al. 2010) and assigned to the nearest reef polygon (Fig. 3A, Appendix S3: Fig. S11). Nutrient-enhanced larval survivorship on a reef was simulated by multiplying the predicted survival to the number of offspring released before dispersal.

**Fig. 3.**
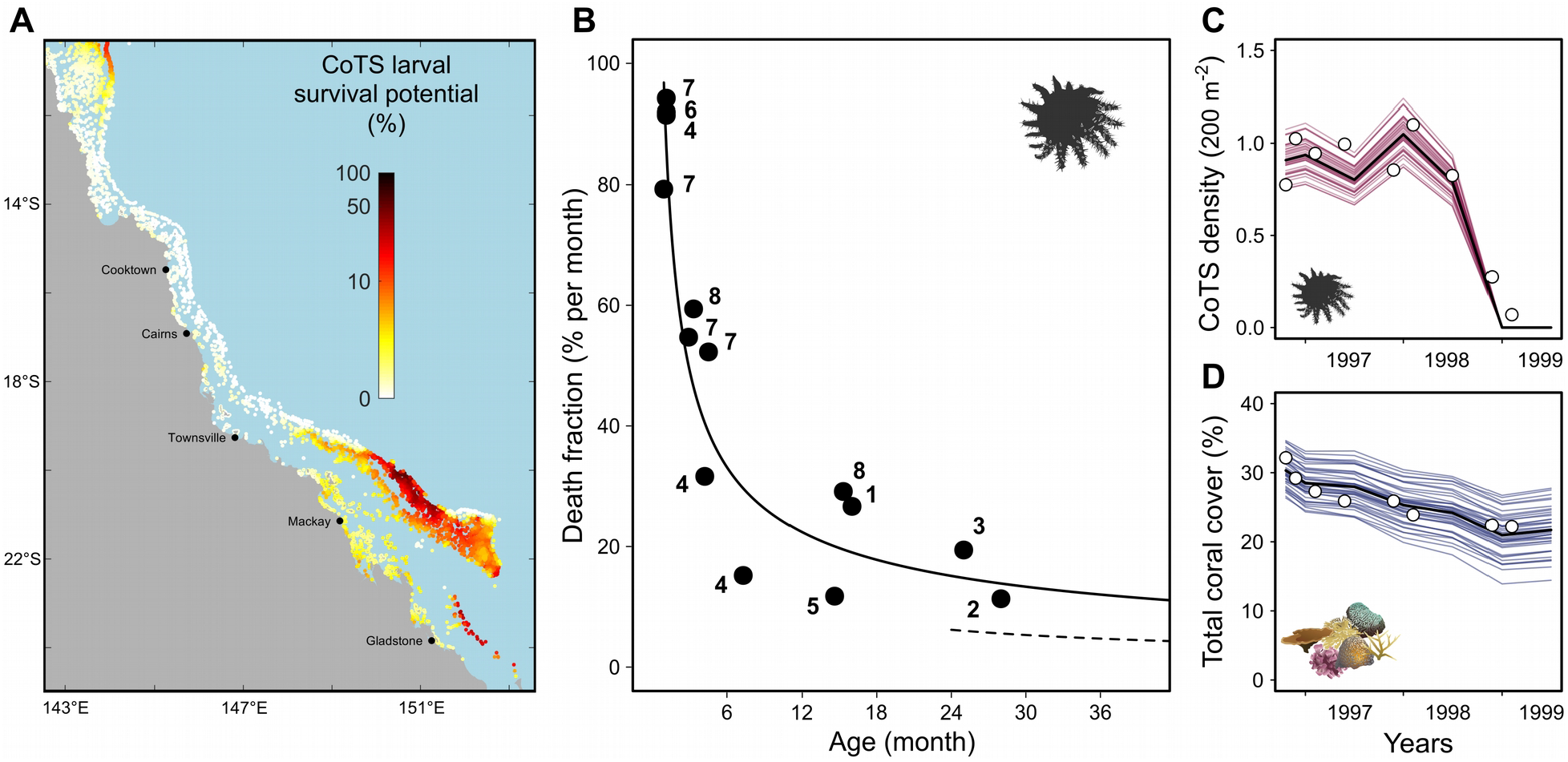
(**A**) Percent survival rate of CoTS larvae before dispersal derived from subsurface (0–3 m) daily predictions (eReefs-GBR4) of Chl a during the spawning season (Dec.–Feb.) averaged over the period 2010–2018. (**B**) Point estimates of CoTS mortality (monthly % death fraction) as a function of individual age derived from manipulative experiments and cohort surveys (Appendix S1: Table S2) with the fitted log-log linear model (loge y = 4.51 – 0.57 · loge x) equivalent to Eq. 5. Age was estimated as the median age of the cohort during the study period. Temporal changes in CoTS densities (**C**) and coral cover (**D**) as observed at Lizard Island (white dots, Pratchett 2005, 2010) and as simulated (colored lines: replicate trajectories; black lines: average trajectories) after calibration of mortality of 2 yr+ starfish (dotted line in B), coral consumption and recruitment parameter β. Temporal changes in starfish size distribution (Fig. S12) were also included in the calibration.

After dispersal, larval supply to a given reef was converted into a number of settlers (Eq. 1) with parameters determined by calibration (detailed below). The fate of newly settled CoTS was determined by age-specific rates of mortality sourced from the literature (Appendix S2: Table S2). To derive this mortality function, we first estimated daily mortality rates from the reported surviving fraction of CoTS individuals and the period of observations. A log-log linear model (*R*^2^ = 0.80, *n* = 8) was then fitted to the resulting mortality-at-age estimates (Fig. 3B):

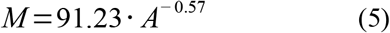

where *M* represents the monthly mortality rate (%) of CoTS at age *A* (month). In simulations, mortality-at-age was converted to a 6-month equivalent (1 – (1 – *M* / 100)^6^) and applied to the corresponding age class at every step. The same mortality function was used for all reefs in the absence of reliable data on predation on CoTS. Maximum CoTS age was set to 8 years (Pratchett et al. 2014) with 100% of individuals older than that dying due to senescence.

The amount of coral surface consumed by CoTS over a 6-month period was determined from published rates of consumption per individual size (starfish diameter) during summer and winter (Keesing and Lucas 1992) after representative size-at-age conversions (Engelhardt et al. 1999). As a result, CoTS substantially feed on corals from the age of 18 mo+ (∼150–200 mm diameter). The amount of coral surface consumed for each coral group was determined using empirically-derived feeding preferences (De’ath and Moran 1998). While relative feeding proportions reflect a strong preference for the three *Acropora* groups (∼75% of CoTS consumption), these are further adjusted to the proportion of each coral group currently available on a reef.

The density of coral-eating CoTS (18 mo+) collapses due to starvation when the cover of all acroporids and pocilloporids drops below 5%. Although this allows reproducing the observed rapid decline of outbreaking CoTS when coral is depleted (Moran 1986), mass mortalities in high-density populations of *Acanthaster* can also be triggered by disease (Zann et al. 1987, 1990, Pratchett 1999) before significant coral damage occurs (Pratchett 2010). To capture this density-dependent process, an outbreaking CoTS population will collapse after a random time period drawn from a uniform distribution of 2–5 years, which is the duration of most observed outbreaks (Moran 1986, Pratchett et al. 2014). A CoTS population is considered outbreaking when the density of 18 mo+ starfish reaches 0.6 individuals/400 m^2^ (Moran and De’ath 1992).

CoTS outbreak dynamics and associated impacts on corals were calibrated using observations from Lizard Island, northern GBR (Pratchett 2005, 2010). Starfish populations were initialized with the density-at-size recorded in Oct-Dec 1996 after appropriate size-age conversion (Engelhardt et al. 1999). Because the first observed starfish size class (diameter < 15 cm) is likely underestimated by visual surveys (MacNeil et al. 2016), its density was deduced from the 15–20 cm class following mortality at the corresponding age. Recruit (0–6 month old starfish) density was set to zero as expected in winter. Here, CoTS populations were forced to collapse after 2 years as observed (Pratchett 1999, 2005). Simulations reproduced the observed changes in CoTS density (Fig. 3C) and size distribution (Appendix S3: Fig. S12) after lowering the mortality of 2 y+ (> 20 cm) starfish (Fig. 3B). Maximum settlement rate (α) was fixed) was fixed to 100 settlers/m^2^, which, with the above adjustment of adult mortality, gives an adult population size of ∼64 adults (> 25 cm) per 400 m^2^ reef area, similar to the maximum adult densities observed on the GBR (Engelhardt et al. 1999, 2001). The steepness of the B-H relationship (*β*) was set to 12,500 larva/m^2^. Starting with the Oct-Dec 1996 average coral cover (*μ* = 30.7%, *σ* = 0.2×*μ*, half being acroporids, Pratchett 2010), reproduction of the observed coral cover changes (Fig. 3D) required a near-doubling (× 1.8) of the published feeding rates.

#### Unconsolidated coral rubble

Coral mortality following acute stress generates loose coral debris that cover the reef substratum and inhibit coral recruitment (Fox et al. 2003, Biggs 2013). As a first approximation, we assume that the percent coral cover lost after disturbance converts into percent rubble cover, although collapsed coral branches might cover a larger area than their standing counterparts. Structural collapse occurs immediately after cyclones but is delayed for three years after bleaching and CoTS predation (Sano et al. 1987). Coral juveniles do not survive on unconsolidated rubble (Fox et al. 2003, Viehman et al. 2018), which amounts to reducing their survivorship by the proportion of the reef area covered by rubble. Loose coral rubble tend to stabilize over time with processes of carbonate binding and cementation (Rasser and Riegl 2002). These dynamics were approximated using an exponential decay function (Appendix S3: Fig. S13) assuming that ∼2/3 of coral rubble is consolidated after 4 years (Biggs 2013).

#### Macroalgae and grazing

The modeling of grazing and algal dynamics is detailed elsewhere (Bozec et al. 2019) so is only briefly described here. The model simulates algal dynamics by 1-month iterations using empirical rates of macroalgal recruitment and growth. Each grid cell can be occupied by four algal groups: (1) closely cropped algal turf (< 5mm), (2) uncropped algal turf (> 5mm), (3) encrusting fleshy macroalgae and (4) upright macroalgae. Cropped algal turf is the default substrate maintained by repeated grazing onto which corals can settle and grow. When a cell is left ungrazed for 1 month, diminutive algal turf becomes uncropped and the two macroalgal groups grow following a logistic curve (Bozec et al. 2019). Due to limited spatial data on fish and algae, macroalgae and turf were assumed to be maintained in a cropped state suitable for coral settlement. Realistic spatial predictions of grazing levels is yet to be developed for the GBR and will require extensive data on the size structure and species composition of herbivorous fish across a range of habitats (Mumby 2006, Fox and Bellwood 2007).

### Reconstruction of recent (2008–2020) reef trajectories of the GBR

Model simulations were run with spatially- and temporally-realistic regimes of water quality (SSC and Chl *a*), storms and thermal stress to reconstruct the trajectory of coral cover of the 3,806 reefs between 2008–2020 (end of winter 2007 to end of winter 2020).

Initial coral cover on each reef was generated at random from a normal distribution *N*(*μ, σ* = 0.2×*μ*) with mean value *μ* derived from AIMS monitoring surveys (Sweatman et al. 2008, Thompson et al. 2019) performed on 204 reefs between 2006–2008 (Appendix S1). Reefs that were not surveyed during this period were initialized with the mean coral cover of the corresponding latitudinal sector (11 sectors, Sweatman et al. 2008) and shelf position (inshore, mid-shelf and outer shelf). Initial cover was generated for each coral group separately following the average community composition of each sector and shelf position. Random covers of loose coral rubble and sand were generated with a mean of 10% and 30%, respectively.

The 2010–2018 regime of water quality (i.e., suspended sediments and Chl *a*) predicted by eReefs was imposed as a recursive sequence over the 2008–2020 period. The same sequence was applied to the selection of connectivity matrices to preserve spatial congruence between larval dispersal and the hydrodynamic forcing of water quality. Past exposure to cyclones was derived from sea-state predictions of wave height (Puotinen et al. 2016). The potential for coral-damaging sea state (wave height > 4 m) was determined using a map of wind speed every hour within 4 km pixels over the GBR for cyclones between 2008–2020. Any reef containing a combination of wind speed and duration capable of generating 4 m waves, assuming sufficient fetch, was scored as positive for potential coral-damaging sea-state in the respective year. Where damaging waves were predicted, an estimate of cyclone category was deduced from the distance to the cyclone track extracted from the BoM historical database. To simulate past exposure to thermal stress, we extracted from the NOAA Coral Reef Watch (CRW) Product Suite version 3.1 (Liu et al. 2017) the 2008–2020 annual maximum DHW available at 5-km resolution, consistent with the DHW-mortality relationship of the 2016 bleaching (Eq. 3, Hughes et al. 2018). Reefs were assigned the maximum DHW value of the nearest 5-km pixel.

Exposure to *Acanthaster* outbreaks was hindcast by combining starfish demographic simulations with observed abundance from monitoring (*n* = 289 reefs with at least one survey between 2008–2020) conducted by the AIMS LTMP (Sweatman et al. 2008) and the Great Barrier Reef Marine Park Authority (GBRMPA) Reef Joint Field Management Program (GBRMPA 2019). Initial CoTS densities were predicted by hindcast (1985–2008) simulations of the Coral Community Network (CoCoNet) model (Condie et al. 2018). This predator-prey model simulated age-structured CoTS populations with fast- and slow-growing coral cover dynamics across ∼3,000 reefs using a representative regime of storms and bleaching (*n* = 50 stochastic runs). Mean densities of adult CoTS (as mean counts per hypothetical manta tow) predicted in 2008 were assigned to the 3,806 reefs and treated as rate parameter values of a Poisson distribution in order to initialize ReefMod with random CoTS densities. At the following steps, CoTS populations on reefs that were not surveyed in the respective year were predicted by population dynamics, whereas reefs surveyed that year were imposed the corresponding observation of adult count. Assuming 0.22 CoTS per tow represents 1,500 CoTS/km^2^ (Moran and De’ath 1992), input count values were transformed into an equivalent starfish density per reef area and dis-aggregated by age following age-specific predictions of starfish mortality. Density-at-age was further corrected for imperfect detectability using empirical predictions from MacNeil et al. (2016).

To evaluate model performance, predictions of total coral cover over time were compared to observed time-series from AIMS monitoring (transects and standardized manta tows, Appendix S1). We selected *n* = 67 individual reefs monitored at least 12 times between 2009–2020 (i.e., excluding 2008 surveys used for model initialization) and calculated for each survey the difference between the observed total coral cover and the mean prediction (*n* = 40 simulations) for the corresponding season. The resulting deviations were averaged over each time-series to assess prediction errors in the different sections of the GBR.

### Assessment of cumulative impacts and resilience during 2008–2020

To investigate temporal coral changes across the GBR, we quantified year-on-year absolute changes (*AC*) in percent coral cover for each reef:

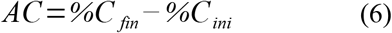

where *%C*_*ini*_ and *%C*_*fin*_ are the percentage total coral cover at the beginning and at the end of a one-year period, respectively. Because the magnitude of coral cover change is likely dependent on the initial value of coral cover (Côté et al. 2005, Graham et al. 2011), we also calculated for every reef and every year the relative rate of coral cover change (*RC*) as follows:

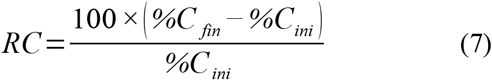

Within one time-step of ReefMod simulation (i.e., 6 months), stress-induced coral mortality (i.e., due to CoTS, cyclones, and bleaching) is applied after the processing of coral recruitment, growth and natural mortality. To quantify the individual impact of each of these stressors, their associated loss of total coral cover was tracked annually and expressed both as absolute (% cover/y) and relative (i.e., proportional to coral cover *before* disturbance, %/y). The latter metric allowed calculation of standardized annual rates of coral mortality (*m*_*r,s*_) on every reef *r* due to each stressor *s* (i.e., CoTS, cyclones and bleaching), a necessary step for assessing the relative importance of the three acute stressors across the entire GBR.

To assess the potential of coral recovery, the absolute change in total coral cover over 6 months was extracted for each reef *before* stress-induced coral mortality, thus providing an estimate of total coral cover growth in the absence of disturbances. Spatial and temporal variations of these rates of coral community growth (*g*, in % cover per 6 month) were analyzed with generalized linear models (GLM). Simulated data of the first two time-steps were excluded to reduce the influence of model initialization. Using GLMs as tools of variance partitioning for simulated data sets (White et al. 2014), we estimated the variance components of *g* per reef (*n* = 3,806) × time-step (*n* = 24) × run (*n* = 40) explained by eight environmental variables: total coral cover before growth; cover of sand patches; cover of loose coral rubble; water quality-driven percentage success of (i) coral reproduction, (ii) recruit survival of acroporids and (iii) juvenile growth; relative proportion of external vs internal (self) larval supply in the connectivity matrices, where external supply is the sum of the connection strengths from source reefs; number of connections from source reefs. The cover of coral rubble was time-averaged for each reef × run because its fluctuations and associated effects on coral juveniles are unlikely to impact coral cover over 6-month. For the same reason, the reef-specific values of water quality and connectivity variables were averaged over time. Residuals were modeled with a gamma distribution with a log link function. Because *g* can be negative (i.e., when natural mortality exceeds recruitment and colony growth) with a minimal value of –1.5% cover per 6 month, it was fitted as *g* + 2 to obtain a strictly positive response variable.

The GLM predictions of *g* for a given reef environment can be used to simulate a stepwise process of coral cover growth using a simple recursive equation:

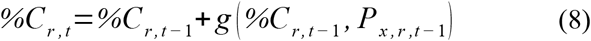

where the incremental growth of total coral cover (*g*) on reef *r* at step *t* is predicted from the previous-step value of coral cover (*%C*_*r, t-1*_) and the other environmental predictors (*P*_*x, r, t-1*_). To assess the influence of water quality on coral recovery on inshore reefs, we simulated coral growth curves from an initial 5% cover using Eq. 8 and the percentage success of early-life coral demographics calculated from representative steady-state (i.e., time-averaged) *SSC* values. The other predictors (sand, coral rubble and connectivity drivers) were set to their median value. Finally, to visualize the recovery potential across the entire GBR, we mapped the standardized annual growth rate of every reef obtained by simulating Eq. 8 over two time-steps (i.e., yearly), from a hypothetical 10% coral cover and with the reef-specific values of water-quality and connectivity predictors.

Assessing the cumulative impacts of multiple stressors requires integrating both their acute and chronic effects on coral mortality and growth. This was performed by simulating coral cover in every reef as a dynamic balance between cover growth *g* and the combined rates of annual mortality *m*_*r, s*_ due to CoTS, cyclones and bleaching:

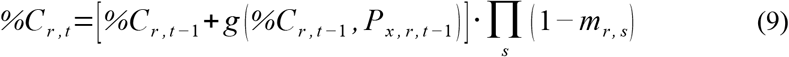

With this formulation, coral cover on a given reef has a single stable equilibrium (i.e., independent of initial cover) which is fully determined by the adverse effects of growth and stress-induced mortality. This equilibrial state approximates the value of coral cover that would be obtained when averaged over a long period of time, provided that the regimes of recovery and disturbance remain unchanged.

The equilibrial cover of each reef was determined based on the associated forcing of water quality, larval connectivity, cyclones, bleaching and CoTS. Although the 2010–2018 fluctuations of *SSC* can be considered as a near-typical regime of water quality, episodic storms and marine heatwaves experienced between 2008–2020 may not adequately represent average exposures. We thus gathered additional data to extend the cyclone and bleaching regimes and calculate more reliable annual mortalities. For cyclones, we used simulated regimes of region-scale occurrence of storm categories that combine GBR historical statistics (1970–2011) with synthetic cyclone tracks (Wolff et al. 2018). For bleaching, we extended the NOAA time series of annual maximum DHW back to 1998 to capture earlier (i.e., 1998, 2002) mass bleaching on the GBR (Berkelmans et al. 2004, Hughes et al. 2017). From these historical rates of disturbances, we generated 100 stochastic scenarios of storm and bleaching events over 20 years for every reef and inferred the associated mortality (relative coral cover loss) from regression models derived from the 2008-2020 reconstruction (Appendix S3: Fig. S14). The predicted coral losses were averaged across all scenarios to generate mean annual mortalities for each reef. For CoTS, we used the mean annual mortalities of the 2008-2020 reconstruction. Eq. 9 was simulated until a near-equilibrium cover was achieved for each reef, and the resulting equilibrial states used as a metric quantifying the ecosystem potential of reefs under their cumulative stress regime of cyclones, bleaching, CoTS and water quality. This metric is a critical asset for the evaluation of engineering resilience (Holling 1996) and allows setting reference values against which ecosystem performance can be measured (Mumby and Anthony 2015, Lam et al. 2020).

## RESULTS

### Reconstructed 2008-2020 reef trajectories

Hindcast simulations of 3,806 reefs (Fig. 4A) indicated an overall decline of corals during the period 2008–2020 with a global mean coral cover that dropped from ∼29% to ∼19% (annual absolute cover loss –0.74 % cover/y over 13 years). This is equivalent to a 33% relative loss of the initial cover. There was considerable variation among the three regions in the annual rate of coral cover change (Table 1) due to geographic differences in the timing and magnitude of coral mortality events and recovery periods. Overall, corals in the northern, central and southern regions declined by –15.2, –2.9 and –8.6 % cover, respectively. This corresponds to a relative loss of the initial cover of 54%, 13% and 26% in each respective region. Cross-shelf variability in reef trajectories was important (Appendix S3: Fig. S15) with the strongest relative losses obtained for the inner-shelf (63-73%), the northern mid-shelf (58%) and southern outer-shelf (44%) regions (Appendix S2: Table S3).

**Table 1.** Mean annual rates (% cover/y) of absolute coral cover change (*AC*), growth and mortality from disturbances (CoTS, cyclones and bleaching) for 2008–2020. Growth represents the net outcome between coral cover growth (due to recruitment and colony extension) and natural mortality, in the virtual absence of disturbances (formally, before disturbances occur).

|  | GBR | North | Central | South |
| --- | --- | --- | --- | --- |
| Net annual cover change | –0.7 | –1.2 | –0.2 | –0.7 |
| Annual cover loss |  |  |  |  |
| due to CoTS | –0.4 | –0.4 | –0.4 | –0.5 |
| due to cyclones | –1.9 | –0.6 | –2.3 | –2.9 |
| due to bleaching | –2.5 | –4.0 | –1.7 | –1.6 |
| total | –4.9 | –5.0 | –4.5 | –5.0 |
| Annual growth | +4.1 | +3.8 | +4.2 | +4.3 |

**Fig. 4.**
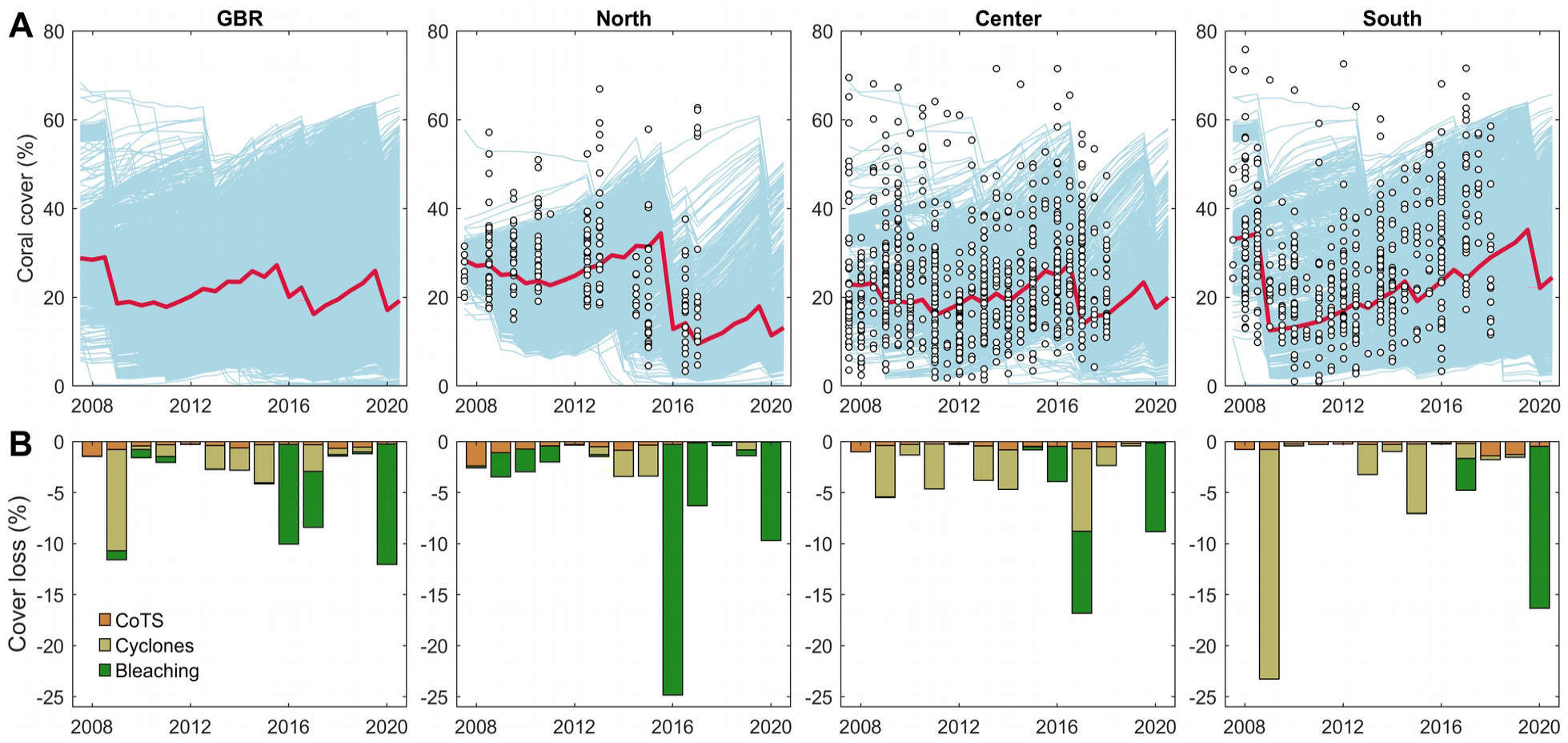
(**A**) Hindcast (2008–2020) reconstruction of coral cover trajectories (blue lines: individual reef trajectories averaged over 40 simulations; red line: regional average weighted by the log-transformed area of reef polygons) for the whole GBR (n = 3,806 reefs) and the northern (n = 1,201 reefs), central (n = 957 reefs) and southern (n = 1,648 reefs) regions. Data points indicate observations of coral coverage from AIMS monitoring (transect and transformed manta tow estimates, Appendix S1). (**B**) Mean annual absolute loss of coral cover due to CoTS, storm damages and heat stress during 2008–2020.

At the reef level, the reconstructed coral trajectories generally matched field observations from monitoring data (Fig. 5) including the magnitude of observed coral declines following acute disturbances and the post-disturbance timing of coral recovery. Among the 67 monitored reefs selected to validate model predictions (Appendix S3: Fig. S16), 75% exhibited a mean deviation (predicted – observed averaged over the time series) between –8.1 and +5.8 % coral cover. The most frequent model errors were due to inaccuracies in the predicted occurrence (false positives and negatives) or intensity of storm damages.

**Fig. 5.**
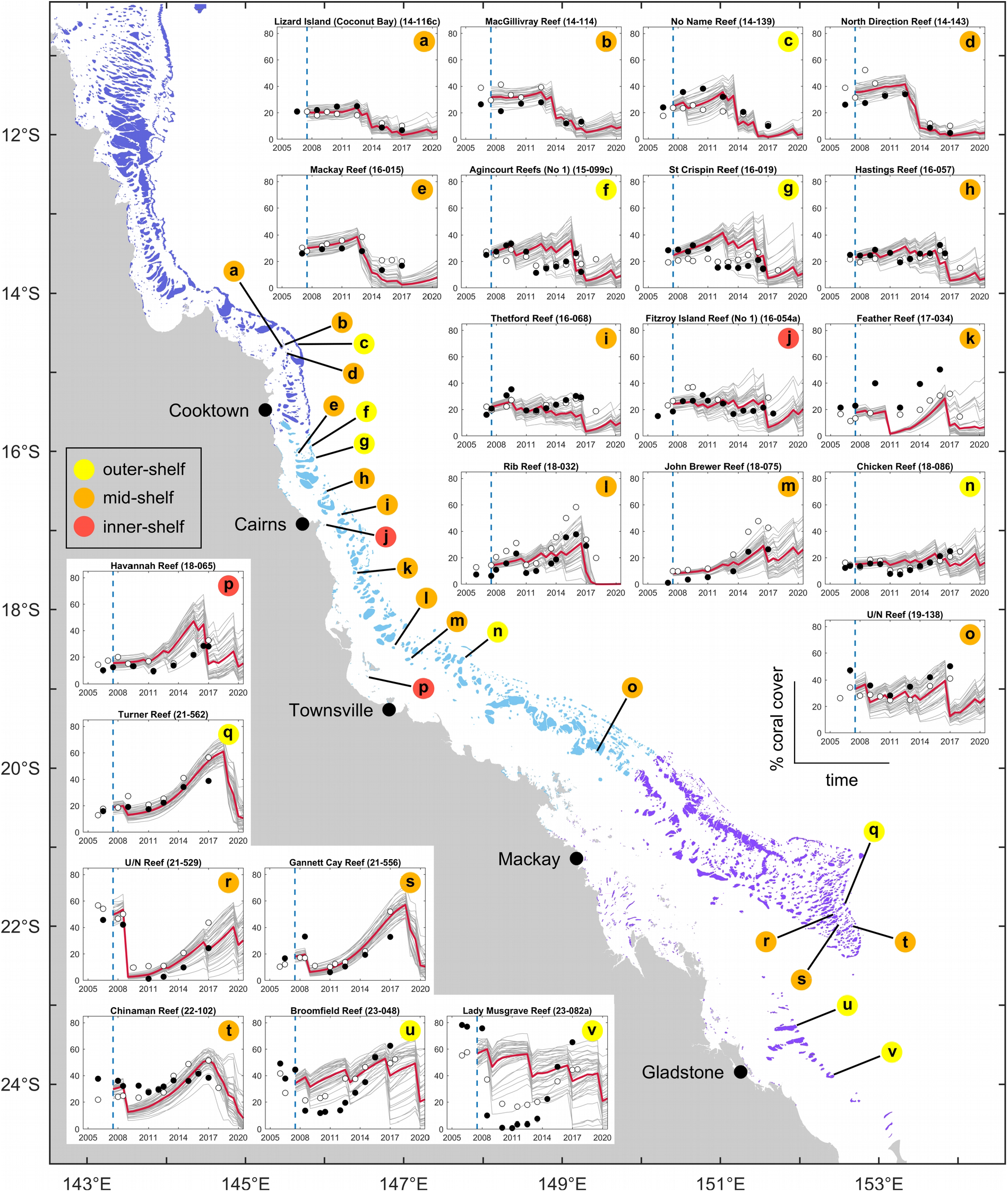
Validation of the reconstructed trajectories of coral cover with field observations from the AIMS LTMP (filled circles: point-intercept transects; open circles: standardized manta tows, Appendix S1). The dashed blue line indicates model initialization (winter 2007), whereby initial coral cover was determined as the mean cover of surveys performed between 2006 and 2008. Reefs selected for validation (n = 22) gathered at least 14 surveys during 2009–2020 (see Appendix S3: Fig. S16 for a broader selection of surveyed reefs).

### Coral loss due to bleaching, cyclones and CoTS

There were considerable variations in the magnitude of coral loss across years and among the three regions (Fig. 4B). Averaged over the 2008–2020 period and across the entire GBR (Table 1), bleaching was the most important driver of coral loss (–2.5 % cover/y mean annual absolute cover loss) followed by cyclones (–1.9 % cover/y), well ahead of CoTS (–0.4 % cover/y). The three stressors resulted in a cumulative annual loss of –4.9 % cover/y throughout the GBR, with the northern and central regions being the most and least affected, respectively.

Impacts of bleaching essentially occurred during the last five years, with intense and widespread heat stress (Fig. 6A) causing an estimated mean absolute decline of –9.8 % cover in 2016, –5.5 % cover in 2017 and –11.8 % cover in 2020 throughout the entire GBR (Table 2, Fig. 6B). The 2020 heatwave produced the most severe impacts in terms of proportional coral loss (40% mean loss of pre-bleaching coral cover, Table 2) and number of impacted reefs (85% of reefs with a proportional loss > 20%; 2016: 39%; 2017: 45%, Fig. 6C). The Northern GBR was the most severely impacted sector with all three bleaching events causing significant coral loss, especially during 2016 (mean absolute loss of – 24.6 % cover). The central region was also affected by the three heatwaves, experiencing increasing levels of coral mortality at each bleaching event. While escaping mass bleaching in 2016, the Southern GBR was hit by the two following heatwaves, especially in 2020 (–15.9 % cover). Overall, only 10% of the GBR experienced less than 20% proportional loss for all three events of mass bleaching. Spatial discrepancies between the footprint of heat stress and absolute cover loss (e.g., in the far north in 2017 and 2020) were likely caused by prior coral depletion, leading to a decoupling between absolute (Fig. 6B) and proportional cover loss (Fig. 6C) in these regions.

**Table 2.** Impacts of the three marine heatwaves (2016, 2017 and 2020) as absolute (% cover) and proportional (in parenthesis) coral cover loss (ie, relative to pre-bleaching total coral cover) averaged by region. Bleaching impacts result from reef-level predictions of heat stress (DHW) and simulated coral community composition.

| Year of mass bleaching | GBR | North | Central | South |
| --- | --- | --- | --- | --- |
| 2016 | -9.8 (26%) | -24.6 (63%) | -3.5 (12%) | -0.1 (0%) |
| 2017 | -5.5 (24%) | -6.2 (37%) | -8.1 (29%) | -3.1 (8%) |
| 2020 | -11.8 (40%) | -9.6 (46%) | -8.7 (33%) | -15.9 (39%) |

**Fig. 6.**
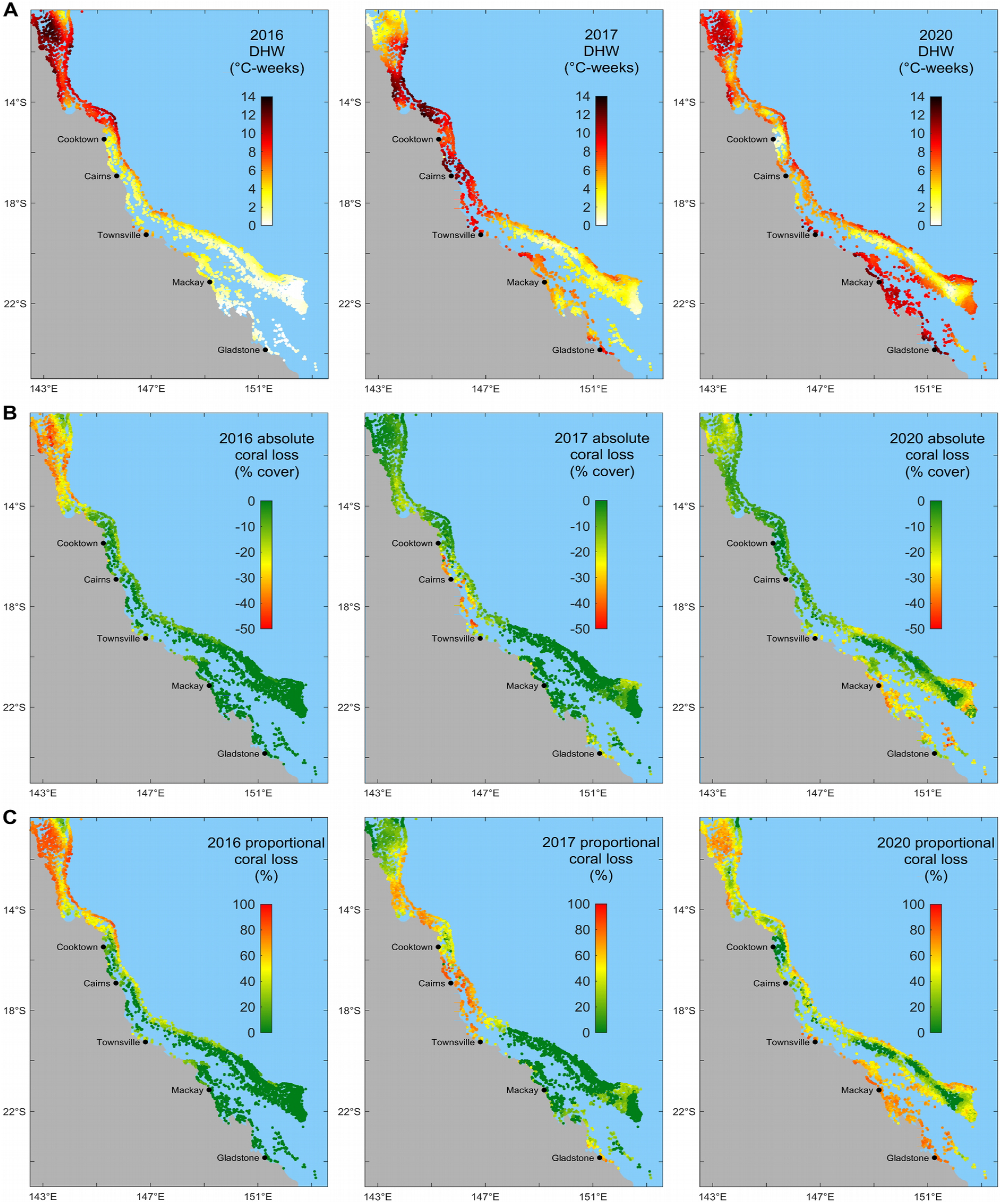
Marine heatwave (2016, 2017 and 2020) associated predictions of reef-level (**A**) heat stress (seasonal maximum DHW), (**B**) absolute loss of total coral cover and (**C**) proportional loss (i.e., relative to pre-bleaching total coral cover) averaged over n = 40 simulations.

While cyclones during 2008–2020 had relatively minor impacts across the Northern GBR, they were an important driver of coral loss in the central and southern regions (Fig. 4B, Table 1). In particular, cyclone Hamish in 2009 caused considerable impacts across the Southern GBR with an average loss of – 22.5 % cover (65% proportional cover loss), making it the most catastrophic disturbance event at a regional level during 2008–2020 (Appendix S2: Table S4). Other notable storm events included cyclones Yasi in 2011 (Central GBR), Ita in 2014 (Northern/Central GBR), Marcia in 2015 (Southern GBR) and Debbie in 2017 (mainly Central GBR). Overall, 26% of the GBR experienced less than 20% proportional loss for all individual storm events.

Impacts of CoTS outbreaks were of similar magnitude in the three regions in terms of annual absolute cover loss (between –0.4 and –0.5 % cover/y, Fig. 4B, Table 1). Because the magnitude of coral loss is dependent on initial reef states, the spatial comparison of stressor impacts requires expressing them as proportional losses relative to the pre-disturbance coral cover (Figs. 7A-C). Across the GBR, CoTS, cyclones and bleaching caused, respectively, a mean 1.8%, 7.1% and 8.5% proportional reduction of total coral cover each year (Fig. 7D). Annual proportional cover loss revealed regional differences with greater CoTS impacts in the Central GBR (2.4%/y) than in the northern (1.7%/y) and southern (1.7%/y) regions. At a reef scale, relative impacts of CoTS outbreaks were extremely patchy with severe coral mortality (> 15%/y) occurring globally in the Cairns-Cooktown area (15°S–18°S) and at the southern end of the GBR (Figs. 7A, D). The distribution of storm impacts (Figs. 7B, D) revealed a region of intense coral cover mortality (> 15%/y) between 19°S–21°S due to recurrent storm events (5-6 storms between 2008– 2020) with some particularly severe (cyclones Hamish in 2009, Marcia in 2015). Bleaching-induced mortality increased from South to North and was generally stronger (> 15%/y) on the outer reefs (Figs. 7C, D).

**Fig. 7.**
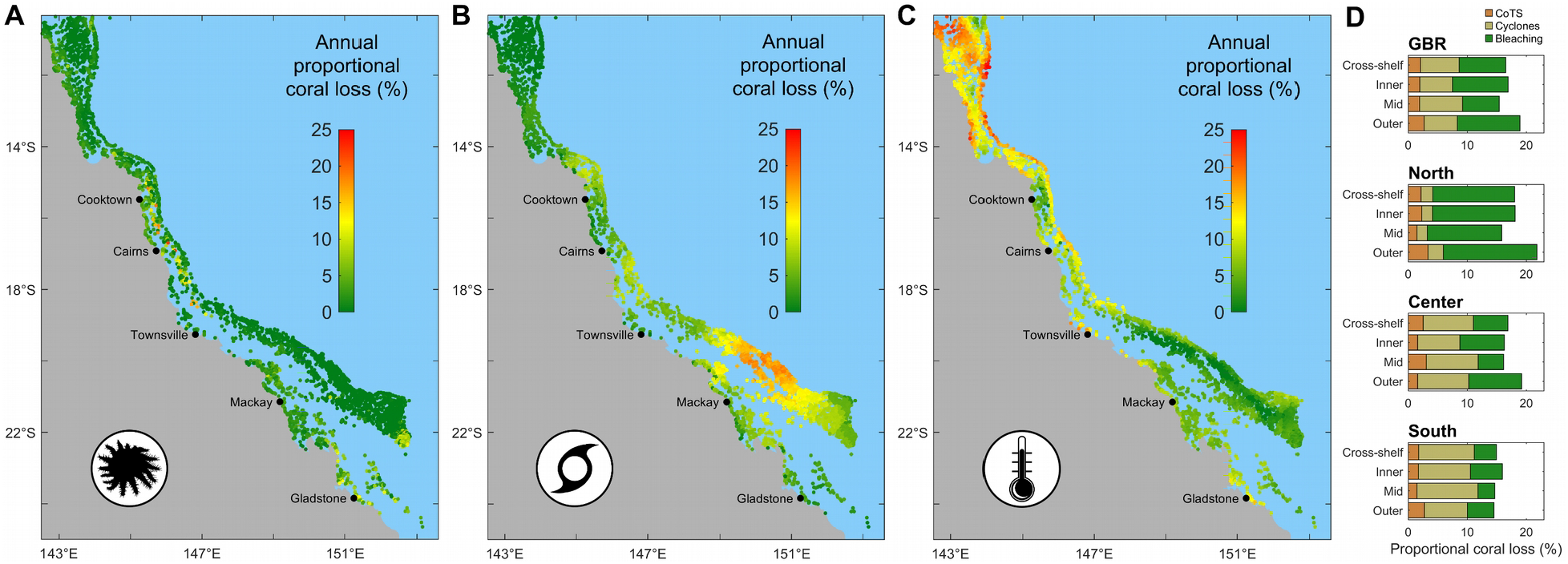
2008-2020 mean annual proportional loss of coral cover across the GBR caused by (**A**) CoTS consumption, (**A**) cyclone damages, (**C**) heat stress. (**D**) Mean annual relative cover loss per shelf position across the GBR.

### Coral recovery potential

Subtracting total annual cover loss from net annual cover change (Table 1) allowed calculating an average rate of coral cover growth for each region: ranging from +3.8 to +4.3 % cover/y. Coral community growth (g) over 6-month, extracted reef by reef before the processing of acute disturbances, was analyzed with GLMs fitted separately with every environmental predictor to assess their relative contribution on coral recovery (Appendix S2: Table S5). Total coral cover was, by far, the most important predictor of subsequent cover growth (25.0% deviance explained when fitted alone), evidenced by a quadratic influence on g (Fig. 8A). Other influential factors were sand cover (3.2% deviance explained) and the three water quality-driven demographic potentials (0.5–0.7%). The relative influence of the water quality drivers on coral recovery increased when the GLMs were fitted on inshore reefs only (2.1–3.8%, vs. 5.7% and 4.2% for coral cover and sand cover, respectively), with the percentage success of coral (i.e., Acropora) recruitment being the prominent factor. Rubble cover and the two connectivity variables (proportion of external supply and number of external links) were the least influential factors on coral recovery. In total, the eight environmental drivers accounted together for 35.6% of the deviance explained by a global GLM fitted on all reefs.

**Fig. 8.**
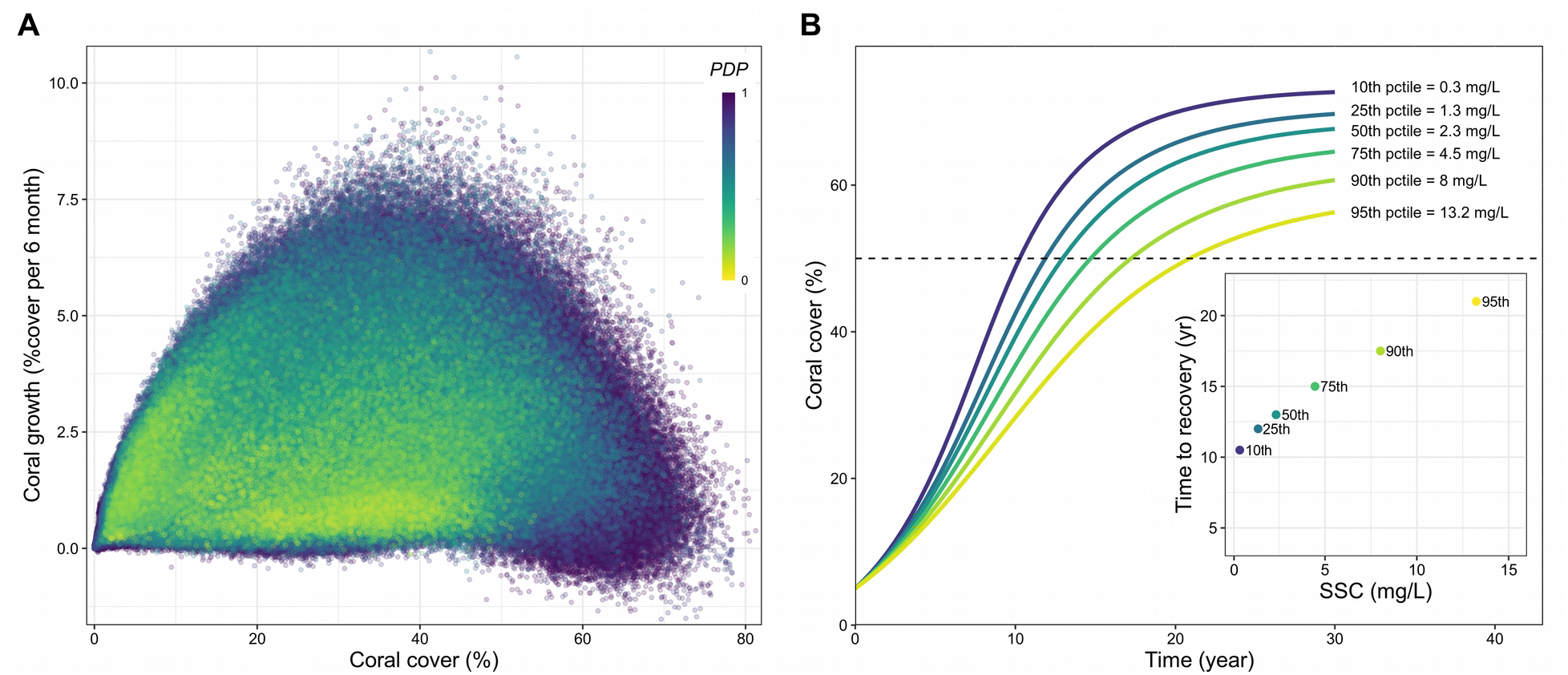
(**A**) Quadratic influence of initial total coral cover on subsequent coral cover growth rate (*g*) on all reefs during 2009–2020 (n = 3,653,760 model realizations). Color code refers to the product *PDP* of the three SSC-driven demographic potentials (reproduction, recruit survival for acroporids and juvenile growth) averaged across all available years; *PDP* ranges from 0 (no viable demographics) to 1 (full demographic performance). (**B**) GLM-based coral recovery curves for hypothetical inshore reef environments exposed to year-round SSC (mg/L), obtained by the recursive prediction of *g* (Eq. 8) from an initial coral cover of 5%. The three water-quality drivers were calculated for representative SSC values of inshore reefs (10th, 25th, 50th, 75th, 90th and 95th percentiles out of 1,374 reefs) with the other predictors set to their median value (GBR-wide across all years) – Sand: 30%; Rubble:11%; Connectnum: 8.5; Connectprop:0.06. The inset displays recovery times to 50% cover under each SSC.

Simulating coral cover growth curves from a recursive equation (Eq. 8) where growth is predicted by the global GLM revealed the impact of *SSC* on recovery dynamics on inshore reefs (Fig. 8B). From an initial 5% coral cover, growth predictions led to ∼50% coral cover after ∼10 years under steady-state (year-averaged) *SSC* < 0.3 mg/L – a concentration that corresponds to the 10^th^ percentile of inshore reefs (*n* = 1,374). Under steady-state *SSC* > 4.5 mg/L (75^th^ percentile), the same level of coral cover (i.e., 50% cover) would be achieved after a minimum of 15 years, equivalent to a 50% increase in recovery time. Recovery to 50% coral cover was delayed by ∼9 month for every 1 mg/L increment of steady-state *SSC* (inset, Fig. 8B).

### Cumulative impacts and reef resilience

The mapping of the standardized growth rate of total coral cover predicted by the GLM from 10% coral cover and reef-specific values of the environmental drivers revealed the geographic footprint of water quality (Fig. 9A). On average, the recovery potential was 14% lower inshore than offshore. On offshore reefs, the slowest growth rates were obtained in the Cairns/Cooktown region (14°S–18°S).

**Fig. 9.**
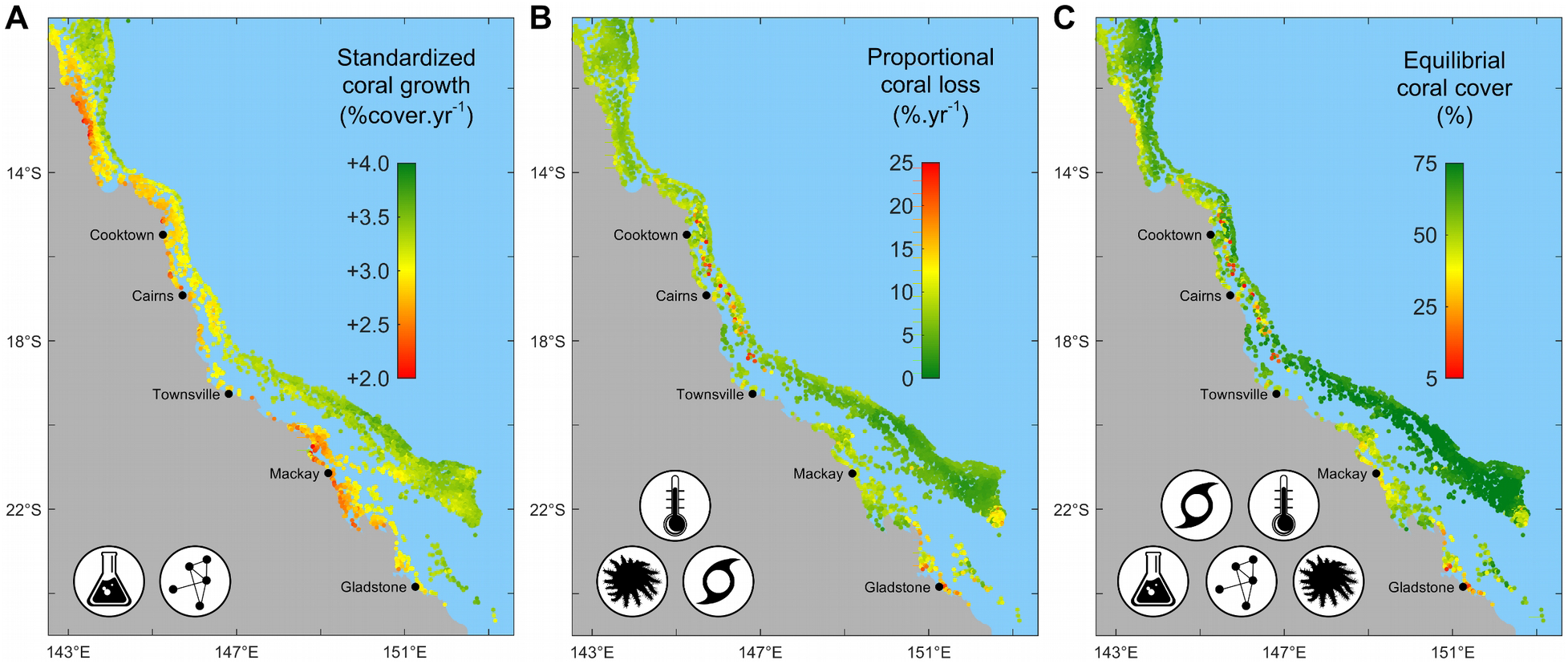
(**A**) Annual growth rate of total coral cover based on GLM predictions from a standard 10% coral cover on all reefs with the reef-specific values of early-life coral demographics (water-quality driven) and larval connectivity. (**B**) Long-term average mortality (mean annual proportional loss of total coral cover) due to CoTS, cyclones and heat stress combined. (**C**) Equilibrial cover determined from long-term simulation of growth and average mortality.

The combined rates of annual mortalities due to CoTS, cyclones and bleaching (Fig. 9B), calculated using longer-term exposures to storms (1970–2011) and heat stress (1998–2020), revealed two regions of high coral mortality (up to 25%/y): on the mid-shelf reefs of the Cairns/Cooktown region (14°S–18°S) and on the southern inshore (near Gladstone) and offshore reefs. Reefs with minimal total mortality were mostly found offshore between 20°S–22°S.

The cumulative impacts of all stressors were reflected in the computed equilibrial covers (Fig. 9C) which approximate the average value of total coral cover under local regimes of water quality, CoTS, cyclones and bleaching (Eq. 9). Using a starting cover of 30%, all reefs achieved their deterministic equilibrium in 100 years (Appendix S3: Fig. S17). The median equilibrium state was 45% coral cover on inshore reefs and 61% offshore (i.e., mid-and outer-shelf combined), reflecting the impact of water quality in the modeled coral dynamics.

## DISCUSSION

Coral populations on the GBR are distributed over a vast network of disparate reef environments, making it extremely difficult to assess the relative contribution of multiple stressors in time and space. We developed a simulation model of coral demographics to quantify the cumulative effects of multiple disturbances and explain how they drive coral cover at local and regional scales. The model integrates existing knowledge on the core underlying mechanisms of coral population dynamics with state-of-the-art spatial data capturing fine-scale environmental forcing across > 3,800 reefs. Our simulation of coral colony-scale processes under a temporally- and spatially-realistic stress regime provided a credible reconstruction of recent (2008–2020) trajectories of coral cover. Overall, the model indicated a general decline of coral cover over the past 13 years, with mass coral bleaching and cyclones dominating the simulated share of total acute stress on the GBR. The model disentangled the individual impacts of acute stressors as proportional cover losses and quantified rates of coral recovery across the entire reefscape. Spatial patterns of standardized coral cover growth highlighted the influence of suspended sediments in creating cross-shelf disparities in the potential of coral recovery. The cumulative impacts of all stressors on coral cover loss and recovery were captured within a single metric (equilibrium states) quantifying how much coral cover can be sustained on a reef given its forcing regime. Overall, our study highlights the value of mechanistic simulations for cumulative impacts assessments and management on coral reefs.

### GBR hindcast (2008–2020)

The reconstructed coral trajectories indicated a general decline of coral cover from ∼29% to ∼19%, equivalent to a loss of one third of corals in 13 years. The corresponding annual rate of absolute cover loss during 2008–2020 (–0.74% cover/y) is greater than during 1985–2012 (–0.53% cover/y) as previously calculated from monitoring data collected on 214 reefs (De’ath et al. 2012). Yet, the 1985– 2012 assessment used only AIMS manta-tow estimates of coral cover (De’ath et al. 2012) whereas our simulations were initialized with transect-equivalent coral cover values (i.e., transect and converted manta-tow estimates), which are ∼7% cover higher on average (Appendix S1). A more recent reconstruction produced a rate of annual cover loss of –1.92% cover/y between 2009–2016 (Mellin et al. 2019) based on spatially-explicit simulations of coral cover changes derived from AIMS transect-equivalent cover estimates. This rate of annual cover loss is considerably higher than the one estimated by our mechanistic simulations, yet it did not include the 2017 and 2020 bleaching events. However, inter-study comparisons are difficult as rates of absolute coral cover loss are dependent on pre-disturbance levels of coral cover, and different start- and end-points will capture a different sequence of disturbance events and recovery periods. Our reconstruction of coral trajectories provides rates of coral loss that are independent of the fluctuating baseline cover, facilitating cross-studies comparisons of the recent spatiotemporal coral dynamics on the GBR and providing a means to make future projections.

Our simulations also provide an assessment of coral reef health after the 2020 mass bleaching (Fig. 12A, Appendix S2: Table S6). We found that 22% of reefs are in a critical state (< 10% coral cover), 42% are in a poor state (10–20% coral cover) and only 19% are currently healthy (> 30% coral cover). Recent manta-tow surveys across the mid- and outer-shelf Central GBR (AIMS 2020) indicate that, by June 2020, 42% of reefs (out of 33) were in a critical state, whereas only 12% would be considered healthy using the above benchmarks. Our predictions for this region (excluding inshore reefs) yield a comparable figure based on 550 reefs after manta-tow adjustment: 39% reefs in a critical state vs. 9% healthy. Overall, the reconstructed trajectories exhibited a good agreement with the observed time-series of coral cover, recognizing that local discrepancies between reef-scale predictions and observations will inevitably arise. Some of these would constitute genuine errors in the model where a process is represented inappropriately – such as overlooking the contribution of key coral taxa – yet many will also reflect the substantive difficulty of capturing field forcing conditions in spatial layers. For example, while a cyclone track can be represented reasonably well, the dissipation of cyclone-induced wave energy around reef structures and islands is difficult to model accurately (Callaghan et al. 2020, Puotinen et al. 2020) and may fail to represent the conditions experienced by the reef from which coral cover measurements were taken. Moreover, storm damage is very patchy (Fabricius et al. 2008, Beeden et al. 2015), generating variable reef responses (Fig. 2C). Failure to predict what a reef actually experienced more likely reflects the difficulty of predicting stress exposure rather than an inappropriate demographic parameterization.

Inaccurate spatial predictions can also arise from the necessary simplification of complex coral assemblages. With coral demographic rates being representative of species typically found on offshore reef habitats, the model may underestimate coral cover on some inshore (turbid-tolerant) reefs (DeVantier et al. 2006, Browne et al. 2012). Moreover, efficient herbivore control of macroalgae was assumed despite evidence of abundant macroalgae on some inshore reefs (De’ath and Fabricius 2010, Thompson et al. 2019, Ceccarelli et al. 2020). How much this simplification affects coral cover predictions will depend on whether seaweed deter coral colonization or simply overgrow the space left vacant by coral mortality. In future, with the integration of further processes affecting reefs locally (e.g., nutrient-driven macroalgal production, realistic grazing), we expect the predictive capacity of ReefMod-GBR will improve.

### Drivers of coral loss

Measured in terms of absolute coral loss, bleaching was the most important stressor GBR-wide during 2008–2020 (–2.5 % cover/y), accounting for 49% of the stress-induced loss, while CoTS outbreaks only contributed to 11%. Manta-tow surveys (De’ath et al. 2012) for 1985–2012 found bleaching and CoTS accounted for, respectively, 10% and 42% of disturbance-driven coral mortality (see also Osborne et al. 2011 for similar figures using AIMS transect surveys between 1995–2009). The relative contribution of cyclones during 2008–2020 was similar to 1985–2012 (40% vs. 48%, respectively). The importance of cyclone impacts during both periods was partly driven by the considerable span of damage produced by Hamish (2009) in the Southern GBR, a severe cyclone with an unusual (coast-parallel) track. With three extreme heatwaves over 2008–2020 vs. only two over 1985–2012, it is no surprise that bleaching accounted for a greater share of stress-induced coral mortality in our study. We note, however, that our simulation of mass bleaching relies on mortalities observed at ∼2 m depth (Hughes et al. 2018), so represent the upper tail of the potential stress at ∼5–10 m depths. Indeed, the incidence of bleaching can decrease substantially with depth due to the attenuation of light stress (Baird et al. 2018).

In the last five years, mass coral bleaching has caused successively a proportional loss of 44% (2016– 2017 combined) and 40% (2020) of the pre-bleaching coral cover across the entire GBR. The fact that only 10% of the GBR escaped significant bleaching-induced mortality (< 20% proportional loss) raises important concerns about the ability of the GBR to cope with more frequent and intense heat stress under a warming climate (e.g., Wolff et al. 2018). Our simulations indicated that the southern region had regained most of its pre-2009 (cyclone Hamish) coral cover by the onset of the 2020 mass bleaching, despite significant loss caused by cyclone Marcia in 2015. In the northern region, the marine heatwave in 2020 erased three years of recovery (+8.4 % cover) that followed the successive impacts of the 2016– 2017 bleaching events. With a 59% proportional reduction of coral cover from 2008 to 2020, northern reefs are the main losers of the past decade. Clearly, anthropogenic bleaching has now become a key driver of coral mortality across the GBR, threatening its ability to recover from other stressors.

Impacts of CoTS outbreaks were relatively minor (–0.4 % cover/y) during 2008–2020 compared to previous assessments (–1.4 % cover/y, De’ath et al. 2012), although this period has coincided with the onset (in 2010) of the 4th cycle of CoTS outbreak since 1960s (Pratchett et al. 2014). Given that CoTS density has been surveyed for only 2% of the GBR, we relied on the random initialization of CoTS populations derived from the spatial predictions of the CoCoNet model (Condie et al. 2018) with the subsequent dynamics driven by larval dispersal and coral abundance. The importance of nutrient-enhanced larval survival in the initiation of CoTS outbreaks is still debated (Pratchett et al. 2014, 2017, Wolfe et al. 2017), and it is noteworthy that survival of CoTS larvae predicted by chlorophyll simulations over eight spawning seasons (2010–2018) was very low in the Cairns–Cooktown area (Fig. 3A, Appendix S3: Fig. S11), a region where all four CoTS outbreaks appear to have initiated (Brodie et al. 2005, Pratchett et al. 2014). Comparisons between eReefs predictions and in situ measurements have revealed a tendency of the model to locally underestimate nutrient and chlorophyll concentrations (Robson et al. 2020). On the other hand, CoTS likely started their gradual build-up several years before the first detection of outbreaking densities in 2010. While eReefs predictions were only available from December 2010, large river floods in this region during CoTS spawning in 2008 and 2009 had the potential of developing primary outbreaks (Fabricius et al. 2010).

High chlorophyll concentrations were prevalent in the southern section of the GBR (Swains and Capricorn/Bunker sectors), both on inner and outer reefs (Appendix S3: Fig. S10). Inshore, this is likely due to runoff events with a culmination during the 2010–2011 wet season (Appendix S3: Fig. S11). This facilitated the propagation of CoTS populations created at initialization, although there is currently no evidence of CoTS outbreaks on southern inner reefs (Thompson et al. 2019). On southern offshore reefs, high Chl a is the result of recurrent intrusions of nutrient-rich waters by upwelling on the shelf break (e.g., Andrews and Furnas 1986, Berkelmans et al. 2010), and it has been hypothesized that primary outbreaks could emerge there with no relation to river-flood events (Moran et al. 1988, Johnson 1992, Miller et al. 2015). Although the causes of primary outbreaks on the GBR are yet to be resolved (Pratchett et al. 2014, 2017), the present model can be used to explore the timing and mechanisms of the propagation of secondary outbreaks facilitated by nutrient availability (Brodie et al. 2017).

### Drivers of coral recovery

The population growth rates that emerged from colony-scale dynamics revealed which environmental factors contributed most to the expansion of coral cover. First and foremost is the influence of initial coral cover which determines the subsequent rate of increase in coral cover, corroborating empirical observations (Graham et al. 2011, Ortiz et al. 2018). With a fixed rate of radial extension, the areal growth increment is greater for larger colonies than for smaller ones, so that, at least at the initial stage of coral colonization, the rate of cover growth becomes gradually faster as corals get bigger. As large and sexually-mature colonies become more prevalent, self-recruitment intensifies because more offspring are produced, so that population size increases and amplifies the rate of cover growth. Subsequently, coral colonization reduces the space available for recruitment (Fig. 2B) and colony extension, thereby slowing down the rate of increase in coral cover until the colonization space is saturated (Fig. 8A). As a result, the influence of initial coral cover on subsequent growth is non-linear and creates a sigmoid recovery curve (Fig. 2A) that is typically observed in *Acropora*-dominated communities (Halford et al. 2004, Emslie et al. 2008). We captured these dynamics at the community scale, first through the statistical modeling of the stepwise changes of total coral cover, then using the resulting model (GLM) to predict cover growth increments and reconstruct coral recovery curves. This enabled the integration of influential drivers of coral growth such as suspended sediments (Fig. 8B) and allowed the systematic exploration of the potential of coral recovery across the entire reefscape (Fig. 9A). This growth model offers an alternative to heuristic inferences of recovery dynamics based on statistical model fits that depend on data availability (Thompson and Dolman 2010, Osborne et al. 2011, 2017, Wolff et al. 2018, Mellin et al. 2019).

Once standardized with the GLM, spatial variations in coral growth revealed the negative impacts of suspended sediments on the recovery potential of inshore reefs. This is consistent with recent analyses (Ortiz et al. 2018, MacNeil et al. 2019) that found reductions in coral cover growth rates with the extent of river flood plumes assessed by satellite imagery. We note, however, that high *SSC* values can also result from wind-driven resuspension of fine sediments as observed during the dry season (Appendix S3: Fig S4B). Our assessment of water quality impacts is based on predictions of transport, sinking and re-suspension of fine (30 μm with a sinking rate of 17m) sediments from hydrodynamic modeling. This enables *SSC* exposure to be integrated over time periods (days to months) that are relevant to the sensitive stages of coral ontogeny (Humanes et al. 2017a, 2017b), allowing physiological impacts to be scaled up to the ecosystem level. Retaining 10 years as a standard recovery time under good water quality conditions (mean annual *SSC* < 0.3 mg/L, corresponding to 10% of inshore reefs), our simulations indicate that an increment of 1 mg/L of steady-state *SSC* retards coral recovery by 9 month (Fig. 8B). While these predictions can help setting water quality targets for management, they are likely biased toward a specific response of acroporids (Appendix S1) and remain to be tested *in situ*. However, detecting these impacts on coral cover is challenging: this would require extended time series as the deleterious effects of *SSC* might only become apparent after a long period of uninterrupted recovery. Although being representative of steady-state *SSC* exposures (annual averages at 4km resolution), our simulated recovery rates are standardized to a given coral cover and can be used to compare the recovery potential (Fig. 9A) and resilience (Fig. 9C) among reefs.

Although larval connectivity is widely regarded as an important driver of coral recovery, a quantitative link between larval supply and coral cover dynamics is yet to be established. Here, larval connectivity had little influence on the reconstructed coral cover growth, but this does not imply that external larval supply is not demographically important. With the current parameterization of larval retention (i.e., a minimum 28% of larvae produced by a reef is retained), the contribution of external supply to total settlement is globally low: based on the transition probabilities (i.e., without accounting for the actual number of larvae produced), external supply represented 6% of larval supply for 50% of the reefs (mean: 15%). Because coral settlement was modeled as a saturating function of larval supply, self-supply was generally sufficient for the making of settlement. There is, however, considerable uncertainty in the set value of larval retention, with likely variations from reef to reef (Black 1993). Moreover, the relative importance of self-recruitment would likely decrease after severe coral mortality, making external supply a key process for local recovery. Future work should model larval dispersal at a finer spatial resolution (i.e., < 1 km) for a better evaluation of the relative contribution of self vs external supply. This information is critical to capture the demographic impacts of larval connectivity and support connectivity-based management interventions.

### Cumulative impacts on coral loss and recovery

Expressing stress-induced coral mortality as proportional loss was key to assessing the spatial distribution of the individual and combined impacts of acute disturbances. This yielded vulnerability maps that reflect the frequency and intensity of recent disturbances contextualized within the coral community composition predicted by the model, while being independent of the levels of coral cover at the time of disturbances. The spatial predictions of standardized coral growth and stress-induced coral mortality allowed computation of the equilibrium state for > 3,800 reefs. Equilibrium states can be viewed as long-term averages around which coral cover fluctuates in a given reef environment. They integrate the combined effects of chronic (water quality) and acute stress (bleaching, cyclones, and CoTS), and their use here is to reveal large-scale patterns in the resilience of the ecosystem (Fig. 9C). Yet, since coral reefs are non-equilibrial systems that frequently experience acute impacts (Done 1992, Connell 1997), the transient state of reefs can be far higher or lower than their long-term equilibrium. With this in mind, the notion of equilibrium state differs from the concept of carrying capacity (the intrinsic limit of a population) as a reef can exhibit episodically higher levels of coral cover until stress-induced mortality brings the reef closer to its equilibrial cover value.

Although equilibria were created by running the model for 100 years, they do not constitute projections for future reef health; they merely set regional expectations for the relative state of the system based on recent stress intensities and frequencies. Like for any resilience metric, transient stress regimes clearly challenge these expectations (i.e., intensifying heat stress), and projecting equilibrium states would require integrating specific forecast scenarios of disturbances into their calculation. Moreover, the present metric of resilience does not account for competitive interactions (e.g., with macroalgae or soft corals) which will favor emergence of multiple equilibrium states (McManus and Polsenberg 2004, Mumby et al. 2007). Future model versions will spatially integrate grazing and macroalgal productivity to assess ecological resilience (*sensu* Holling 1996: the ability to move towards alternate community types) and define ecological thresholds of coral persistence (Mumby et al. 2007, 2014, Bozec et al. 2016) across the GBR.

### Mechanistic approach to cumulative effects assessment

Cumulative impacts on coral reefs have been traditionally assessed through the analysis of monitored coral cover changes attributed to specific stressors. Yet, disentangling the individual effects of multiple drivers requires extensive monitoring data due to inherent difficulties in attributing causality to observed coral changes (Fabricius and De’ath 2004). Moreover, impacts that manifest as a slowing down of coral growth are easily overlooked by monitoring. While these can be evidenced at the scale of individual colonies in controlled environments, experimental designs can only manipulate a small number of stressors and have a limited ability to infer responses at the community level (Hodgson and Halpern 2019). We show that mechanistic simulations that integrate key demographic processes provide important insights to cumulative effects assessments. Here, the core mechanisms underlying coral demography were simulated at the scale of coral colonies to quantify stressor impacts on specific biological processes and developmental life stages. This enabled the emergence of complex interactions and feedbacks that compound the cumulative effects of multiple drivers and determine the dynamics of coral cover. Whilst incomplete knowledge on key demographic parameters has inhibited individual-based approaches for cumulative impacts assessments, we address this issue by providing a suite of empirical relationships between common stressors and coral demographics to promote a mechanistic evaluation of coral reef health.

To meet the challenge of understanding the behavior emerging from colony-scale simulations, we used statistical approaches that disentangled the contribution of different drivers to coral cover changes. Note that this does not imply perfect mechanistic knowledge overall; what holds is that given our current knowledge on how these mechanisms operate individually, we can aim to understand how they interact in driving coral cover virtually. Testing these predictions empirically will be difficult at any scale, yet the grounding in underlying mechanisms combined with the successful validation of model behavior provides a basis for making future predictions outside of the input model parameter space.

### Implications for reef monitoring and resilience-based management

Managing for coral resilience requires evaluating the current state of reefs, their exposure to disturbances and their ability to recover from those pressures. Our simulations predict the current state of > 3,800 reefs on the GBR based on mechanistic expectations and spatio-temporal data on drivers. They provide an assessment in space and time of the stress regime of each reef covering both chronic environmental forcing (water quality and larval connectivity) and acute mortality events. This portfolio of reef vulnerability across the GBR can be combined with present-day spatial predictions of coral cover (Fig. 10A), community composition and demographic structure, and potential for coral recovery (incorporating exposure to CoTS and loose coral rubble) to complement reef monitoring. This is especially important considering that existing monitoring only represents ∼40% only of the environmental regimes of the GBR (Mellin et al. 2020). While the present model informs about recent trends and status of unmonitored reef areas (∼96% of the 3,806 reefs between 2008–2020), it can also help designing more representative and efficient coral and CoTS surveillance programs in support of reef management.

**Fig. 10.**
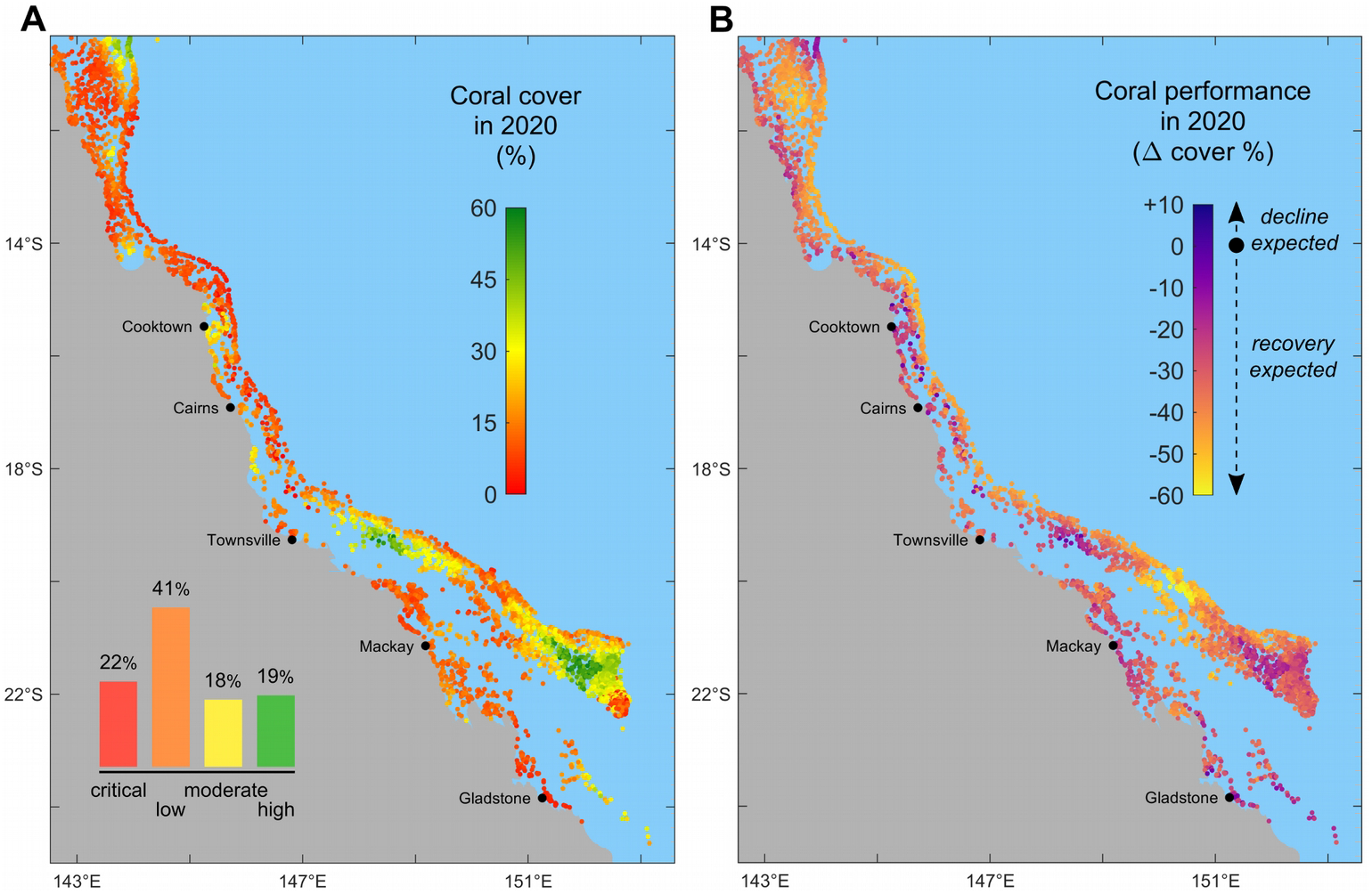
(**A**) Present-day (2020) model predictions of total coral cover. Inset: GBR-wide distribution of reef health status: critical (<10% coral cover); low (10–20%); moderate (20–30%); high (>30%). (**B**) coral performance in 2020 as the difference between total coral cover and simulated equilibrial coral cover. A positive performance value indicates that present-day coral cover on a reef is greater than expected under its regime of disturbance and recovery; a decline is expected in a near future. Inversely, under-performing reefs (i.e., negative performance values) are expected to recover closer or beyond their equilibrium.

Of particular significance for an improved management of the GBR is the equilibrial cover as a metric of reef resilience. While recognizing the limits of predicting coral cover for non-equilibrial systems, equilibrium states set expectations of future changes in the short term: a coral cover value higher than the reef’s equilibrium state indicates that the reef is performing better than expected, a performance that is unlikely to persist. Inversely, a reef that is largely under-performing relative to its equilibrium state is expected to recover beyond the equilibrial value. Comparing the current and potential performance of reefs (Fig. 10B) may help identify those most likely to respond to interventions and sustain improvements over the longer term.

With ReefMod-GBR, we provide a simulation tool to evaluate management scenarios and help developing a structured decision-making process. Multiple scenarios of stress mitigation and/or restoration can be simulated and their performance compared in time and space using an array of model variables (e.g., coral cover, mortality and recovery rates, CoTS density). To this aim, the equilibrial cover is an operational metric that can capture changes in cumulative impacts in response to a given intervention. Equilibrial cover pertains to the associated regime of disturbance, so that a relaxation of acute (e.g., CoTS control) and chronic stress (e.g., water quality improvement) would lead to a different equilibrium. Expanding the model with projections of carbon emissions will provide opportunities for exploring management strategies under climate change, and for prioritizing tactical interventions with the greatest benefits to the resilience of the GBR.

## ACKNOWLEDGMENTS

This research was funded by the Australian Research Council (ARC) Centre of Excellence for Coral Reef Studies, the Great Barrier Reef Foundation (GBRF), an ARC Discovery Project, the Australian Government’s National Environmental Science Programme (NESP 4.5 and 3.1.1), the Reef Restoration and Adaptation Project (RRAP) case, and a government consortium (Department of Environment, GBRMPA and the Queensland Government) to PJM, Y-MB and KH. The current field used for connectivity calculations and the sediment and chlorophyll fields were developed in the *eReefs* Project, a public-private collaboration between Australia’s leading operational and scientific research agencies, government, and corporate Australia. We thank C. Doropoulos, G. Roff, N. Wolff, K.R.N. Anthony and B. Schaffelke for fruitful discussions.

## REFERENCES

Andrews, J. C., and M. J. Furnas. 1986. Subsurface intrusions of Coral Sea water into the central Great Barrier Reef–I. Structures and shelf-scale dynamics. Continental Shelf Research 6:491–514.

AIMS (Australian Institute of Marine Science). 2020. Annual Summary Report on coral reef condition for 2019/20. Long-term Reef Monitoring Program. Australian Institute of Marine Science.

Babcock, R. C., G. D. Bull, P. L. Harrison, A. J. Heyward, J. K. Oliver, C. C. Wallace, and B. L. Willis. 1986. Synchronous spawnings of 105 scleractinian coral species on the Great Barrier Reef. Marine Biology 90:379– 394.

Babcock, R. C., and C. N. Mundy. 1992. Reproductive biology, spawning and field fertilization rates of Acanthaster planci. Marine and Freshwater Research 43:525–533.

Baird, A. H., and P. Marshall. 2002. Mortality, growth and reproduction in scleractinian corals following bleaching on the Great Barrier Reef. Marine Ecology Progress Series 237:133–141.

Baird, A. H., J. S. Madin, M. Álvarez-Noriega, L. Fontoura, J. T. Kerry, C.-Y. Kuo, K. Precoda, D. Torres-Pulliza, R. M. Woods, K. J. Zawada, and others. 2018. A decline in bleaching suggests that depth can provide a refuge from global warming in most coral taxa. Marine Ecology Progress Series 603:257–264.

Baird, M. E., M. P. Adams, J. Andrewartha, N. Cherukuru, M. Gustafsson, S. Hadley, M. Herzfeld, E. Jones, N. Margvelashvili, M. Mongin, and others. 2017. CSIRO environmental modelling suite: scientific description of the optical, carbon chemistry and biogeochemical models (BGC1p0). https://research.csiro.au/ereefs/wp-content/uploads/sites/34/2015/08/eReefsOpticalBGCframeBGC1p0.pdf.

Ban, S. S., N. A. Graham, and S. R. Connolly. 2014. Evidence for multiple stressor interactions and effects on coral reefs. Global Change Biology 20:681–697.

Baria, M. V. B., R. D. Villanueva, and J. R. Guest. 2012. Spawning of three-year-old Acropora Millepora corals reared from larvae in northwestern Philippines. Bulletin of Marine Science 88:61–62.

Beeden, R., J. Maynard, M. Puotinen, P. Marshall, J. Dryden, J. Goldberg, and G. Williams. 2015. Impacts and recovery from severe tropical Cyclone Yasi on the Great Barrier Reef. PLoS ONE 10:e0121272.

Berkelmans, R. 2002. Time-integrated thermal bleaching thresholds of reefs and their variation on the Great Barrier Reef. Marine Ecology Progress Series 229:73–82.

Berkelmans, R., G. De’ath, S. Kininmonth, and W. J. Skirving. 2004. A comparison of the 1998 and 2002 coral bleaching events on the Great Barrier Reef: spatial correlation, patterns, and predictions. Coral Reefs 23:74–83.

Berkelmans, R., S. J. Weeks, and C. R. Steinberga. 2010. Upwelling linked to warm summers and bleaching on the Great Barrier Reef. Limnology and Oceanography 55:2634–2644.

Biggs, B. C. 2013. Harnessing natural recovery processes to improve restoration outcomes: an experimental assessment of sponge-mediated coral reef restoration. PLoS ONE 8:e64945.

Birkeland, C., and J. Lucas. 1990. Acanthaster planci: major management problem of coral reefs. CRC press.

Black, K. P. 1993. The relative importance of local retention and inter-reef dispersal of neutrally buoyant material on coral reefs. Coral Reefs 12:43–53.

Bozec, Y.-M., and P. J. Mumby. 2015. Synergistic impacts of global warming on the resilience of coral reefs. Philosophical Transactions of the Royal Society B 370:20130267.

Bozec, Y.-M., L. Alvarez-Filip, and P. J. Mumby. 2015. The dynamics of architectural complexity on coral reefs under climate change. Global Change Biology 21:223–235.

Bozec, Y.-M., S. O’Farrell, J. H. Bruggemann, B. E. Luckhurst, and P. J. Mumby. 2016. Tradeoffs between fisheries harvest and the resilience of coral reefs. Proceedings of the National Academy of Sciences 113:4536–4541.

Bozec, Y.-M., and P. J. Mumby. 2019. Detailed description of ReefMod-GBR and simulation results. Pages 72–113 Reef Restoration and Adaptation Program: Modelling Methods and Findings. A report provided to the Australian Government. Townsville, Australia.

Bozec, Y.-M., C. Doropoulos, G. Roff, and P. J. Mumby. 2019. Transient grazing and the dynamics of an unanticipated coral–algal phase shift. Ecosystems 22:296–311.

Bozec, Y. M., and P. J. Mumby. 2020. Coral reef models as assessment and reporting tools for the Reef 2050 Integrated Monitoring and Reporting Program–a review. Great Barrier Reef Marine Park Authority.

Brodie, J., K. Fabricius, G. De’ath, and K. Okaji. 2005. Are increased nutrient inputs responsible for more outbreaks of crown-of-thorns starfish? An appraisal of the evidence. Marine Pollution Bulletin 51:266–278.

Brodie, J., and J. Waterhouse. 2012. A critical review of environmental management of the ‘not so Great’ Barrier Reef. Estuarine, Coastal and Shelf Science 104:1–22.

Brodie, J., M. Devlin, and S. Lewis. 2017. Potential enhanced survivorship of crown of thorns starfish larvae due to near-annual nutrient enrichment during secondary outbreaks on the central mid-shelf of the Great Barrier Reef, Australia. Diversity 9:17.

Browne, N. K., S. G. Smithers, and C. T. Perry. 2012. Coral reefs of the turbid inner-shelf of the Great Barrier Reef, Australia: an environmental and geomorphic perspective on their occurrence, composition and growth. Earth-Science Reviews 115:1–20.

Callaghan, D. P., P. J. Mumby, and M. S. Mason. 2020. Near-reef and nearshore tropical cyclone wave climate in the Great Barrier Reef with and without reef structure. Coastal Engineering 157:103652.

Ceccarelli, D. M., R. D. Evans, M. Logan, P. Mantel, M. Puotinen, C. Petus, G. R. Russ, and D. H. Williamson. 2020. Long-term dynamics and drivers of coral and macroalgal cover on inshore reefs of the Great Barrier Reef Marine Park. Ecological Applications 30:e02008.

Cheal, A. J., M. A. MacNeil, M. J. Emslie, and H. Sweatman. 2017. The threat to coral reefs from more intense cyclones under climate change. Global Change Biology 23:1511–1524.

Condie, S. A.É. E. Plagányi, E. B. Morello, K. Hock, and R. Beeden. 2018. Great Barrier Reef recovery through multiple interventions. Conservation Biology 32:1356–1367.

Connell, J. H. 1997. Disturbance and recovery of coral assemblages. Coral Reefs 16:S101–S113.

Connolly, S. R., and A. H. Baird. 2010. Estimating dispersal potential for marine larvae: dynamic models applied to scleractinian corals. Ecology 91:3572–3583.

Côté, I. M., J. A. Gill, T. A. Gardner, and A. R. Watkinson. 2005. Measuring coral reef decline through meta-analyses. Philosophical Transactions of the Royal Society B: Biological Sciences 360:385–395.

Crain, C. M., K. Kroeker, and B. S. Halpern. 2008. Interactive and cumulative effects of multiple human stressors in marine systems. Ecology Letters 11:1304–1315.

Darling, E. S., and I.M. Côté. 2008. Quantifying the evidence for ecological synergies. Ecology Letters 11:1278– 1286.

Darling, E. S., T. R. McClanahan, and I.M. Côté. 2013. Life histories predict coral community disassembly under multiple stressors. Global Change Biology 19:1930–1940.

De’ath, G., and P. J. Moran. 1998. Factors affecting the behaviour of crown-of-thorns starfish (Acanthaster planci L.) on the Great Barrier Reef: 2: Feeding preferences. Journal of Experimental Marine Biology and Ecology 220:107–126.

De’ath, G., and K. Fabricius. 2010. Water quality as a regional driver of coral biodiversity and macroalgae on the Great Barrier Reef. Ecological Applications 20:840–850.

De’ath, G., K. E. Fabricius, H. Sweatman, and M. Puotinen. 2012. The 27–year decline of coral cover on the Great Barrier Reef and its causes. Proceedings of the National Academy of Sciences 109:17995–17999.

DeVantier, L. M., G. De’Ath, E. Turak, T. J. Done, and K. E. Fabricius. 2006. Species richness and community structure of reef-building corals on the nearshore Great Barrier Reef. Coral Reefs 25:329–340.

Done, T. J. 1992. Phase shifts in coral reef communities and their ecological significance. Hydrobiologia 247:121– 132.

Done, T. J. 1995. Ecological criteria for evaluating coral reefs and their implications for managers and researchers. Coral Reefs 14:183–192.

Doropoulos, C., S. Ward, G. Roff, M. González-Rivero, and P. J. Mumby. 2015. Linking demographic processes of juvenile corals to benthic recovery trajectories in two common reef habitats. PLoS ONE 10:e0128535.

Doropoulos, C., G. Roff, Y.-M. Bozec, M. Zupan, J. Werminghausen, and P. J. Mumby. 2016. Characterizing the ecological trade–offs throughout the early ontogeny of coral recruitment. Ecological Monographs 86:20–44.

Eakin, C. M., J. A. Morgan, S. F. Heron, T. B. Smith, G. Liu, L. Alvarez-Filip, B. Baca, E. Bartels, C. Bastidas, and C. Bouchon. 2010. Caribbean corals in crisis: record thermal stress, bleaching, and mortality in 2005. PLoS ONE 5:e13969.

Edmunds, P. J., and B. Riegl. 2020. Urgent need for coral demography in a world where corals are disappearing. Marine Ecology Progress Series 635:233–242.

Edwards, H. J., I. A. Elliott, C. M. Eakin, A. Irikawa, J. S. Madin, M. McField, J. A. Morgan, R. van Woesik, and P.J. Mumby. 2011. How much time can herbivore protection buy for coral reefs under realistic regimes of hurricanes and coral bleaching? Global Change Biology 17:2033–2048.

Emslie, M., A. Cheal, H. Sweatman, and S. Delean. 2008. Recovery from disturbance of coral and reef fish communities on the Great Barrier Reef, Australia. Marine Ecology Progress Series 371:177–190.

Engelhardt, U., M. Hartcher, J. Cruise, D. Engelhardt, M. Russell, N. Taylor, G. Thomas, and D. Wiseman. 1999. Finescale surveys of crown-of-thorns starfish (Acanthaster planci) in the central Great Barrier Reef region. CRC Reef Research Centre technical report.

Engelhardt, U., M. Hartcher, N. Taylor, J. Cruise, D. Engelhardt, M. Russel, I. Stevens, G. Thomas, D. Williamson, and D. Wiseman. 2001. Crown-of-thorns starfish (Acanthaster planci) in the central Great Barrier Reef region. Results of fine-scale surveys conducted in 1999-2000. Technical Report.

Evans, R. D., S. K. Wilson, R. Fisher, N. M. Ryan, R. Babcock, D. Blakeway, T. Bond, P. Dorji, F. Dufois, P. Fearns, and others. 2020. Early recovery dynamics of turbid coral reefs after recurring bleaching events. Journal of Environmental Management 268:110666.

Fabricius, K. E., and G. De’ath. 2004. Identifying ecological change and its causes: a case study on coral reefs. Ecological Applications 14:1448–1465.

Fabricius, K. E. 2005. Effects of terrestrial runoff on the ecology of corals and coral reefs: review and synthesis. Marine Pollution Bulletin 50:125–146.

Fabricius, K. E., G. De’Ath, M. L. Puotinen, T. Done, T. F. Cooper, and S. C. Burgess. 2008. Disturbance gradients on inshore and offshore coral reefs caused by a severe tropical cyclone. Limnology and Oceanography 53:690– 704.

Fabricius, K., K. Okaji, and G. De’Ath. 2010. Three lines of evidence to link outbreaks of the crown-of-thorns seastar Acanthaster planci to the release of larval food limitation. Coral Reefs 29:593–605.

Filbee-Dexter, K., and T. Wernberg. 2018. Rise of turfs: a new battlefront for globally declining kelp forests. BioScience 68:64–76.

Fox, H. E., J. S. Pet, R. Dahuri, and R. L. Caldwell. 2003. Recovery in rubble fields: long-term impacts of blast fishing. Marine Pollution Bulletin 46:1024–1031.

Fox, R. J., and D. R. Bellwood. 2007. Quantifying herbivory across a coral reef depth gradient. Marine Ecology Progress Series 339:49–59.

Graham, N. A. J., K. L. Nash, and J. T. Kool. 2011. Coral reef recovery dynamics in a changing world. Coral Reefs 30:283–294.

Graham, N. A., S. Jennings, M. A. MacNeil, D. Mouillot, and S. K. Wilson. 2015. Predicting climate-driven regime shifts versus rebound potential in coral reefs. Nature 518:94–97.

GBRMPA (Great Barrier Reef Marine Park Authority). 2007. Great Barrier Reef (GBR) Features (Reef boundaries, QLD Mainland, Islands, Cays, Rocks and Dry Reefs). eAtlas. https://eatlas.org.au/data/uuid/ac8e8e4f-fc0e-4a01-9c3d-f27e4a8fac3c.

GBRMPA (Great Barrier Reef Marine Park Authority). Great Barrier Reef Outlook Report 2019. GBRMPA, Townsville.

Haddon, M. 2011. Modelling and quantitative methods in fisheries. CRC Press/Chapman and Hall.

Halford, A., A. J. Cheal, D. Ryan, and D. McB. 2004. Resilience to Large-Scale Disturbance in Coral and Fish Assemblages on the Great Barrier Reef. Ecology:1892–1905.

Hall, V., and T. Hughes. 1996. Reproductive strategies of modular organisms: comparative studies of reef-building corals. Ecology 77:950–963.

Halpern, B. S., and R. Fujita. 2013. Assumptions, challenges, and future directions in cumulative impact analysis. Ecosphere 4:1–11.

Harborne, A. R., A. Rogers, Y. M. Bozec, and P. J. Mumby. 2017. Multiple Stressors and the Functioning of Coral Reefs. Annual Review of Marine Science 9:445–468.

Heron, S., L. Johnston, G. Liu, E. Geiger, J. Maynard, J. De La Cour, S. Johnson, R. Okano, D. Benavente, and T. Burgess. 2016. Validation of reef-scale thermal stress satellite products for coral bleaching monitoring. Remote Sensing 8:59.

Herzfeld, M., J. Andrewartha, M. Baird, R. Brinkman, M. Furnas, P. Gillibrand, M. Hemer, K. Joehnk, E. Jones, D. McKinnon, N. Margvelashvili, M. Mongin, P. Oke, F. Rizwi, B. Robson, S. Seaton, J. Skerratt, H. Tonin, and K. Wild-Allen. 2016. eReefs Marine Modelling: Final Report. Page 497 pp. CSIRO, Hobart.

Hock, K., N. H. Wolff, J. C. Ortiz, S. A. Condie, K. R. Anthony, P. G. Blackwell, and P. J. Mumby. 2017. Connectivity and systemic resilience of the Great Barrier Reef. PLoS Biology 15:e2003355.

Hock, K., C. Doropoulos, R. Gorton, S. A. Condie, and P. J. Mumby. 2019. Split spawning increases robustness of coral larval supply and inter-reef connectivity. Nature Communications 10:3463.

Hodgson, E. E., and B. S. Halpern. 2019. Investigating cumulative effects across ecological scales. Conservation Biology 33:22–32.

Hoegh-Guldberg, O., P. J. Mumby, A. J. Hooten, R. S. Steneck, P. Greenfield, E. Gomez, C. D. Harvell, P. F. Sale, A. J. Edwards, and K. Caldeira. 2007. Coral reefs under rapid climate change and ocean acidification. Science 318:1737–1742.

Holling, C. S. 1996. Engineering Resilience versus Ecological Resilience. Engineering Within Ecological Constraints 31:32.

Hughes, T. P., A. H. Baird, D. R. Bellwood, M. Card, S. R. Connolly, C. Folke, R. Grosberg, O. Hoegh-Guldberg, J. B. C. Jackson, and J. Kleypas. 2003. Climate change, human impacts, and the resilience of coral reefs. Science 301:929–933.

Hughes, T. P., and J. H. Connell. 1999. Multiple stressors on coral reefs: A long–term perspective. Limnology and Oceanography 44:932–940.

Hughes, T. P., and J. E. Tanner. 2000. Recruitment failure, life histories, and long-term decline of Caribbean corals. Ecology 81:2250–2263.

Hughes, T. P., D. R. Bellwood, A. H. Baird, J. Brodie, J. F. Bruno, and J. M. Pandolfi. 2011. Shifting base-lines, declining coral cover, and the erosion of reef resilience: comment on Sweatman et al.(2011). Coral Reefs 30:653–660.

Hughes, T. P., J. T. Kerry, M. Álvarez-Noriega, J.G. Álvarez-Romero, K. D. Anderson, A. H. Baird, R. C. Babcock, M. Beger, D. R. Bellwood, R. Berkelmans, and others. 2017. Global warming and recurrent mass bleaching of corals. Nature 543:373.

Hughes, T. P., J. T. Kerry, A. H. Baird, S. R. Connolly, A. Dietzel, C. M. Eakin, S. F. Heron, A. S. Hoey, M. O. Hoogenboom, G. Liu, and others. 2018. Global warming transforms coral reef assemblages. Nature 556:492.

Humanes, A., A. Fink, B. L. Willis, K. E. Fabricius, D. de Beer, and A. P. Negri. 2017a. Effects of suspended sediments and nutrient enrichment on juvenile corals. Marine Pollution Bulletin 125:166–175.

Humanes, A., G. F. Ricardo, B. L. Willis, K. E. Fabricius, and A. P. Negri. 2017b. Cumulative effects of suspended sediments, organic nutrients and temperature stress on early life history stages of the coral Acropora tenuis. Scientific Reports 7:44101.

Johnson, C. 1992. Settlement and recruitment of Acanthatser planci on the Great Barrier Reef: Questions of process and scale. Marine and Freshwater Research 43:611–627.

Jones, R., G. Ricardo, and A. Negri. 2015. Effects of sediments on the reproductive cycle of corals. Marine Pollution Bulletin 100:13–33.

Keesing, J. K., and A. R. Halford. 1992. Importance of postsettlement processes for the population dynamics of Acanthaster planci (L.). Marine and Freshwater Research 43:635–651.

Keesing, J., and J. Lucas. 1992. Field measurement of feeding and movement rates of the crown-of-thorns starfish Acanthaster planci (L.). Journal of Experimental Marine Biology and Ecology 156:89–104.

Kettle, B., and J. Lucas. 1987. Biometric relationships between organ indices, fecundity, oxygen consumption and body size in Acanthaster planci (L.)(Echinodermata; Asteroidea). Bulletin of Marine Science 41:541–551.

Kuffner, I. B., L. J. Walters, M. A. Becerro, V. J. Paul, R. Ritson-Williams, and K. S. Beach. 2006. Inhibition of coral recruitment by macroalgae and cyanobacteria. Marine Ecology Progress Series 323:107–117.

Lam, V. Y., M. Chaloupka, A. Thompson, C. Doropoulos, and P. J. Mumby. 2018. Acute drivers influence recent inshore Great Barrier Reef dynamics. Proceedings of the Royal Society B: Biological Sciences 285:20182063.

Lam, V. Y., C. Doropoulos, Y.-M. Bozec, and P. J. Mumby. 2020. Resilience Concepts and Their Application to Coral Reefs. Frontiers in Ecology and Evolution 8:49.

Liu, G., W. J. Skirving, E. F. Geiger, J. L. De La Cour, B. L. Marsh, S. F. Heron, K. V. Tirak, A. E. Strong, and C. M. Eakin. 2017. NOAA Coral Reef Watch’s 5km satellite coral bleaching heat stress monitoring product suite version 3 and four-month outlook version 4. Reef Encounter 32:39–45.

Loya, Y., K. Sakai, K. Yamazato, Y. Nakano, H. Sambali, and R. Van Woesik. 2001. Coral bleaching: the winners and the losers. Ecology Letters 4:122–131.

Lucas, J. S. 1984. Growth, maturation and effects of diet in Acanthasterplanci (L.)(Asteroidea) and hybrids reared in the laboratory. Journal of Experimental Marine Biology and Ecology 79:129–147.

MacNeil, M. A., C. Mellin, M. S. Pratchett, J. Hoey, K. R. Anthony, A. J. Cheal, I. Miller, H. Sweatman, Z. L. Cowan, S. Taylor, and others. 2016. Joint estimation of crown of thorns (Acanthaster planci) densities on the Great Barrier Reef. PeerJ 4:e2310.

MacNeil, M. A., C. Mellin, S. Matthews, N. H. Wolff, T. R. McClanahan, M. Devlin, C. Drovandi, K. Mengersen, and N. A. Graham. 2019. Water quality mediates resilience on the Great Barrier Reef. Nature Ecology & Evolution 3:620–627.

Madin, J. S., A. H. Baird, M. Dornelas, and S. R. Connolly. 2014. Mechanical vulnerability explains size-dependent mortality of reef corals. Ecology Letters 17:1008–1015.

Margvelashvili, N., J. Andrewartha, M. Baird, M. Herzfeld, E. Jones, M. Mongin, F. Rizwi, B. J. Robson, J. Skerratt, and K. Wild-Allen. 2018. Simulated fate of catchment-derived sediment on the Great Barrier Reef shelf. Marine Pollution Bulletin 135:954–962.

Massel, S. R., and T. J. Done. 1993. Effects of cyclone waves on massive coral assemblages on the Great Barrier Reef: meteorology, hydrodynamics and demography. Coral Reefs 12:153–166.

McManus, J. W., and J. F. Polsenberg. 2004. Coral–algal phase shifts on coral reefs: ecological and environmental aspects. Progress in Oceanography 60:263–279.

Mellin, C., S. Matthews, K. R. Anthony, S. C. Brown, M. J. Caley, K. A. Johns, K. Osborne, M. Puotinen, A. Thompson, and N. H. Wolff. 2019. Spatial resilience of the Great Barrier Reef under cumulative disturbance impacts. Global Change Biology 25:2431–2445.

Mellin, C., E. Peterson, M. Puotinen, and B. Schaffelke. 2020. Representation and complementarity of the long-term coral monitoring on the Great Barrier Reef. Ecological Applications in press.

Miller, I., H. Sweatman, A. Cheal, M. Emslie, K. Johns, M. Jonker, and K. Osborne. 2015. Origins and implications of a primary crown-of-thorns starfish outbreak in the southern Great Barrier Reef. Journal of Marine Biology 2015.

Moran, P. J. 1986. The Acanthaster phenomenon. Oceanography and Marine Biology: An Annual Review 24:379– 480.

Moran, P., and G. De’ath. 1992. Estimates of the abundance of the crown-of-thorns starfish Acanthaster planci in outbreaking and non-outbreaking populations on reefs within the Great Barrier Reef. Marine Biology 113:509– 515.

Moran, P. J., R. H. Bradbury, and R. E. Reichelt. 1988. Distribution of recent outbreaks of the crown-of-thorns starfish (Acanthaster planci) along the Great Barrier Reef: 1985–1986. Coral Reefs 7:125–137.

Mumby, P. J. 1999. Bleaching and hurricane disturbances to populations of coral recruits in Belize. Marine Ecology Progress Series 190:27–35.

Mumby, P. J. 2006. The impact of exploiting grazers (Scaridae) on the dynamics of Caribbean coral reefs. Ecological Applications 16:747–769.

Mumby, P. J., A. Hastings, and H. J. Edwards. 2007. Thresholds and the resilience of Caribbean coral reefs. Nature 450:98–101.

Mumby, P. J., and R. S. Steneck. 2008. Coral reef management and conservation in light of rapidly evolving ecological paradigms. Trends in Ecology & Evolution 23:555–563.

Mumby, P. J., N. H. Wolff, Y.-M. Bozec, I. Chollett, and P. Halloran. 2014. Operationalizing the resilience of coral reefs in an era of climate change. Conservation Letters 7:176–187.

Mumby, P. J., and K. Anthony. 2015. Resilience metrics to inform ecosystem management under global change with application to coral reefs. Methods in Ecology and Evolution 6:1088–1096.

Okaji, K. 1996. Feeding ecology in the early life stages of the crown-of-thorns starfish, Acanthaster planci (L.). PhD thesis, James Cook University, Australia.

Ortiz, J. C., Y.-M. Bozec, N. H. Wolff, C. Doropoulos, and P. J. Mumby. 2014. Global disparity in the ecological benefits of reducing carbon emissions for coral reefs. Nature Climate Change 4:1090.

Ortiz, J.-C., N. H. Wolff, K. R. Anthony, M. Devlin, S. Lewis, and P. J. Mumby. 2018. Impaired recovery of the Great Barrier Reef under cumulative stress. Science advances 4:eaar6127.

Osborne, K., A. M. Dolman, S. C. Burgess, and K. A. Johns. 2011. Disturbance and the dynamics of coral cover on the Great Barrier Reef (1995–2009). PLoS ONE 6:e17516.

Osborne, K., A. A. Thompson, A. J. Cheal, M. J. Emslie, K. A. Johns, M. J. Jonker, M. Logan, I. R. Miller, and H. Sweatman. 2017. Delayed coral recovery in a warming ocean. Global Change Biology 23:3869–3881.

Paine, R. T., M. J. Tegner, and E. A. Johnson. 1998. Compounded perturbations yield ecological surprises. Ecosystems 1:535–545.

Pratchett, M. S. 1999. An infectious disease in crown-of-thorns starfish on the Great Barrier Reef. Coral Reefs 18:272–272.

Pratchett, M. S. 2005. Dynamics of an outbreak population of Acanthaster planci at Lizard Island, northern Great Barrier Reef (1995–1999). Coral Reefs 24:453–462.

Pratchett, M. S. 2010. Changes in coral assemblages during an outbreak of Acanthaster planci at Lizard Island, northern Great Barrier Reef (1995–1999). Coral Reefs 29:717–725.

Pratchett, M. S., C. F. Caballes, J. A. Rivera-Posada, and H. P. Sweatman. 2014. Limits to understanding and managing outbreaks of crown-of-thorns starfish (Acanthaster spp.). Oceanography and Marine Biology: An Annual Review 52:133–200.

Pratchett, M. S., C. F. Caballes, J. C. Wilmes, S. Matthews, C. Mellin, H. Sweatman, L. E. Nadler, J. Brodie, C. A. Thompson, and J. Hoey. 2017. Thirty Years of Research on Crown-of-Thorns Starfish (1986–2016): Scientific Advances and Emerging Opportunities. Diversity 9:p 41.

Puotinen, M., J. A. Maynard, R. Beeden, B. Radford, and G. J. Williams. 2016. A robust operational model for predicting where tropical cyclone waves damage coral reefs. Scientific Reports 6:26009.

Puotinen, M., E. Drost, R. Lowe, M. Depczynski, B. Radford, A. Heyward, and J. Gilmour. 2020. Towards modelling the future risk of cyclone wave damage to the world’s coral reefs. Global Change Biology (in press).

R Core Team. 2018. R: A language and environment for statistical computing. R Foundation for statistical computing, Vienna, Austria.

Rasser, M., and B. Riegl. 2002. Holocene coral reef rubble and its binding agents. Coral Reefs 21:57–72.

Richmond, R. H. 1997. Reproduction and recruitment in corals: critical links in the persistence of reefs. Page Life and Death of Coral Reefs. Chapman & Hall, New York.

Robson, B., J. Skerratt, M. Baird, C. Davies, M. Herzfeld, E. Jones, M. Mongin, A. Richardson, F. Rizwi, K. Wild-Allen, and others. 2020. Enhanced assessment of the eReefs biogeochemical model for the Great Barrier Reef using the Concept/State/Process/System model evaluation framework. Environmental Modelling & Software:104707.

Sammarco, P. W., and J. C. Andrews. 1989. The Helix experiment: differential localized dispersal and recruitment patterns in Great Barrier Reef corals. Limnology and Oceanography 34:896–912.

Sano, M., M. Shimizu, and Y. Nose. 1987. Long-term effects of destruction of hermatypic corals by Acanthaster planci infestation on reef fish communities at Iriomote Island, Japan. Marine Ecology Progress Series:191–199.

Schaffelke, B., J. Carleton, M. Skuza, I. Zagorskis, and M. J. Furnas. 2012. Water quality in the inshore Great Barrier Reef lagoon: Implications for long-term monitoring and management. Marine Pollution Bulletin 65:249–260.

Schaffelke, B., C. Collier, F. Kroon, J. Lough, L. Mckenzie, M. Ronan, S. Uthicke, and J. Brodie. 2017. Scientific Consensus Statement 2017. A Synthesis of the Science of Land-based Water Quality Impacts on the Great Barrier Reef, Chapter 1: The Condition of Coastal and Marine Ecosystems of the Great Barrier Reef and Their Responses to Water Quality and Disturbances. State of Queensland.

Sweatman, H. H., A. A. Cheal, G. G. Coleman, M. M. Emslie, K. K. Johns, M. M. Jonker, I. I. Miller, and K. K. Osborne. 2008. Long-term Monitoring of the Great Barrier reef, Status Report 8. Australian Institute of Marine Science, Townsville, Australia.

Sweatman, H., S. Delean, and C. Syms. 2011. Assessing loss of coral cover on Australia’s Great Barrier Reef over two decades, with implications for longer-term trends. Coral Reefs 30:521–531.

Thompson, A. A., and A. M. Dolman. 2010. Coral bleaching: one disturbance too many for near-shore reefs of the Great Barrier Reef. Coral Reefs 29:637–648.

Thompson, A., T. Schroeder, V. E. Brando, and B. Schaffelke. 2014. Coral community responses to declining water quality: whitsunday Islands, Great Barrier Reef, Australia. Coral Reefs 33:923–938.

Thompson, A., P. Costello, J. Davidson, M. Logan, and G. Coleman. 2019. Marine Monitoring Program: Annual report for inshore coral reef monitoring 2017-18. Australian Institute of Marine Science: Report for the Great Barrier Reef Marine Park Authority., Great Barrier Reef Marine Park Authority, Townsville, Australia.

Trapon, M. L., M. S. Pratchett, and A. S. Hoey. 2013. Spatial variation in abundance, size and orientation of juvenile corals related to the biomass of parrotfishes on the Great Barrier Reef, Australia. PLoS ONE 8:e57788.

Vercelloni, J., K. Mengersen, F. Ruggeri, and M. J. Caley. 2017. Improved coral population estimation reveals trends at multiple scales on Australia’s Great Barrier Reef. Ecosystems 20:1337–1350.

Viehman, T. S., J. L. Hench, S. P. Griffin, A. Malhotra, K. Egan, and P. N. Halpin. 2018. Understanding differential patterns in coral reef recovery: chronic hydrodynamic disturbance as a limiting mechanism for coral colonization. Marine Ecology Progress Series 605:135–150.

Waterhouse, J., J. Brodie, D. Tracey, R. Smith, M. VanderGragt, C. Collier, C. Petus, M. Baird, F. Kroon, and R. Mann. 2017. 2017 Scientific Consensus Statement: land use impacts on the Great Barrier Reef water quality and ecosystem condition, Chapter 3: the risk from anthropogenic pollutants to Great Barrier Reef coastal and marine ecosystems.

White, J. W., A. Rassweiler, J. F. Samhouri, A. C. Stier, and C. White. 2014. Ecologists should not use statistical significance tests to interpret simulation model results. Oikos 123:385–388.

Wolfe, K., A. Graba-Landry, S. A. Dworjanyn, and M. Byrne. 2017. Superstars: Assessing nutrient thresholds for enhanced larval success of Acanthaster planci, a review of the evidence. Marine Pollution Bulletin 116:307–314.

Wolff, N. H., P. J. Mumby, M. Devlin, and K. Anthony. 2018. Vulnerability of the Great Barrier Reef to climate change and local pressures. Global Change Biology.

Zann, L., J. Brodie, C. Berryman, and M. Naqasima. 1987. Recruitment, ecology, growth and behavior of juvenile Acanthaster planci (L.) (Echinodermata: Asteroidea). Bulletin of Marine Science 41:561–575.

Zann, L., J. Brodie, and V. Vuki. 1990. History and dynamics of the crown-of-thorns starfish Acanthaster planci (L.) in the Suva area, Fiji. Coral Reefs 9:135–144.

