## Appendix S1 for "Cumulative impacts across Australia’s Great Barrier Reef: A mechanistic evaluation"

### APPENDIX S1: CORAL POPULATION MODEL

#### General description

The model represents a mid-depth (5–10 m) reef environment and simulates with a 6-month time step the settlement, growth and mortality of coral colonies and the dynamics of patches of reef algae onto a 20 × 20 grid lattice of 1 $m^2$ cells. The lattice grid has a toroidal structure (i.e., wrapped around) so that every cell has continuous boundaries formed by 4 neighboring cells. Each cell can be occupied by multiple coral colonies and patches of turf and macroalgae. Corals are stylized by the cross-sectional, basal area of a hemispherical colony ($cm^2$). A number of cells are assigned to the class "ungrazable substrate" (e.g., sand) and prevented from colonization.

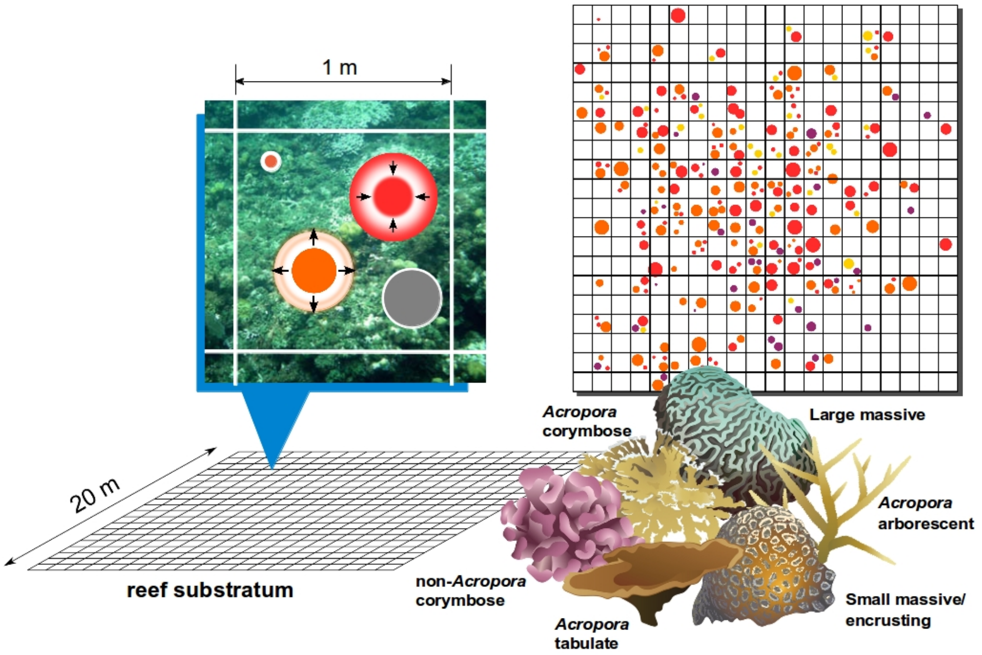


Schematic representation of a modeled mid-depth (5-10m) reef environment using a 20 m × 20 m horizontal space on which circular coral colonies can settle, grow, shrink and die. Corals are modeled by their size (planimetric area in $cm^2$) and belong to six functional groups: Arborescent (staghorn) acroporids; plating (tabular) acroporids; corymbose/small branching acroporids; pocilloporids and other non-acroporid corymbose corals; small massive/encrusting corals; large massive corals. Graphics of corals courtesy of the Integration and Application Network, University of Maryland Center for Environmental Science.

Coral demographics, ecological interactions and the impact of acute disturbances (e.g., storms, bleaching) are explicit at colony scales and follow probabilistic rules supported by empirical observations representative of a mid-depth reef environment. Corals belong to six functional groups defined as follow:

1. Arborescent (staghorn) acroporids (e.g., *Acropora muricata*, *Acropora nobilis*, *Acropora robusta*);
2. Plating acroporids (e.g., *Acropora hyacinthus*, *Acropora cytherea*);
3. Corymbose/small branching acroporids (e.g., *Acropora millepora*, *Acropora humilis*);
4. Pocilloporids/non-acroporid corymbose (e.g., *Stylophora pistillata*);
5. Small massive/submassive/encrusting corals (e.g., Lobophylliidae, favids, *Goniastrea*);
6. Large massive corals (*Porites lutea*, *Porites lobata*).

A focus on *Acropora* corals is justified as they represent the key habitat-forming species on the GBR and account for around 70% of the coral biodiversity in the Indo-Pacific region (Wallace 1999). Other model agents include patches of closely cropped algal turf (< 5mm), uncropped algal turf (> 5mm height), encrusting fleshy (i.e., *Lobophora*) and upright fleshy macroalgae. A spatially-explicit process of grazing maintains macroalgae in a cropped state, which facilitates coral settlement and growth. Grazing affects all algal groups and always results in cropped turf, on which corals can settle. Grazing is spatially limited, so that only a proportion of the grazable substrate can be efficiently grazed over 6 months. Competitive interactions between corals and macroalgae reduce the growth rate of each taxa and are the only process modeled to occur across cell boundaries (within a 4-cell von Neumann neighborhood).

At initial step, a number of cells are randomly designated as "ungrazable" until the specified cover of ungrazable substrates (i.e., sand) of the grid lattice is achieved. Grazable cells are then filled at random with coral colonies of random size generated by lognormal distributions (Meesters et al. 2001) until the sum of all colony sizes matches the specified coral cover for each species at the reef (i.e., grid) scale. Finally, algal patches are created by randomly filling the remaining space in every grazable cell until the specified algal cover is reached.

All model parameters are assumed to be constant rather than allowed to vary probabilistically. This approach avoids unnecessary variation from relatively well-established parameters. However, probabilistic rules in the spatial distribution of recruitment, natural mortality, grazing and mortality caused by disturbance events generate stochasticity in model simulations. Model outputs include the percent cover of each coral and algal species, the percent cover of loose coral rubble created by disturbances, and the density of coral colonies at different life stage and size.

#### Demographic processes

***Coral fecundity.*** Broadcast coral spawning occurs at the beginning of the wet season (austral summer). Coral fecundity $F_{coral}$, expressed as the total volume of eggs per colony, is a function of colony size (Hall and Hughes 1996). :

$$F_{coral} = \exp(a + b \times \log_e x) \qquad \text{(S1)}$$

where $x$ is the cross-sectional basal area of a colony in $cm^2$, and $a$ and $b$ are two parameters determined empirically by Hall and Hughes (1996) for some coral species (Appendix S2: Table S1). Coral colonies are considered sexually mature when their size reaches a specific threshold (Appendix 2: Table S1). Summing $F_{coral}$ over all gravid colonies with an average egg volume of 0.1 $mm^3$ (for *Acropora hyacinthus*, Hall and Hughes 1996) allows estimating a number of offspring released by each coral group during the reproductive season. The number of offspring produced on a reef is further reduced according to the concentration of suspended sediment predicted during spawning events (see below Eq. S10, S11).

***Coral recruitment.*** Corals successfully recruit to cropped turf algae (Kuffner et al. 2006, Arnold et al. 2010) at a size of 1 $cm^2$ (~6-month old corals). Recruitment is density-dependent and proportional to the space available for settlement. First, a potential density of settlers is determined from larval supply (number of incoming coral larvae from external and self-supply) following a Beverton-Holt function (main text Eq. 1). The actual density of 6-month-old recruits is generated at random for each cell given the potential density of settlers and the amount of settlement space in the cell (main text Eq. 2). Recruitment rates are further adjusted for each coral group based on empirical observations (Appendix 2: Table S1) and reduced by suspended sediments for the three acroporid groups (see below Eq. S12).

***Coral growth.*** Coral colony growth is modeled as lateral extension (radius increment) at each time step (6 month). The grow rate of coral juveniles (colony diameters ≤ 4 cm) is set to 1 cm $y^{-1}$ (Doropoulos et al. 2015, 2016), further reduced by the concentration of suspended sediments (see below Eq. S13). Adult corals grow following taxa-specific extension rates derived from literature data (Appendix 2: Table S1).

***Colonization of cropped algae.*** Cropped turf algae is the default substratum and arises (i) when long turf and macroalgae are grazed and (ii) after all coral mortality events (Jompa and McCook 2002) except those due to macroalgal overgrowth (see coral-algal competition below).

***Colonization and vegetative growth of macroalgae.*** The model simulates with a 1-month time step the spatial dynamics of macroalgal colonization following observations on 4–6m depth forereefs in Palau (Bozec et al. 2019). Diminutive algal turf is the default substrate maintained by repeated grazing. Grazing affects all algal substrates following specified feeding preferences and all consumed algal surfaces are converted into diminutive algal turf. When a cell is left ungrazed for 1 month, diminutive algal turf converts into uncropped algal turf and defines the space available for macroalgal colonization: propagules of macroalgae then settle and grow over the uncropped turf following a logistic curve. Encrusting fleshy macroaglae can also colonize from the margins of a cell due to the horizontal expansion of the surrounding vegetation.

Macroalgal settlement occurs on uncropped turf and only in cells that are left ungrazed for 1 month. Macroalgal settlement in a cell (in $cm^2$) is proportional to the area of uncropped turf in that cell, with a maximum of 2.5 $cm^2$ per $m^2$ of uncropped turf (Diaz-Pulido and McCook 2004). Then, macroalgae expand horizontally over uncropped turf following two mechanisms. First, a cell can be overgrown by the surrounding vegetation of encrusting fleshy macroalgae that expands from the neighboring cells, which is calculated as follow:

$$E_{EFM} = MO_{EFM} \times EFM_{4cell} \qquad \text{(S2)}$$

where $MO_{EFM}$ is the rate of marginal overgrowth of encrusting fleshy macroalgae per unit cover of surrounding vegetation (45 $cm^2$ per $m^2$ of uncropped turf for a 60% surrounding cover of encrusting fleshy macroalgae (De Ruyter van Steveninck and Breeman 1987) and $EFM_{4cell}$ is the proportional cover of encrusting fleshy macroalgae calculated over the 4 neighboring cells (von Neumann neighborhood). The surrounding vegetation of encrusting fleshy macroalgae is calculated before growth for every cell, and $E_{EFM}$ adds to the current amount of encrusting fleshy macroalgae only in those cells that escape grazing and where enough uncropped turf is available for macroalgal expansion. Uncropped turf is immediately reduced accordingly.

Second, each class of macroalgae (*MA*) expands horizontally over uncropped turf (*TURF*) following a logistic growth increment $Growth_{MA}$ ($cm^2$) calculated as follow:

$$Growth_{MA} = r_{MA} \times MA \times (1 - MA / (MA + TURF)) \qquad \text{(S3)}$$

where $r_{MA}$ is the intrinsic growth rate of the corresponding macroalgal group (0.598 for encrusting fleshy macroalgae; 0.843 for upright macroalgae, Bozec et al. 2019). The sum $MA + TURF$ represents the current carrying capacity of the cell for that macroalgal group. The growth of encrusting fleshy macroalgae is processed before that of upright macroalgae. The cover of uncropped turf is reduced after each macroalgal increment.

***Competition between corals.*** Coral growth is constrained by the space currently available in a cell, so that competition between corals occurs when the total area over which corals would potentially expand exceeds the amount of available space. In that case, free space is shared between all coral colonies proportionally to their growth potential. This reflects the ability of faster/larger colonies to overtake slower/smaller colonies, as growth potential depends both on extension rate and current colony size.

***Competition between corals and cropped algae.*** Corals always overgrow cropped turf algae (Jompa and McCook 2002).

***Competition between corals and macroalgae: reduction of coral growth rate.*** The growth rate of coral juveniles is set to zero if the cover of all macroalgae in the local environment formed by the focal cell and the von Neumann 4-cell neighborhood is > 80%, and reduced by 70% if total macroalgal cover lies between 40% and ≤ 80% (Box and Mumby 2007). Growth rate of larger corals is reduced by up to 90% if macroalgal cover exceeds 40% (Lirman 2001), implemented as a step function.

***Competition between corals and macroalgae: macroalgal overgrowth.*** Limited direct overgrowth of coral by macroalgae can occur. The macroalgal overgrowth of a living coral colony *i* ($O_{i \to M}$ in $cm^2$) results in partial mortality of the colony and is calculated as:

$$O_{C \to M} = MA_{5cells} \times P_i \times a_{MA,i} \qquad \text{(S4)}$$

where $MA_{5cells}$ is the proportion of encrusting fleshy or erect macroalga in the local environment formed by the focal cell and the von Neumann 4-cell neighborhood, $P_i$ is the perimeter (cm) of the coral colony *i* and $a_{MA,i}$ the average overgrowth (cm) of *i* due to the macroalga per cm length of coral edge.

In situ experiments in Curaçao, Nugues and Bak (2006) found that the average overgrowth of *Agaricia* and *Porites* corals by the encrusting fleshy macroalga *Lobophora* was 8 $cm^2$ per annum across a ~7 cm length of coral edge, which translates to $a_{L,i}$ = 0.57 $cm^2$ per cm length of coral edge in each 6 month time step of the model. Values of $a_{L,i}$ for Pacific corals were set to 0.11 $cm^2$ based on results for *Mycetophyllia alicia*e, *Meandrina meandrites* and *Porites porites.*

Lirman (2001) found no direct overgrowth of brooders by the erect macroalga *Dictyota* but an overgrowth of 0.25 – 0.43 $cm^2$ $month^{-1}$ per cm length of coral edge on the coral spawner *Orbicella faveolata*. The lowest value of this range (no herbivory exclusion) translates to $a_{D,i}$ = 1.5 cm for a 6 month time step and was applied to all corals except for pocilloporids (brooders).

***Competition between corals and macroalgae: effect of corals on macroalgae.*** The probability with which macroalgae spread vegetatively over cropped algae, $P_{A \to M}$, is reduced by 25% when at least 50% of the local von Neumann neighborhood includes coral (De Ruyter van Steveninck et al. 1988, Jompa and McCook 2002):

$$P_{A \to M} = 0.75 \times MA_{5cells}, \text{ if } C_{5cells} \geq 0.5 \qquad \text{(S5)}$$

$$P_{A \to M} = MA_{5cells}, \text{ if } C_{5cells} < 0.5 \qquad \text{(S6)}$$

where $C_{5cells}$ is the proportion of corals in the focal cell and the 4-cell von Neumann neighborhood.

***Background (chronic) whole-colony mortality of juveniles.*** Incidence of mortality in juvenile corals (diameter ≤4 cm, equivalent to an area of live cover <13 $cm^2$) is set to 0.2 $y^{-1}$ as recorded for *Acropora* spp. on a reef slope at Heron Island (Doropoulos et al. 2015). This level of mortality occur at every time step in addition to macroalgal overgrowth and mortality caused by acute disturbances.

***Background (chronic) whole-colony mortality of adults.*** Incidence of mortality of corals between 13–250 $cm^2$ is set to 0.04 $y^{-1}$ and 0.02 $y^{-1}$ for corals above 250 $cm^2$ (Bythell et al. 1993, further adjusted to each coral group, see Appendix 2: Table S1). These levels of mortality occur at every time step in addition to macroalgal overgrowth and mortality caused by acute disturbances.

***Background (chronic) partial-colony mortality of corals.*** Partial mortality is colony size-dependent, following empirical observations from Curaçao before major bleaching or hurricane disturbances (Meesters et al. 1997). State variables reported in literature were converted to dynamic variables using least squares optimization until the equilibrial state in the model matched observed data (Mumby et al. 2014) leading to the two equations below, where $P_{pm}$ is the probability of a partial mortality event, $A_{pm}$ is the area of tissue lost in a single event, and $x$ is the size (planimetric area) of the coral in cm$^2$ before shrinkage:

$$P_{pm} = 1 - [(88.9 - 11.2 \times \log_e x) \ / \ 100] \qquad \text{(S7)}$$

$$\log_e(A_{pm} \times 100) = -2.9 + 1.59 \times \log_e x \qquad \text{(S8)}$$

The extent of tissue lost ($A_{pm}$) is further adjusted to each coral group following calibration with Pacific corals (Ortiz et al. 2014, see Appendix 2: Table S1). These levels of mortality occur at every time step in addition to macroalgal overgrowth and mortality caused by acute disturbances.

***Herbivory.*** Grazing is spatially constrained (Williams et al. 2001) and is expressed as the proportion of the total reef surface efficiently maintained in a cropped state every month, which essentially represents the overall net impact of grazing resulting from the balance between continuous growth and consumption of algae at the reef scale (Mumby et al. 2007, Bozec et al. 2019). This net grazing impact (*GI*) can either be fixed, follow a stochastic stationary process or coupled with expected temporal variations in herbivorous fish abundance (Bozec et al. 2016). At each monthly time step $t$, $GI(t)$ is adjusted to the reef surface that is currently available for grazing:

$$GA(t) = GI(t) \ / \ (1 - UNGRAZ(t)) \qquad \text{(S9)}$$

where $UNGRAZ(t)$ represents the proportional area that is not available to algal colonization at time step $t$. Thus, $GA(t)$ represents the current proportion of the grazable area that is maintained in a cropped state. This essentially allows integrating the functional impact of ungrazable substrates (e.g., patches of sand, corals and other sessile invertebrates) which, by reducing the grazable area of the reef, intensify grazing on the remaining algal substrates (Williams et al. 2001). $GA(t)$ is then converted into the equivalent algal surface (cm$^2$) grazed over the reef grid. Algal removal is conducted randomly across grid cells and distributed among each algal group following specified feeding preferences (Bozec et al. 2019), which are merely rules of consumption reflecting community-wide algal selectivity of fish herbivores. Due to limited spatial data on fish and algae for the GBR, grazing across the GBR is assumed to be fully efficient (i.e., $GI$=1) in maintaining macroalgae and turf in a cropped state. All consumed algal surfaces are converted into cropped algal turf for the next model iteration.

***Impacts of suspended sediments on coral larvae, recruits and juveniles.*** Using dose-response experiments, Humanes et al. (2017a) assessed the effects of suspended sediment concentrations (*SSC)*, temperature and nutrient concentrations on the rate of fertilization of the broadcast spawning coral *Acropora tenuis*. Increasing *SSC* (5 levels: 0, 5, 10, 30, 100 mg/L) significantly affected the proportion of fertilized eggs, whereas nutrients and temperature had no or negligible impact. From the reported data (effect of each treatment averaged over 12 replicates), a dose-response curve of fertilization success to increasing *SSC* (mg/L) can be fitted to the proportion of fertilized eggs across all nutrient treatments at ambient temperature ($R^2 = 0.88$, $n = 20$ treatments, Appendix S3: Fig. S1A):

$$FERT\% = 97.417 \times \exp(-0.010 \times SSC) \qquad \text{(S10)}$$

A second experiment exposed 8-hour-old embryos of *A. tenuis* to *SSC* treatments for 28 hours (i.e., until embryos became ciliated larvae) and measured the rates of subsequent survival and settlement (Humanes et al. 2017a). While early stress exposure did not affect survival to the settlement stage, the proportion of settled larvae was significantly influenced by all stressors, with *SSC* having the strongest negative effect. The reported mean effects of *SSC* treatments at low and medium nutrient concentrations under ambient temperature (12 replicates for each treatment) can be combined to fit a dose-response curve of settlement success to increasing *SSC* in mg/L ($R^2 = 0.88$, $n = 20$ treatments, Appendix S3: Fig. S1A):

$$SETT\% = 99.571 - 10.637 \times \log(SSC + 1) \qquad \text{(S11)}$$

In another study, Humanes et al. (2017b) assessed the survival and growth of young (< 6 month-old) recruits of *Acropora millepora*, *A. tenuis* and *Pocillopora acuta* after 40 days of exposure to increased concentrations of suspended sediments (4 levels: 0, 10, 30, 100 $mg.L^{-1}$) and nutrients. The reported mean effect of *SSC* relative to the null *SSC* treatment across all nutrient treatments (3 replicate tanks for each treatment) can be combined for the two *Acropora* species to fit a dose-response curve of survival (over 40 days) to increasing *SSC* ($R^2 = 0.89$, $n = 8$ treatments, Appendix S3: Fig. S1B):

$$SURV = 1 - 1.88\text{e-}03 \times SSC \qquad \text{(S12)}$$

After conversion do daily survival rates, Eq. S12 was applied to the three acroporid groups only, since no significant effect of *SSC* was observed on the survival of *P. acuta* recruits. Practically, this amounts to reducing $D_{recruits}$ to the predicted surviving fraction.

In the same study (Humanes et al. 2017b), *SSC* was also found to reduce the growth of recruits (Appendix S3: Fig. S1C) and we assume here that a single curve fits the growth response of the three species ($R^2 = 0.79$, $n = 12$ treatments) and can be extrapolated to the growth of any coral juvenile:

$$RelG = 1 - 0.176 \times \log(SSC + 1) \qquad \text{(S13)}$$

***Cyclone impact on juvenile and adult corals: colony dislodgement.*** Storm-induced (acute) whole-colony coral mortality (i.e., dislodgement) is modeled as a function of colony size and storm strength (Mumby et al. 2007, Edwards et al. 2011). For category 5 cyclones, the probability (incidence) of whole-colony mortality $P_{wcm\_cyc5}$ was represented using a quadratic function where $x$ is the cross-sectional basal area of the colony in $cm^2$ (Bythell et al. 1993, Massel and Done 1993):

$$P_{wcm\_cyc5} = -3\text{e-}07\, x^2 + 7\text{e-}05\, x + 0.0551\, \varepsilon \qquad \text{(S14)}$$

Small colonies avoid dislodgement due to their low drag. Intermediate-sized corals have greater drag and are light enough to be dislodged, whereas large colonies are heavy enough to prevent dislodgement. A Gaussian-distributed noise $\varepsilon \sim (\mu = 0, \sigma = 0.1)$ adds variability to mortality predictions. For lower cyclone categories, this function is modified by lowering the peak by the predicted impacts of each storm category relative to the impacts of a category 5 cyclone (Edwards et al. 2011). These relative predicted impacts (category 1: 4.6%; category 2: 11.8%; category 3: 25.0%; category 4: 56.8%) were determined by a simple relationship between storm intensity (wind speed), wave height, and predicted dislodgement (Madin and Connolly 2006). Details on these calculations can be found in Edwards et al. (2011).

***Cyclone impact on mature corals (> 250 $cm^2$): partial mortality.*** The extent of partial mortality due to cyclones ($A_{pm\_cyc}$) is modeled using a Gaussian distribution with mean and standard deviation dependent on storm strength (with maximum mean of 0.30 and standard deviation of 0.20 for a category 5 cyclone, reduced as above for other cyclone categories). $A_{pm\_cyc}$ represents the percentage of original colony tissue that is lost due to the cyclone and is generated at random for every single colony. If $A_{pm\_cyc} \leq 0$, there is no partial mortality. If $A_{pm\_cyc} \geq 1$, the entire colony is lost (though this is a rare event). Data come from monitoring of impact of Hurricane Mitch in Belize (Mumby et al. 2005).

***Cyclone impact on coral recruits (1-60 $cm^2$): scouring by sand.*** Scouring by sand during a cyclone causes 80% whole-colony mortality in small corals (Mumby 1999).

***Cyclone impact on macroalgae.*** Cyclones reduce the cover of macroalgae to 10% of its pre-cyclone level (Mumby et al. 2005).

***Bleaching-induced whole-colony mortality.*** Whole-colony mortality caused my mass bleaching is a function of thermal stress (main text Eq. 3) obtained by regression of shallow (2m depth) observations of initial bleaching mortality (Hughes et al. 2018) against satellite-derived DHW (Liu et al. 2017) across the GBR during the 2016 marine heatwave. For a reef with predicted heat stress ≥ 3 DHW, the incidence of bleaching mortality is generated with a random noise (Gaussian distribution of mean 0 and variance equal to the estimated error variance of the regression model). The resulting incidence of initial mortality is adjusted to each coral group (Appendix S2: Table S1) following bleaching susceptibilities reported at the taxon level by Hughes et al. (2018), and extended to 6 months by calibration with the observed 8-months changes of coral cover following the 2016 mass beaching (see details below). Bleaching on a reef does not occur if the predicted DHW<3 or if that reef was hit by a cyclone during the same season.

***Partial-colony mortality due to bleaching.*** The incidence of partial mortality due to bleaching is equal to that of whole-colony mortality. For a coral affected by partial mortality, the extent of tissue lost (Baird and Marshall 2002) is set to 40% of the colony area for small massive/submassive (observations on *Platygyra daedalea*), 20% for large massive corals (*Porites lobota*), and a minimal 5% for the three acroporid groups (*A. hyacinthus* and *A. millepora*) extended to pocilloporids due to morphological similarities (i.e., branching corals).

### Calibration

***Calibration of parameters of coral recruitment.*** Because the processes that link larval supply and the number of 6-month old recruits are largely uncertain, recruitment parameters α and $\beta$ (main text Eq. 1) for corals were determined by calibration against GBR observations from offshore (mid- and outer-shelf) reefs. We simulated coral recovery on hypothetical reefs ($n$ = 100) and adjusted these two parameters with the constraint of reproducing simultaneously two observed data sets: 1) the recovery dynamics of coral cover following extensive coral loss (Emslie et al. 2008, Fig. 2A); 2) spatial variations in the density of coral juveniles (Trapon et al. 2013, Fig. 2B). Reefs were initialized with a random coral cover generated from a normal distribution $N(\mu = 5\%, \sigma = 0.2\times\mu)$, equally distributed among all groups. Initial proportions of loose coral rubble and sand patches were generated using a uniform distribution $U(0.10, 0.50)$. Coral connectivity and stress-induced mortality were turned off. With $\alpha$ = 15 settler/m$^2$ distributed across the six coral groups (Trapon et al. 2013, Appendix S2: Table S1) and $\beta$ = 75,000 larva/m$^2$ (among all groups), the model achieved realistic recovery rates (from ~10% to ~60% in 6-7 years) with juvenile densities within the range of empirical observations.

***Calibration of storm-induced coral mortality.*** The above calculations of cyclone impacts inherited from previous model parameterizations specific to Caribbean reefs (Mumby et al. 2007, 2014, Edwards et al. 2011, Bozec et al. 2015). To obtain more reliable predictions for GBR corals, cyclone-driven mortalities were calibrated with observations of storm damages from the benthic photo-transect survey database of the Australian Institute of Marine Science (AIMS) Long-Term Monitoring Program (LTMP). Coral cover data were extracted on reefs surveyed within one year of a designated storm exposure (pre- and post-disturbance), leading to the selection of 13 reefs (with 3 sites per reef) exposed to 4 cyclones: Justin (March 1997), Tessi (April 2000), Hamish (March 2009) and Yasi (February 2011). Cyclone exposure for each reef was reconstructed using the Database of Past Tropical Cyclone Tracks of the Australian Bureau of Meteorology (BoM). For each cyclone track (6-hour position of the storm center) a category of storm intensity defined on the Saffir-Simpson scale was assigned based on the reported value of maximum sustained winds converted into a 1-minute equivalent. Storm intensity experienced by a surveyed reef was estimated from its distance to the cyclone track, assuming asymmetric threshold distances of wind intensity from the storm center (Edwards et al. 2011). Because the spatial extent of wind intensity (and the resultant sea state that produces damaging waves) greatly varies with storm size and translation speed, threshold distances used to assign each cyclone category were further adjusted following spatial predictions of damaging sea state (average top 1/3 of wave heights [significant wave height – Hs] ≥ 4 m) relative to threshold distances for a given storm size (mean radius of gale force winds) as per Puotinen et al. 2016). As a result, the expected storm intensities fall within cyclone category 1 (*n* = 15 sites), 2 (*n* = 18) and 4 (*n* = 6). Simulations of coral damage under the expected storm intensity were run to adjust the predictions of partial- and whole-colony mortality to the observed changes in coral cover. For each surveyed reef, the model was initialized with the observed pre-disturbance cover of 37 coral taxa assigned to the modeled coral groups. Multiplying by 5 the Caribbean predictions of partial and whole-colony mortalities, further adjusted to each coral group following the response observed in the corresponding coral taxa (Appendix S2: Table S1), produced a reasonable match between the simulated and observed coral cover changes for the expected cyclone categories.

***Calibration of coral mortality due to bleaching.*** The relationship between DHW and coral mortality (main text Eq. 3) was derived from observations collected at the peak of the 2016 marine heatwave (Hughes et al. 2018). The response of corals to thermal stress over 6 months (i.e., initial + post-bleaching mortality) was determined by calibration with the observed coral cover changes (*n* = 63) sampled 8 months after the survey of initial mortality of the 2016 GBR bleaching (Hughes et al. 2018). We simulated 63 hypothetical reefs with a pre-bleaching coral cover generated at random from the frequency distribution observed in March 2016 dis-aggregated per coral type, assuming 50% acroporids. The modeled reefs were then randomly subjected to the reported DHW values (*n* = 63 DHW estimates). For each thermal stress above 3 DHW (threshold of significant mortality during the 2016 heatwave, Hughes et al. 2018), a stochastic estimate of $M_{BleachInit}$ was predicted (main text Eq. 3) with a Gaussian random noise. Extending $M_{BleachInit}$ to 6 times the period over which initial mortality was recorded (i.e., $1 - (1 - M_{BleachInit} / 100)^6$) allowed to reproduce the observed changes in coral cover 8 months after bleaching (main text. Figs. 2D-E).

### GBR observations of coral cover

The benthic survey database of the Australian Institute of Marine Science (Sweatman et al. 2008, Thompson et al. 2019) was used (1) to calibrate cyclone-driven coral mortality, (2) to initialize the 2008–2020 reconstruction of reef trajectories with realistic reef-level coral cover and (3) to validate this reconstruction.

On mid-shelf and outer reefs, monitoring surveys followed two protocols (Sweatman et al. 2008, Miller et al. 2009): (1) permanent photo transects, conducted at 6–9 m depth on three sites on the northeast flank of a reef and providing a fine-scale assessment of multiple coral taxa; (2) manta tows, conducted over the entire perimeter of a reef by a towed snorkel diver and providing a rapid assessment of coral cover on a categorical scale. Data on inshore reef (Thompson et al. 2019) were obtained from permanent photo transects conducted at 5 m depth at each of two sites.

Although the two methods produce similar assessments when applied to the same reef portion (Miller and Müller 1999), reef-wide coral cover estimates from manta tows tend to be lower due to the inclusion of non-coral habitats in the tow path (Osborne et al. 2011, Sweatman and Syms 2011). These biases were minimized by converting manta-tow cover estimates into a transect equivalent from the linear regression of 1992–2018 joint estimates from the same reef and year ($\%C_{tow} = 7.2 + 0.9 \times \%C_{transect}$, $R^2 = 0.53$, $n = 893$ joint observations).

### References


Arnold, S. N., R. S. Steneck, and P. J. Mumby. 2010. Running the gauntlet: inhibitory effects of algal turfs on the processes of coral recruitment. Marine Ecology Progress Series 414:91–105.

Baird, A., and P. Marshall. 2002. Mortality, growth and reproduction in scleractinian corals following bleaching on the Great Barrier Reef. Marine Ecology Progress Series 237:133–141.

Box, S. J., and P. J. Mumby. 2007. Effect of macroalgal competition on growth and survival of juvenile Caribbean corals. Marine Ecology Progress Series 342:139–149.

Bozec, Y.-M., L. Alvarez-Filip, and P. J. Mumby. 2015. The dynamics of architectural complexity on coral reefs under climate change. Global Change Biology 21:223–235.

Bozec, Y.-M., C. Doropoulos, G. Roff, and P. J. Mumby. 2019. Transient grazing and the dynamics of an unanticipated coral–algal phase shift. Ecosystems 22:296–311.

Bozec, Y.-M., S. O'Farrell, J. H. Bruggemann, B. E. Luckhurst, and P. J. Mumby. 2016. Tradeoffs between fisheries harvest and the resilience of coral reefs. Proceedings of the National Academy of Sciences 113:4536–4541.

Bythell, J. C., E. H. Gladfelter, and M. Bythell. 1993. Chronic and catastrophic natural mortality of three common Caribbean reef corals. Coral Reefs 12:143–152.

De Ruyter van Steveninck, E. D., and A. M. Breeman. 1987. Deep water populations of Lobophora variegata (Phaeophyceae) on the coral reef of Curaçao: influence of grazing and dispersal on distribution patterns. Marine Ecology Progress Series 38:241–250.

De Ruyter van Steveninck, E. D., L. L. Van Mulekom, and A. M. Breeman. 1988. Growth inhibition of Lobophora variegata (Lamouroux) Womersley by scleractinian corals. Journal of Experimental Marine Biology and Ecology 115:169–178.

Diaz-Pulido, G., and L. J. McCook. 2004. Effects of live coral, epilithic algal communities and substrate type on algal recruitment. Coral Reefs 23:225–233.

Doropoulos, C., G. Roff, Y.-M. Bozec, M. Zupan, J. Werminghausen, and P. J. Mumby. 2016. Characterizing the ecological trade–offs throughout the early ontogeny of coral recruitment. Ecological Monographs 86:20–44.

Doropoulos, C., S. Ward, G. Roff, M. González-Rivero, and P. J. Mumby. 2015. Linking demographic processes of juvenile corals to benthic recovery trajectories in two common reef habitats. PLoS ONE 10:e0128535.

Edwards, H. J., I. A. Elliott, C. M. Eakin, A. Irikawa, J. S. Madin, M. McField, J. A. Morgan, R. van Woesik, and P. J. Mumby. 2011. How much time can herbivore protection buy for coral reefs under realistic regimes of hurricanes and coral bleaching? Global Change Biology 17:2033–2048.

Hall, V., and T. Hughes. 1996. Reproductive strategies of modular organisms: comparative studies of reef-building corals. Ecology 77:950–963.

Hughes, T. P., J. T. Kerry, A. H. Baird, S. R. Connolly, A. Dietzel, C. M. Eakin, S. F. Heron, A. S. Hoey, M. O. Hoogenboom, G. Liu, and others. 2018. Global warming transforms coral reef assemblages. Nature 556:492.

Humanes, A., G. F. Ricardo, B. L. Willis, K. E. Fabricius, and A. P. Negri. 2017a. Cumulative effects of suspended sediments, organic nutrients and temperature stress on early life history stages of the coral Acropora tenuis. Scientific Reports 7:44101.

Humanes, A., A. Fink, B. L. Willis, K. E. Fabricius, D. de Beer, and A. P. Negri. 2017b. Effects of suspended sediments and nutrient enrichment on juvenile corals. Marine Pollution Bulletin 125:166–175.

Jompa, J., and L. J. McCook. 2002. Effects of competition and herbivory on interactions between a hard coral and a brown alga. Journal of Experimental Marine Biology and Ecology 271:25–39.

Kuffner, I. B., L. J. Walters, M. A. Becerro, V. J. Paul, R. Ritson-Williams, and K. S. Beach. 2006. Inhibition of coral recruitment by macroalgae and cyanobacteria. Marine Ecology Progress Series 323:107–117.

Lirman, D. 2001. Competition between macroalgae and corals: effects of herbivore exclusion and increased algal biomass on coral survivorship and growth. Coral Reefs 19:392–399.

Liu, G., W. J. Skirving, E. F. Geiger, J. L. De La Cour, B. L. Marsh, S. F. Heron, K. V. Tirak, A. E. Strong, and C. M. Eakin. 2017. NOAA Coral Reef Watch's 5km satellite coral bleaching heat stress monitoring product suite version 3 and four-month outlook version 4. Reef Encounter 32:39–45.

Madin, J. S., and S. R. Connolly. 2006. Ecological consequences of major hydrodynamic disturbances on coral reefs. Nature 444:477–480.

Massel, S. R., and T. J. Done. 1993. Effects of cyclone waves on massive coral assemblages on the Great Barrier Reef: meteorology, hydrodynamics and demography. Coral Reefs 12:153–166.

Meesters, E. H., M. Hilterman, E. Kardinaal, M. Keetman, M. deVries, and R. P. M. Bak. 2001. Colony size-frequency distributions of scleractinian coral populations: spatial and interspecific variation. Marine Ecology Progress Series 209:43–54.

Meesters, E. H., I. Wesseling, and R. P. Bak. 1997. Coral colony tissue damage in six species of reef-building corals: partial mortality in relation with depth and surface area. Journal of Sea Research 37:131–144.

Miller, I., M. Jonker, and G. Coleman. 2009. Crown-of-thorns starfish and coral surveys using the manta tow and SCUBA search techniques. Australian Institute of Marine Science, Townsville, Australia.

Miller, I., and R. Müller. 1999. Validity and reproducibility of benthic cover estimates made during broadscale surveys of coral reefs by manta tow. Coral Reefs 18:353–356.

Mumby, P. J. 1999. Bleaching and hurricane disturbances to populations of coral recruits in Belize. Marine Ecology Progress Series 190:27–35.

Mumby, P. J., N. L. Foster, and E. A. G. Fahy. 2005. Patch dynamics of coral reef macroalgae under chronic and acute disturbance. Coral Reefs 24:681–692.

Mumby, P. J., A. Hastings, and H. J. Edwards. 2007. Thresholds and the resilience of Caribbean coral reefs. Nature 450:98–101.

Mumby, P. J., N. H. Wolff, Y.-M. Bozec, I. Chollett, and P. Halloran. 2014. Operationalizing the resilience of coral reefs in an era of climate change. Conservation Letters 7:176–187.

Nugues, M. M., and R. P. Bak. 2006. Differential competitive abilities between Caribbean coral species and a brown alga: a year of experiments and a long-term perspective. Marine Ecology Progress Series 315:75–86.

Ortiz, J. C., Y.-M. Bozec, N. H. Wolff, C. Doropoulos, and P. J. Mumby. 2014. Global disparity in the ecological benefits of reducing carbon emissions for coral reefs. Nature Climate Change 4:1090.

Osborne, K., A. M. Dolman, S. C. Burgess, and K. A. Johns. 2011. Disturbance and the dynamics of coral cover on the Great Barrier Reef (1995–2009). PLoS ONE 6:e17516.

Puotinen, M., J. A. Maynard, R. Beeden, B. Radford, and G. J. Williams. 2016. A robust operational model for predicting where tropical cyclone waves damage coral reefs. Scientific Reports 6:26009.

Sweatman, H. H., A. A. Cheal, G. G. Coleman, M. M. Emslie, K. K. Johns, M. M. Jonker, I. I. Miller, and K. K. Osborne. 2008. Long-term Monitoring of the Great Barrier reef, Status Report 8. Australian Institute of Marine Science, Townsville, Australia.

Sweatman, H., and C. Syms. 2011. Assessing loss of coral cover on the Great Barrier Reef: A response to Hughes et al.(2011). Coral Reefs 30:661.

Thompson, A., P. Costello, J. Davidson, M. Logan, and G. Coleman. 2019. Marine Monitoring Program: Annual report for inshore coral reef monitoring 2017-18.

Wallace, C. 1999. Staghorn corals of the world: a revision of the genus Acropora. CSIRO publishing.

Williams, I. D., N. V. Polunin, and V. J. Hendrick. 2001. Limits to grazing by herbivorous fishes and the impact of low coral cover on macroalgal abundance on a coral reef in Belize. Marine Ecology Progress Series 222:187–196.
