## Appendix S2 for "Cumulative impacts across Australia’s Great Barrier Reef: A mechanistic evaluation"

### APPENDIX S2: SUPPLEMENTARY TABLES

**Table S1.** Coral demographic parameters of ReefMod-GBR.

| | Arborescent (staghorn) acroporids | Plating acroporids | Corymbose / small branching acroporids | Pocilloporids / non-acroporid corymbose | Small massive / submassive / encrusting | Large massive |
|---|---|---|---|---|---|---|
| Representative coral taxa | *Acropora muricata, Acropora nobilis, Acropora robusta* | *Acropora hyacinthus, Acropora cytherea* | *Acropora millepora, Acropora humilis* | *Stylophora pistillata, Pocillopora damicornis, Seriatopora hystrix* | *Lobophylliidae, favids, Goniastrea* | *Porites lutea, Porites lobata* |
| Radial extension of adult colonies (cm $yr^{-1}$) | 4.4 [a] | 4.4 [b] | 3.0 [c] | 2.4 [d] | 0.8 [e] | 1.2 [f] |
| Radial extension of juveniles (cm $yr^{-1}$) [g] | 1.0 | 1.0 | 1.0 | 1.0 | 1.0 | 1.0 |
| Fecundity parameter *a* [h] | 1.03 | 1.03 | 1.69 | -1.20 | 0.86 | 0.86 |
| Fecundity parameter *b* [h] | 1.28 | 1.28 | 1.05 | 2.27 | 1.21 | 1.21 |
| Minimum size of full fecundity ($cm^2$) [h] | 123 | 123 | 134 | 31 | 38 | 38 |
| Maximum settler density $\alpha$ ($m^{-2}$) [i] | 0.5 | 2.5 | 2.5 | 1.5 | 1.5 | 1.5 |
| Larval density for $\alpha/2$ ($m^{-2}$) [i] | 5,000 | 5,000 | 5,000 | 5,000 | 5,000 | 5,000 |
| Juvenile mortality ($yr^{-1}$) [j] | 0.2 | 0.2 | 0.2 | 0.2 | 0.2 | 0.2 |
| Whole-colony mortality of small (< 250 $cm^2$) corals ($yr^{-1}$) [k] | 0.004 | 0.190 | 0.172 | 0.226 | 0.040 | 0.040 |
| Whole-colony mortality of large (≥ 250 $cm^2$) corals ($yr^{-1}$) [k] | 0.002 | 0.098 | 0.088 | 0.116 | 0.020 | 0.020 |
| Relative sensitivity to partial mortality [l] | 2.0 | 0.4 | 1.4 | 1.8 | 1.0 | 1.0 |
| Relative sensitivity to | 1.4 | 1.6 | 1.5 | 1.1 | 0.6 | 0.5 |

| | Arborescent (staghorn) acroporids | Plating acroporids | Corymbose / small branching acroporids | Pocilloporids / non-acroporid corymbose | Small massive / submassive / encrusting | Large massive |
|---|---|---|---|---|---|---|
| cyclones [m] | | | | | | |
| Relative sensitivity to bleaching [n] | 1.5 | 1.6 | 1.4 | 1.7 | 0.25 | 0.25 |

[a] Horizontal projection of branch extension 8.2 $cm.yr^{-1}$ × cosinus (45°): Crossland 1981, Oliver et al. 1983, Dennison and Barnes 1988, Simpson 1988, Harriott 1998, Anderson et al. 2017.

[b] Stimson 1985, Jokiel and Tyler 1992.

[c] Stimson 1985, Jokiel and Tyler 1992, Clark and Edwards 1995, Morgan and Kench 2012.

[d] Crossland 1981, Oliver 1985, Stimson 1985, Simpson 1988, Jokiel and Tyler 1992, Harriott 1999, Anderson et al. 2012.

[e] Knutson et al. 1972, Buddemeier 1974, Highsmith 1979, Babcock 1991, Clark and Edwards 1995.

[f] Knutson et al. 1972, Buddemeier 1974, Highsmith 1979, Barnes and Lough 1989, Jokiel and Tyler 1992, Clark and Edwards 1995, Morgan and Kench 2012.

[g] Extension rate of Pocilloporid and Acroporid juveniles (1 mm diameter) on a reef slope (≤ 5 m depth) at Heron Island (Doropoulos et al. 2015).

[h] Hall and Hughes 1996.

[i] Determined by calibration against observations of coral juvenile densities (Trapon et al. 2013) and rates of coral recovery (Emslie et al. 2008); assumes the juvenile taxonomic composition observed by Trapon et al. (2013): *Acropora* ~40-60%, *Pocillopora* ~5-15%, *Porites* ~15-30%, others ~10-20%.

[j] Mortality of juveniles (≤ 5 cm maximum diameter) of *Acropora* spp. on a reef slope (≤ 5 m depth) at Heron Island (Doropoulos et al. 2015).

[k] Chronic (background) whole-colony mortality rates based on *Porites astreoides* in the Caribbean (Bythell et al. 1993, Mumby and Dytham 2006) further adjusted to each coral group following calibration with Pacific corals (Ortiz et al. 2014).

[l] Coefficients to adjust the extent of tissue lost (partial mortality) estimated from colony size (empirical relationship based on 6 Caribbean species, Meesters et al. 1997, detailed in Appendix S1). Coefficients were determined by calibration with Pacific corals (Ortiz et al. 2014).

[m] Based on taxa-specific responses to storms of different categories assessed using the AIMS LTMP transect surveys database: cover change of each taxa relative to total cover change after cyclone exposure. Multiplied by 6 after calibration with the observed changes in coral cover.

[n] Based on (Hughes et al. 2018), taking as baseline an average bleaching mortality of ~20% across taxa.

**Table S2.** Litterature data on mortality rates from manipulative in situ experiments (Keesing and Lucas 1992, Sweatman 1995, Okaji 1996, Keesing et al. 1996, 2018) and monitoring of CoTS cohorts (Zann et al. 1987, 1990). Daily mortality rates were estimated from the reported surviving fractions and the duration of each study. Median ages are the representative age of the surviving fractions, extrapolated from initial age and study duration (except for Sweatman 1995), and subsequently used as point estimates for the empirical model of mortality at age.

| Location | Initial age (months) | Initial size range or mean (mm) | Study duration (days) | Median age (months) | Surviving fraction (%) | Daily mortality (%.$d^{-1}$) | Reference |
|---|---|---|---|---|---|---|---|
| Suva Reef, Fiji | 8–23 | NA | 487 | 16 | 0.7 [c] | 1.01 | [1] (Zann et al. 1987) |
| GBR mid-shelf, various habitats | 22–34 | NA | 365 | 28.0 | 23.7 | 0.39 | [2] (Doherty and Davidson 1988) |
| Suva Reef, Fiji | 22–28 | NA | 182 | 25 | 27.5 [d] | 0.71 | [3] (Zann et al. 1990) |
| Davies Reef (12-15m) | 1 | 0.6–1.7 (1.0) | 5.75 | 1.1 | 62 | 7.98 | [4] (Keesing and Halford 1992) |
| Davies Reef (12-15m) | 4 | 0.7–4.7 (2.8) | 13 | 4.2 | 85 | 1.24 | [4] (Keesing and Halford 1992) |
| Davies Reef (12-15m) | 7 | 1.4–6.0 [a] (4.9) | 16 | 7.3 | 94 | 0.54 | [4] (Keesing and Halford 1992) |
| Davies Reef back reef (3-9m) | NA | 15–79 | 35 | ~14.6 [b] | NA | 0.41 | [5] (Sweatman 1995) |
| Davies Reef (12-15m) | 1 | 1.0–1.6 | 9 | 1.1 | NA | 7.80 | [6] (Keesing et al. 1996) |
| Davies Reef leeward slope (10m) | 0.3 | 0.6 | 49 | 1.1 | 1 | 8.97 | [7] (Okaji 1996) |
| Davies Reef leeward slope (10m) | 0.3 | 0.7 | 37 | 0.9 | 14.7 | 5.05 | [7] (Okaji 1996) |
| Davies Reef leeward slope (10m) | 2 | 1.2 | 57 | 2.9 | 22.6 | 2.57 | [7] (Okaji 1996) |
| Davies Reef leeward slope (10m) | 3 | 3.2 | 92 | 4.5 | 38.7 | 2.40 | [7] (Okaji 1996) |
| Davies Reef (10m) | 3 | 3 | 17 | 3.3 | 60.4 | 2.92 | [8] (Keesing et al. 2018) |
| Davies Reef (10m) | 15 | 13 | 17 | 15.3 | 82.5 | 1.13 | [8] (Keesing et al. 2018) |

[a] 7mo-old starfish stunted due to lack of food (expected size at this age is 12-28mm, Zann et al. 1990).

[b] Median age extrapolated from size-at-age data (Zann et al. 1987).

[c] Mass mortality due to disease.

[d] Uncertain estimates of abundance and aggregative CoTS distribution.

**Table S3.** Coral cover changes across the GBR and shelf from 2008 to 2020 (13 years).

| | Initial coral cover (%cover) | Absolute change (%cover) | Proportional change |
|---|---|---|---|
| **GBR** | | | |
| Cross-shelf | 28.8 | –9.6 | –33% |
| Inner-shelf | 33.4 | –19.0 | –57% |
| Mid-shef | 27.5 | –5.2 | –19% |
| Outer-shelf | 27.4 | –11.2 | –41% |
| **Northern GBR** | | | |
| Cross-shelf | 28.3 | –15.2 | –54% |
| Inner-shelf | 31.7 | –20.2 | –64% |
| Mid-shef | 28.4 | –15.1 | –53% |
| Outer-shelf | 24.8 | –10.4 | –42% |
| **Central GBR** | | | |
| Cross-shelf | 22.9 | –2.9 | –13% |
| Inner-shelf | 30.4 | –15.9 | –52% |
| Mid-shef | 21.5 | +1.7 | +8% |
| Outer-shelf | 21.7 | –7.6 | –35% |
| **Southern GBR** | | | |
| Cross-shelf | 33.1 | –8.6 | –26% |
| Inner-shelf | 36.6 | –19.0 | –52% |
| Mid-shef | 30.9 | –2.7 | –9% |
| Outer-shelf | 36.1 | –15.5 | –43% |

**Table S4.** Impacts of significant cyclones during 2008–2020 as absolute (% cover) and proportional (in parenthesis) coral cover loss (ie, relative to pre-cyclone total coral cover) averaged by region. Cyclone impacts result from reef-level predictions cyclone severity (Saffir-Simpson category) and simulated coral community composition.

| Cyclone | GBR | North | Central | South |
|---|---|---|---|---|
| Hamish (2009) | –9.9 (–30%) | | –5.0 (–19%) | –22.5 (–65%) |
| Olga (2010) | –0.4 (–2%) | | –1.0 (–4%) | –0.2 (–2%) |
| Yasi (2011) | –1.1 (–6%) | | –4.4 (–23%) | |
| Tim (2013) | –2.3 (–10%) | –0.8 (–2%) | –3.0 (–15%) | –3.0 (–14%) |
| Ita/Dylan (2014) | –2.2 (–8%) | –2.6 (–8%) | –3.9 (–16%) | –0.7 (–3.5%) |
| Nathan/Marcia (2015) | –3.7 (–14%) | –3.0 (–10%) | | –6.8 (–26%) |
| Debbie (2017) | –2.6 (–9%) | | –8.1 (–24%) | –1.5 (–8%) |
| Iris (2018) | –0.6 (–3%) | | –1.8 (–9%) | –0.4 (–3%) |
| Trevor (2019) | –0.4 (–3%) | –0.8 (–5%) | | |

**Table S5.** Variance component analysis of coral recovery. Relative contribution of simulated environmental variables to the variations of coral community growth ($g$) measured as the percent deviance explained by separate GLMs with a single predictor, and by a global GLM with all predictors included. Code for environmental variables: *Coral* – coral cover; *Sand* – cover of sand; *Rubble* – cover of loose coral rubble; $WQ_{repro}$/$WQ_{recruit}$/$WQ_{juv}$ – sediment-driven potential of coral reproduction/recruit survival/juvenile; $Supply_{prop}$ – relative proportion of external larval supply; $Supply_{num}$ – number of external supply links. Final regression model (*R* software pseudo-syntax): $g$ + 2 = poly(*Coral*, 2) : ( asin($WQ_{juv}$) + asin($WQ_{recruit}$) + log(1+*Rubble*) + *Sand* + sqrt($Supply_{num}$) + sqrt($Supply_{prop}$) + asin($WQ_{repro}$) ), family = Gamma(link = log).

| GLMs | % deviance explained (all reefs) | % deviance explained (inshore reefs) |
|---|---|---|
| *Coral* | 25.0 | 5.7 |
| *Sand* | 3.2 | 4.2 |
| $WQ_{recruit}$ | 0.7 | 3.8 |
| $WQ_{juv}$ | 0.5 | 2.5 |
| $WQ_{repro}$ | 0.5 | 2.1 |
| $Supply_{prop}$ | 0.1 | 0.2 |
| $Supply_{num}$ | 0.0 | 0.0 |
| *Rubble* | 0.0 | 0.2 |
| *Coral* : ($WQ_{juv}$ + $WQ_{recruit}$ + *Rubble* + *Sand* + $Supply_{num}$ + $Supply_{prop}$ + $WQ_{repro}$) | 35.6 | 18.4 |

**Table S6.** Distribution of coral health status predicted in 2020 (post-bleaching) among reefs of the entire GBR (*n*=3,806 reefs) and in each region region (North: *n*=1,200; Central: *n*=958; South: *n*=1,648).

| Reef status | GBR | North | Central | South |
|---|---|---|---|---|
| Critical (<10%) | 22% | 41% | 21% | 9% |
| Poor (10–20%) | 41% | 44% | 46% | 38% |
| Moderate (20–30%) | 18% | 9% | 18% | 24% |
| Healthy (>30%) | 19% | 6% | 15% | 29% |

### REFERENCES


Anderson, K. D., N. E. Cantin, S. F. Heron, C. Pisapia, and M. S. Pratchett. 2017. Variation in growth rates of branching corals along Australia's Great Barrier Reef. Scientific reports 7:1–13.

Anderson, K., M. Pratchett, and A. Baird. 2012. Summer growth rates of corals at Lord Howe Island, Australia.

Babcock, R. C. 1991. Comparative demography of three species of scleractinian corals using age□and size□dependent classifications. Ecological Monographs 61:225–244.

Barnes, D. J., and J. M. Lough. 1989. The nature of skeletal density banding in scleractinian corals: fine banding and seasonal patterns. Journal of Experimental Marine Biology and Ecology 126:119–134.

Buddemeier, R. W. 1974. Environmental controls over annual and lunar monthly cycles in hermatypic coral calcification. Pages 259–267 Proc 2nd Int Coral Reef Symp.

Bythell, J. C., E. H. Gladfelter, and M. Bythell. 1993. Chronic and catastrophic natural mortality of three common Caribbean reef corals. Coral Reefs 12:143–152.

Clark, S., and A. J. Edwards. 1995. Coral transplantation as an aid to reef rehabilitation: evaluation of a case study in the Maldive Islands. Coral reefs 14:201–213.

Crossland, C. J. 1981. Seasonal growth of Acropora cf. formosa and Pocillopora damicornis on a high latitude reef (Houtman Abrolhos, Western Australia). Proceedings of the Fourth International Coral Reef Symposium:663–667.

Dennison, W. C., and D. J. Barnes. 1988. Effect of water motion on coral photosynthesis and calcification. Journal of Experimental Marine Biology and Ecology 115:67–77.

Doherty, P. P., and J. J. Davidson. 1988. Monitoring the distribution and abundance of juvenile Acanthaster planci in the central Great Barrier Reef. Page Proceedings of the 6th International Coral Reef Symposium, Townsville, Australia 8-12 August 1988-pages: 2: 131-136.

Doropoulos, C., S. Ward, G. Roff, M. González-Rivero, and P. J. Mumby. 2015. Linking demographic processes of juvenile corals to benthic recovery trajectories in two common reef habitats. PLoS ONE 10:e0128535.

Emslie, M., A. Cheal, H. Sweatman, and S. Delean. 2008. Recovery from disturbance of coral and reef fish communities on the Great Barrier Reef, Australia. Marine Ecology Progress Series 371:177–190.

Hall, V., and T. Hughes. 1996. Reproductive strategies of modular organisms: comparative studies of reef-building corals. Ecology 77:950–963.

Harriott, V. J. 1998. Growth of the staghorn coral Acropora formosa at Houtman Abrolhos, Western Australia. Marine Biology 132:319–325.

Harriott, V. J. 1999. Coral growth in subtropical eastern Australia. Coral Reefs 18:281–291.

Highsmith, R. C. 1979. Coral growth rates and environmental control of density banding. Journal of Experimental Marine Biology and Ecology 37:105–125.

Hughes, T. P., J. T. Kerry, A. H. Baird, S. R. Connolly, A. Dietzel, C. M. Eakin, S. F. Heron, A. S.

Hoey, M. O. Hoogenboom, G. Liu, and others. 2018. Global warming transforms coral reef assemblages. Nature 556:492.

Jokiel, P. L., and W. A. Tyler. 1992. Distribution of stony corals in Johnston Atoll lagoon. Pages 683–692 Proceedings of the Seventh International Coral Reef Symposium.

Keesing, J. K., and A. R. Halford. 1992. Importance of postsettlement processes for the population dynamics of Acanthaster planci (L.). Marine and Freshwater Research 43:635–651.

Keesing, J. K., A. R. Halford, and K. C. Hall. 2018. Mortality rates of small juvenile crown-of-thorns starfish Acanthaster planci on the Great Barrier Reef: implications for population size and larval settlement thresholds for outbreaks. Marine Ecology Progress Series 597:179–190.

Keesing, J. K., W. L. Wiedermeyer, K. Okaji, A. R. Halford, K. C. Hall, and C. M. Cartwright. 1996. Mortality rates of juvenile starfish Acanthaster planci and Nardoa spp measured on the Great Barrier Reef, Australia and in Okinawa, Japan. Oceanologica acta 19:441–448.

Knutson, D. W., R. W. Buddemeier, and S. V. Smith. 1972. Coral chronometers: seasonal growth bands in reef corals. Science 177:270–272.

Meesters, E. H., I. Wesseling, and R. P. Bak. 1997. Coral colony tissue damage in six species of reef-building corals: partial mortality in relation with depth and surface area. Journal of Sea Research 37:131–144.

Morgan, K. M., and P. S. Kench. 2012. Skeletal extension and calcification of reef-building corals in the central Indian Ocean. Marine environmental research 81:78–82.

Mumby, P. J., and C. Dytham. 2006. Metapopulation dynamics of hard corals. Pages 157–203 Marine metapopulations. Elsevier.

Okaji, K. 1996. Feeding ecology in the early life stages of the crown-of-thorns starfish, Acanthaster planci (L.). PhD thesis, James Cook University, Australia.

Oliver, J. 1985. An evaluation of the biological and economic aspects of commercial coral collecting in the Great Barrier Reef Region. Report of the Great Barrier Reef Marine Park Authority, Townsville, Australia 106.

Oliver, J. K., B. E. Chalker, and W. C. Dunlap. 1983. Bathymetric adaptations of reef-building corals at Davies Reef, Great Barrier Reef, Australia. I. Long-term growth responses of Acropora formosa (Dana 1846). Journal of Experimental Marine Biology and Ecology 73:11–35.

Ortiz, J. C., Y.-M. Bozec, N. H. Wolff, C. Doropoulos, and P. J. Mumby. 2014. Global disparity in the ecological benefits of reducing carbon emissions for coral reefs. Nature Climate Change 4:1090.

Simpson, C. J. 1988. Ecology of Scleractinian corals in the Dampier Archipelago. Western Australia Technical Series. Environmental Protection Authority, Perth.

Stimson, J. 1985. The effect of shading by the table coral Acropora hyacinthus on understory corals. Ecology 66:40–53.

Sweatman, H. P. A. 1995. A field study of fish predation on juvenile crown-of-thorns starfish. Coral Reefs 14:47–53.

Trapon, M. L., M. S. Pratchett, and A. S. Hoey. 2013. Spatial variation in abundance, size and orientation of juvenile corals related to the biomass of parrotfishes on the Great Barrier Reef, Australia. PLoS ONE 8:e57788.

Zann, L., J. Brodie, C. Berryman, and M. Naqasima. 1987. Recruitment, ecology, growth and behavior of juvenile Acanthaster planci (L.) (Echinodermata: Asteroidea). Bulletin of Marine Science 41:561–575.

Zann, L., J. Brodie, and V. Vuki. 1990. History and dynamics of the crown-of-thorns starfish Acanthaster planci (L.) in the Suva area, Fiji. Coral Reefs 9:135–144.
