## Appendix S3 for "Cumulative impacts across Australia’s Great Barrier Reef: A mechanistic evaluation"

### APPENDIX S3: SUPPLEMENTARY FIGURES

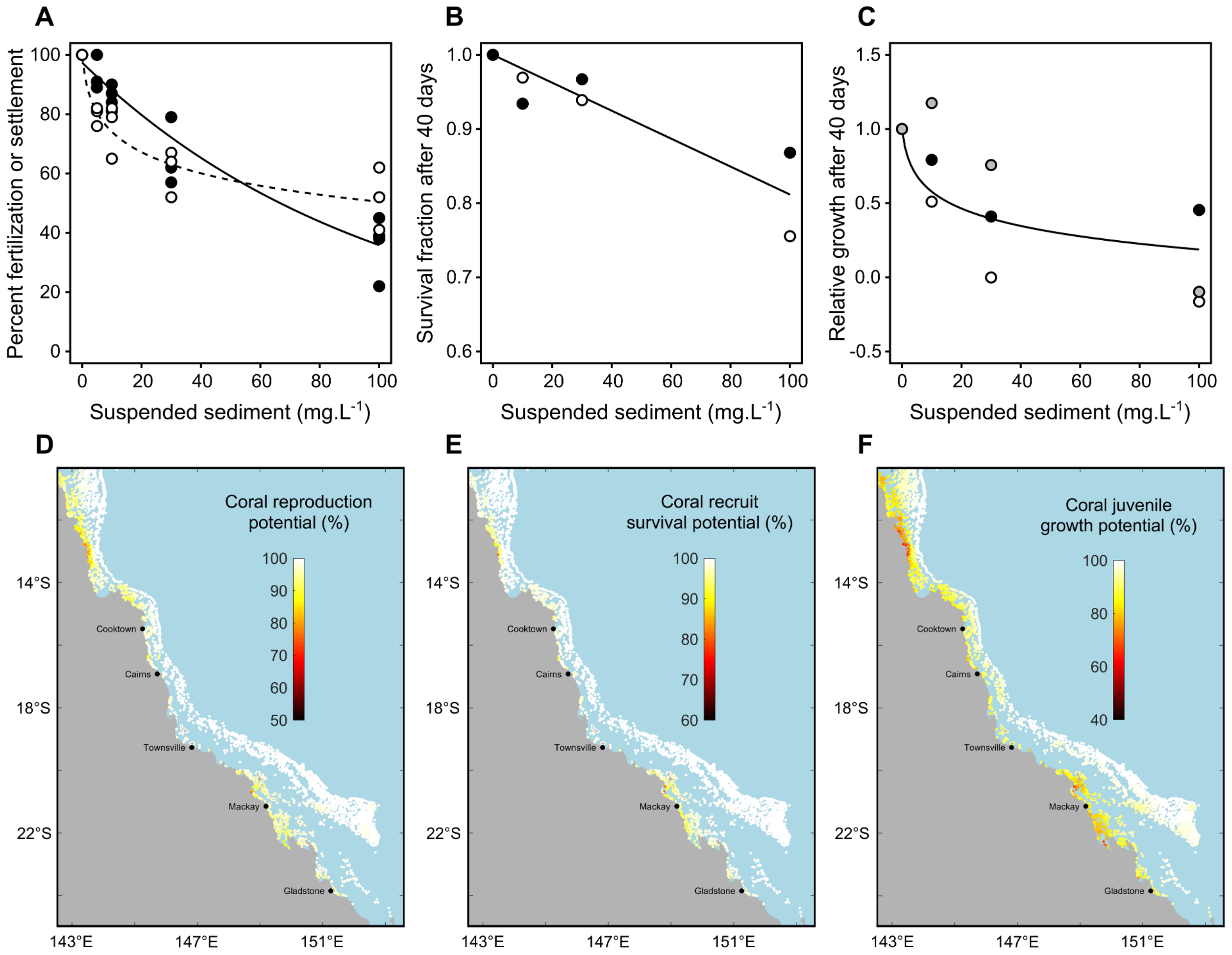


**Fig. S1**. GBR predictions of coral pre-settlement and post-recruitment success from spatio-temporal distributions of suspended sediment concentrations (SSC) and experimental dose-response relationships (Humanes et al. 2017a, b). **(A)** Relative fertilization (*FERT*%, filled circles, *n* = 20 treatments) and settlement (*SETT*%, open circles, *n* = 20 treatments) success following exposure of embryos to increasing SSC, fitted with linear models. Solid line: model for *FERT*% ($R^2$ = 0.88, Appendix S1: Eq. S10); dashed line: model for *SETT*% ($R^2$ = 0.88, Appendix S1: Eq. S10). **(B)** Relative survival (*SURV*, survived fraction relative to control) of 3-6 month old recruits after 40 days of SSC exposure (filled circles: *Acropora tenuis*; open circles: *Acropora millepora; n* = 8 treatments) fitted with a linear model ($R^2$ = 0.89, Appendix S1: Eq. S12). **(C)** Proportional growth of coral recruits relative to control after 40 days of SSC exposure (*RelG*; black-filled circles: *A. tenuis*; open circles: *A. millepora*; gray-filled circles: *Pocillopora acuta; n* = 12 treatments) fitted with a linear model ($R^2$ = 0.79, Appendix S1: Eq. S13). GBR year-averaged (geometric mean) predictions of **(D)** coral reproduction potential (*FERT*% × *SETT*%), **(E)** survival of 6-month old acroporid recruits and **(F)** relative growth of juveniles from daily SSC predictions of eReefs at 4 km resolution (GBR4_H2p0_B3p1_Cq3b) assigned to the nearest reef polygon. See Figs. S3, S5-S7 for annual maps.

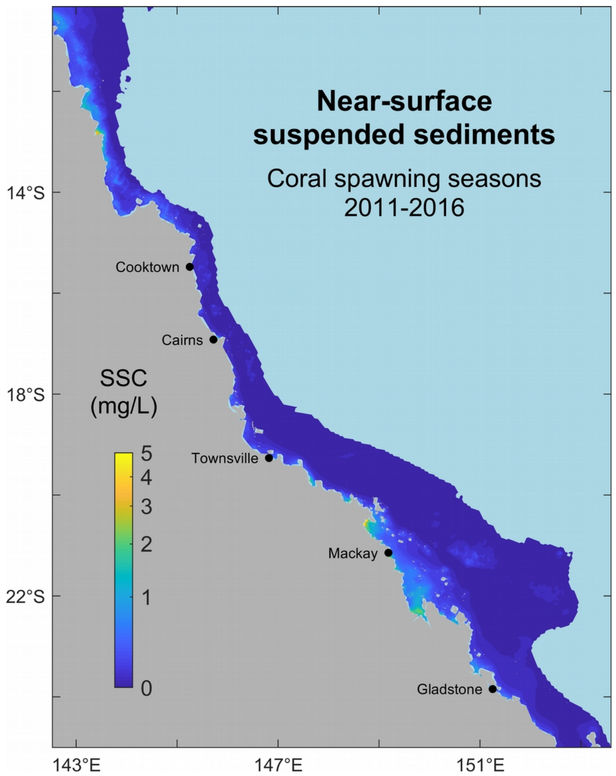


**Fig. S2.** Average predictions of near-surface (-0.5 m) suspended sediment concentrations (SSC, on a logarithmic scale) during coral spawning (reproductive seasons 2011–2016) at 4km resolution from eReefs. Predictions were based on daily SSC values starting from and extending to 3 days after the observed/expected dates of spawning across the GBR.

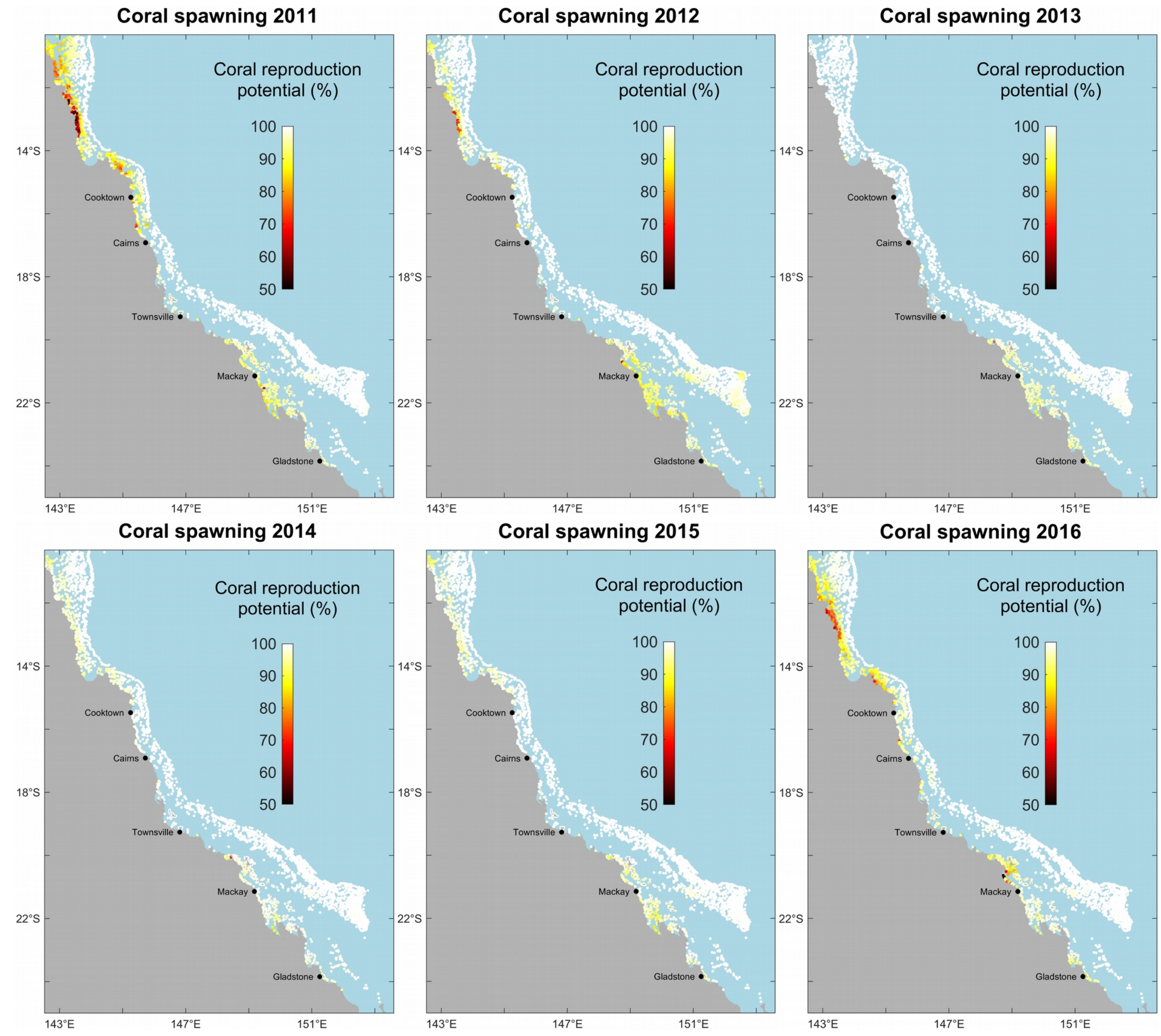


**Fig. S3.** GBR 2011–2016 predictions of the reproduction potential of corals as a function of concentrations of suspended sediments predicted by eReefs during coral spawning.

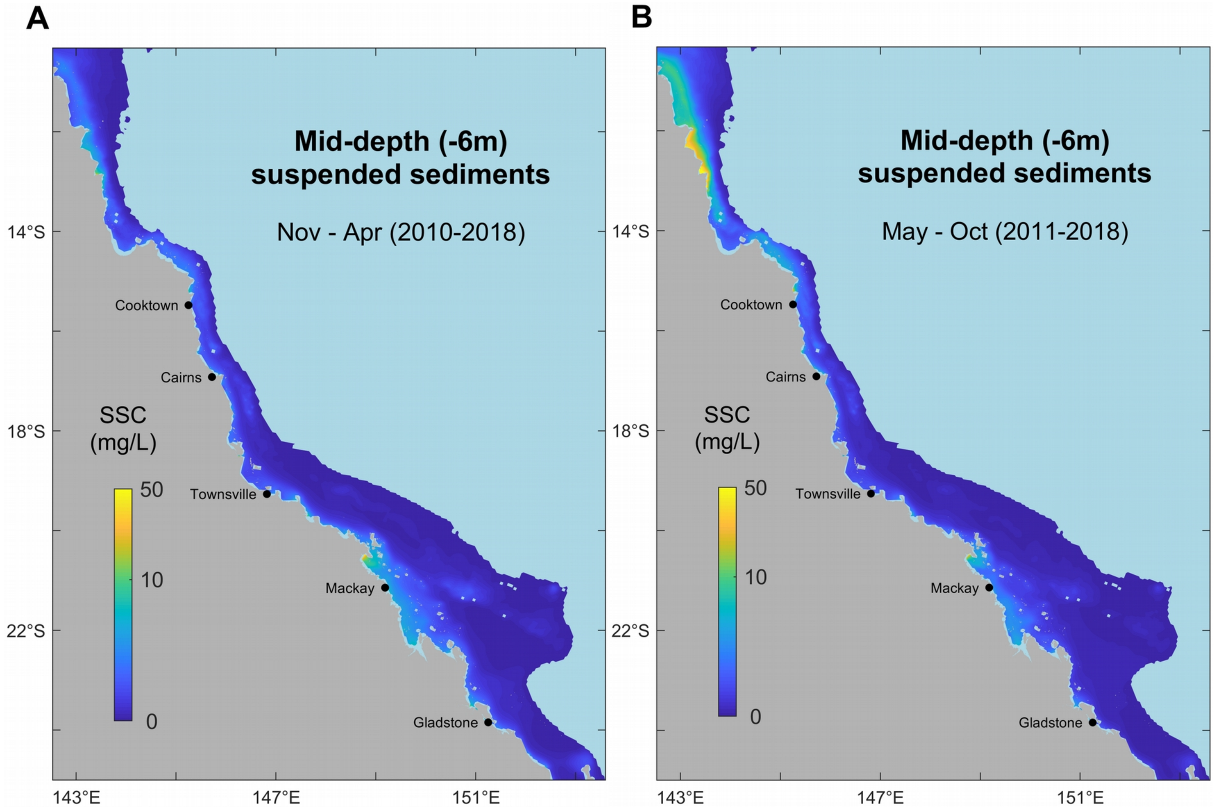


**Fig. S4.** Average summer (wet season, **A**) and winter (dry season, **B**) predictions of mid-depth (~ -6 m) suspended sediment concentrations (SSC, on a logarithmic scale) at 4km resolution from the eReefs hydrodynamic/biogeochemical model GBR4_H2p0_B3p1_Cq3b. Predictions were based on daily SSC values averaged over 6 months between 2010–2018 (wet season: Nov–Apr; dry season: May–Oct).

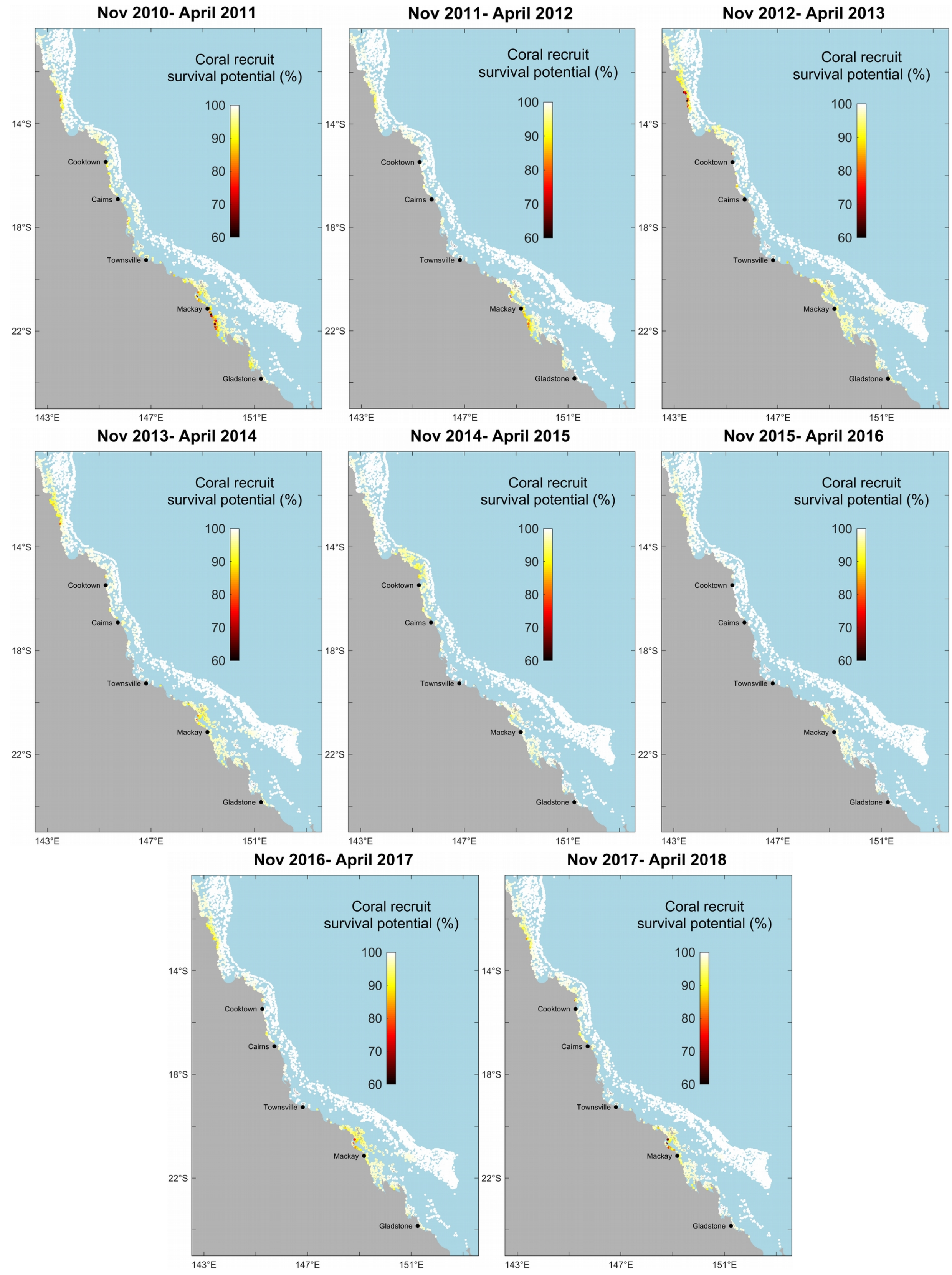


**Fig. S5.** GBR 2010–2018 predictions of the survival potential of coral acroporid recruits (six-month old corals) from concentrations of suspended sediments during the wet (summer) season.

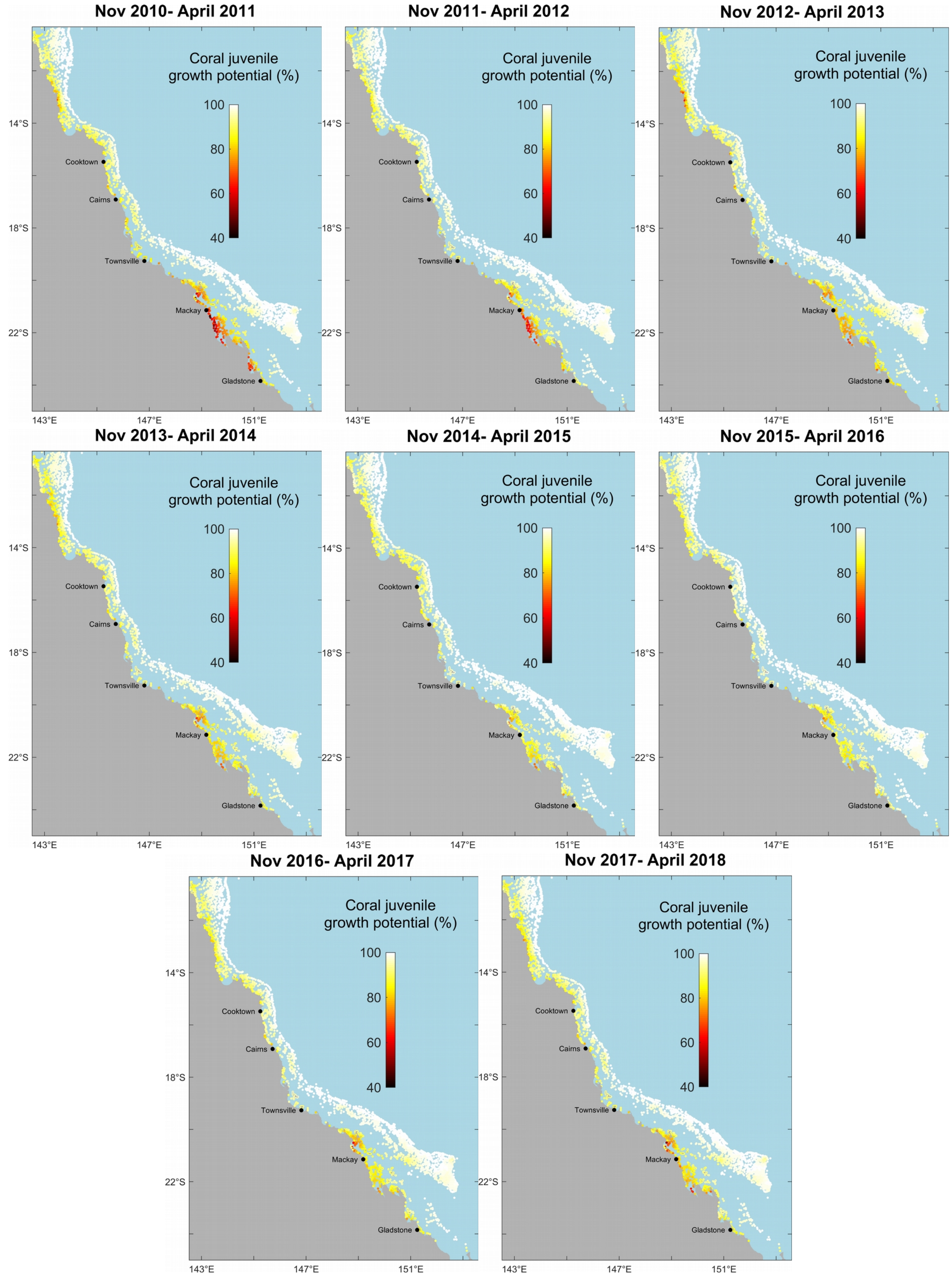


**Fig. S6.** GBR 2010–2018 predictions of the growth potential of coral juveniles from concentrations of suspended sediments during the wet (summer) season.

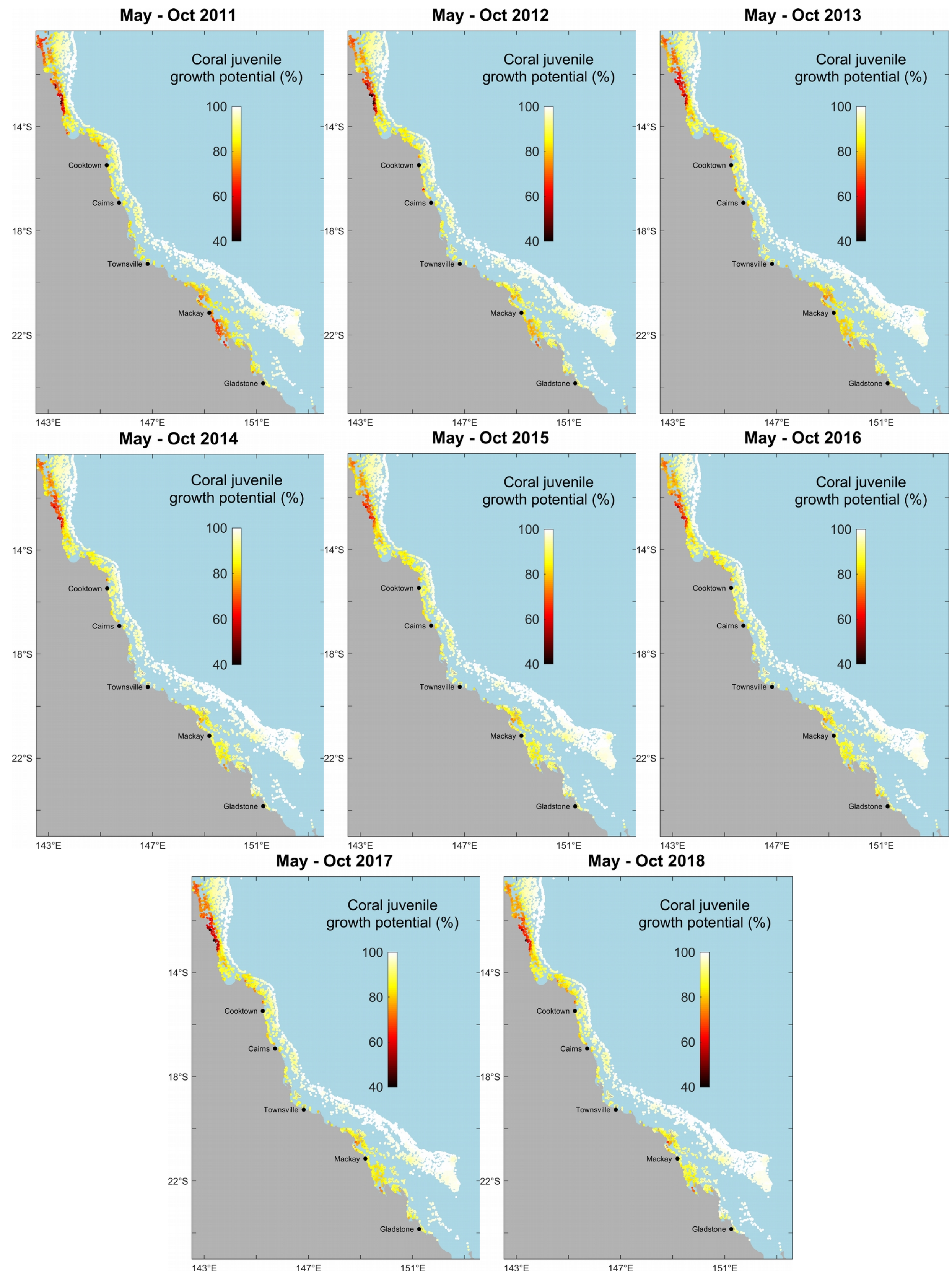


**Fig. S7.** GBR 2011–2018 predictions of the growth potential of coral juveniles from concentrations of suspended sediments during the dry (winter) season.

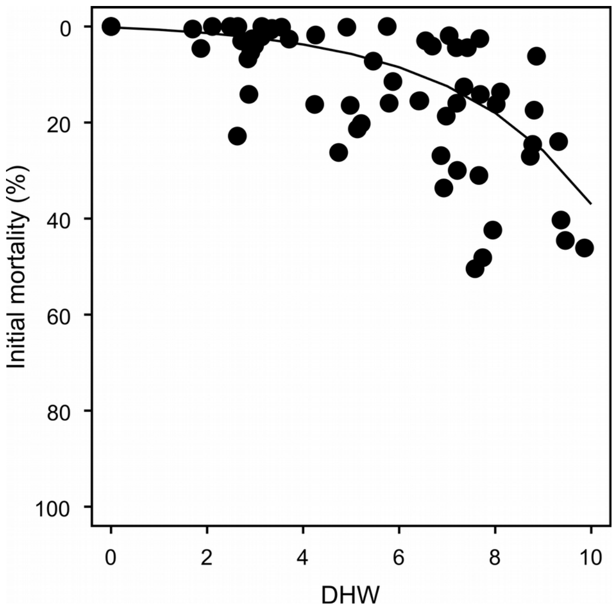


**Fig. S8.** Modelling of coral mortality following mass coral bleaching. Initial coral mortality (dots) recorded at the peak of the 2016 bleaching events by Hughes et al. (2018) in the Northern GBR, fitted with a linear model (modelled variable is $\log_e$(mortality+1), $R^2 = 0.45$).

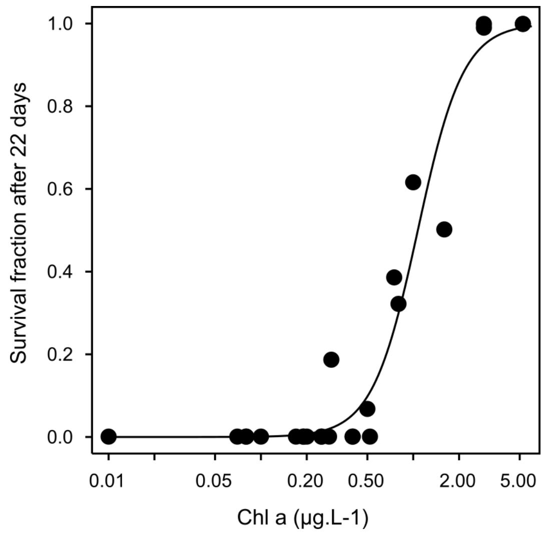


**Fig. S9.** Dose-response of the survival of crown-of-thorns starfish (CoTS) larvae to chlorophyll concentrations reproduced after Fabricius et al. (2010). Data points represent the mean proportion of surviving starfish that completed their development at age 22 days for increasing concentrations of chlorophyll *a* (eight experiments from Okaji 1996, with various enrichment treatments combined). The response curve was fitted by logistic regression (*glm* function of the *stats* R package) weighted by the number of larvae in each experiment (100 or 150 larvae). Logistic equation:
$y = 1 / (1 + \exp(0.20 - 2.91 \cdot \log x))$.

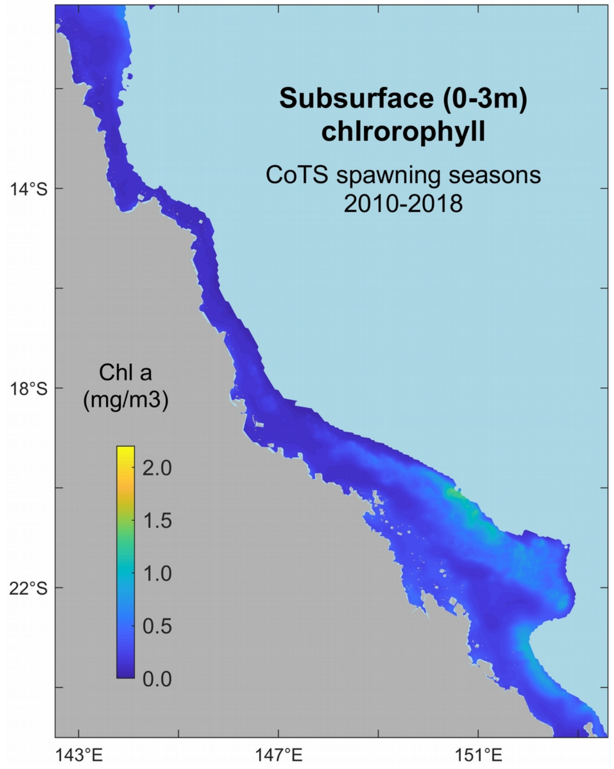


**Fig. S10.** Predictions of subsurface (maximum within 0-3 m depth) concentrations of total chlorophyll *a* at 4km resolution from eReefs averaged over eight CoTS spawning seasons (Dec–Feb 2010–2018).

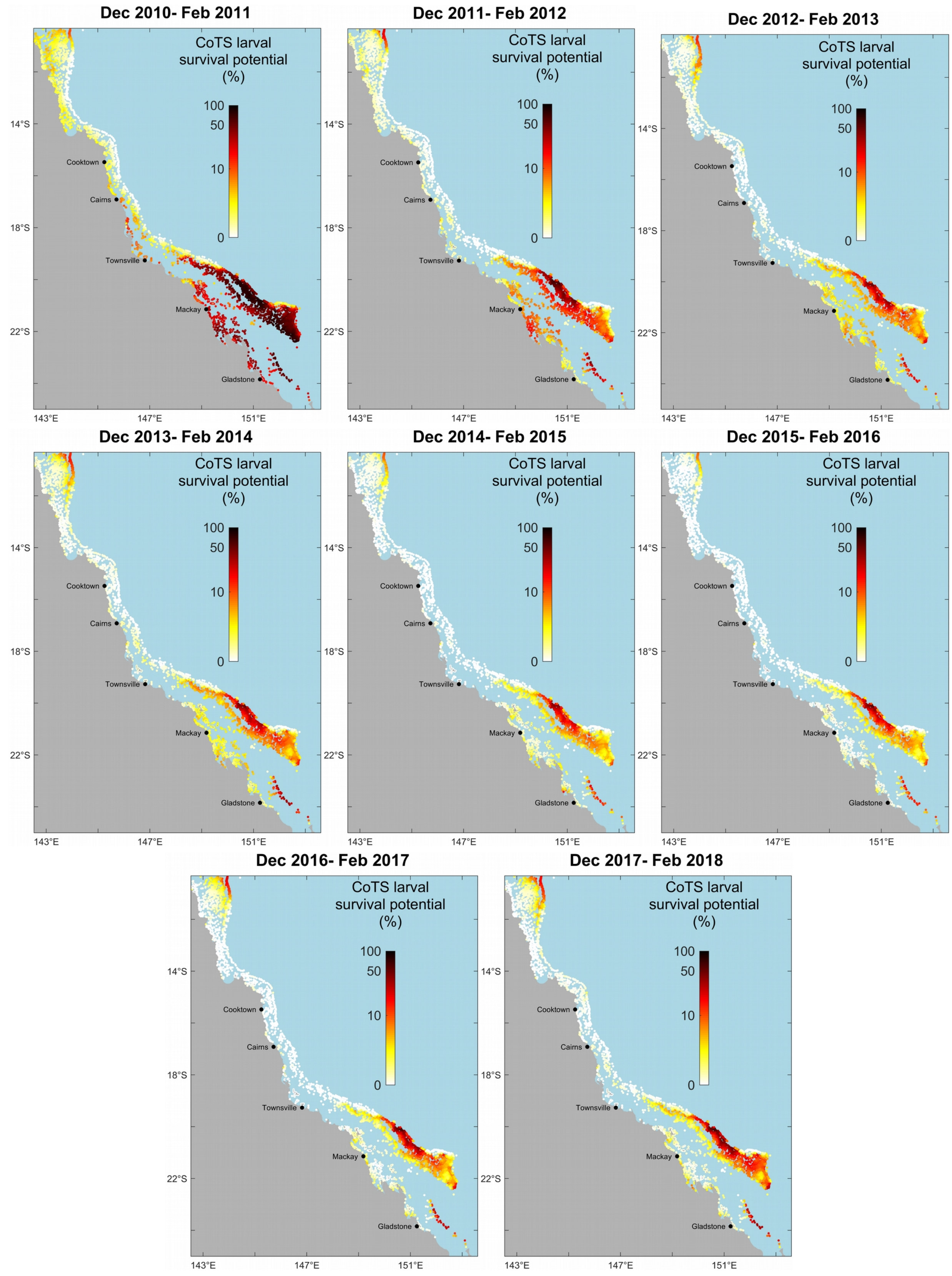


**Fig. S11.** GBR 2010–2018 predictions of the pre-dispersal survival potential of CoTS larvae (on a logarithmic scale) from total chlorophyll *a* concentrations during spawning (Dec–Feb).

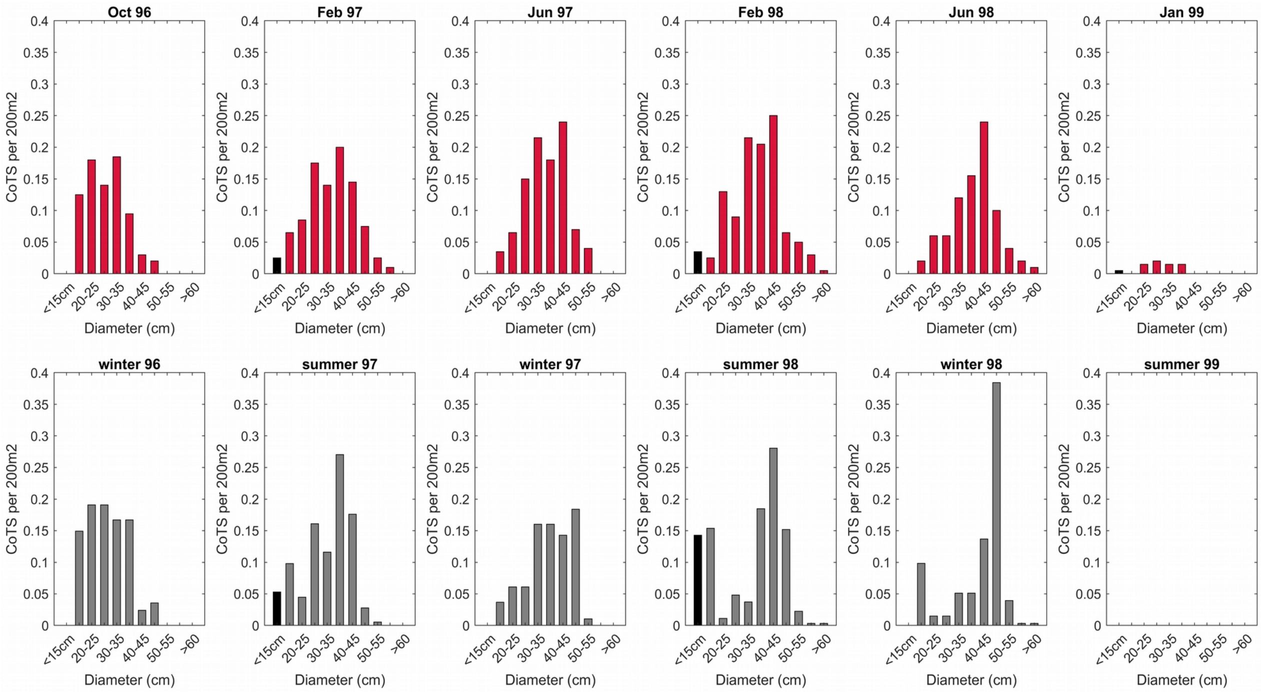


**Fig. S12.** Calibration of modelled CoTS outbreak dynamics and impacts on corals using empirical observations from Lizard Island (Pratchett 2005). Top (red bars): temporal changes in the frequency distribution of individual sizes (starfish maximum diameter) as observed on 200 $m^2$ transects. Bottom (grey bars): simulated size distribution after appropriate age-size conversion (Engelhardt et al. 1999). The model was initialized with the observed CoTS density and size distribution at the beginning of the time series. While the smallest size class (<15 cm) is likely under-represented in the survey, a reasonable match was obtained between simulated and observed density of sub-adult and adult starfish by adjusting mortality rates and parameter β of the recruitment (Beverton-Holt) function (see details in text).

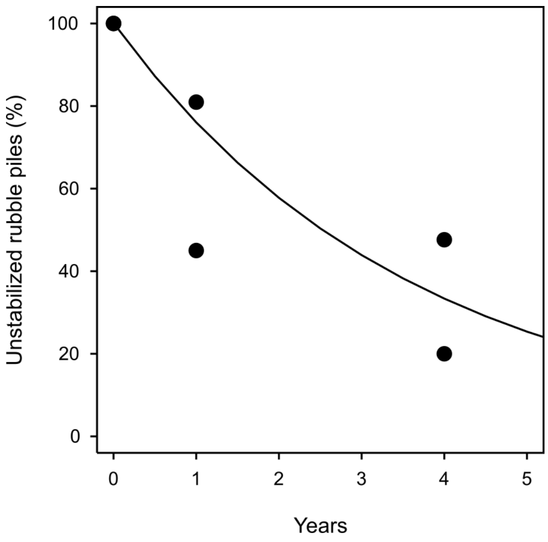


**Fig. S13.** Modelling of the natural stabilization of loose coral rubble as a simple exponential decay (solid line) against empirical observations (dots) of rubble stabilization from *in situ* experiments in two reef sites of Curaçao, Netherlands Antilles (Biggs 2013). Data points represent the changing proportion over time of experimental rubble piles that showed no sign of stabilization over time. Biggs (2013) followed ~20 piles of fragments of branching *Acropora* over 4 years, recording the number of piles that became stabilized by turf algae.

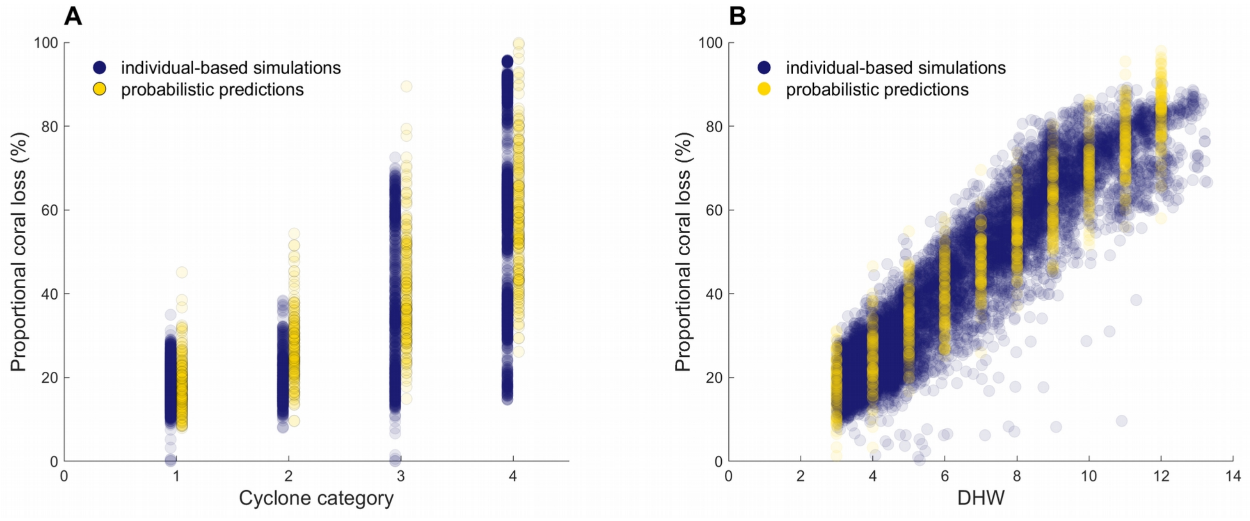


**Fig. S14.** Comparison of proportional coral cover losses generated by the individual-based simulations (ReefMod 2008-2020 hindcast, $n$ = 3,806 reefs × 13 years) and the probabilistic predictions (GLMs fitted to the hindcast cover loss, $n$ = 1,000) for different cyclone categories (**A**) and levels of heat stress (**B**). The GLMs were fitted to the proportional loss of total coral cover extracted from the hindcast simulations using the 2008-2020 exposure layers of cyclones and heat stress (DHW) as predictors. Probabilistic predictions were obtained from $n$ = 1,000 random generations of cyclone categories and DHW values.

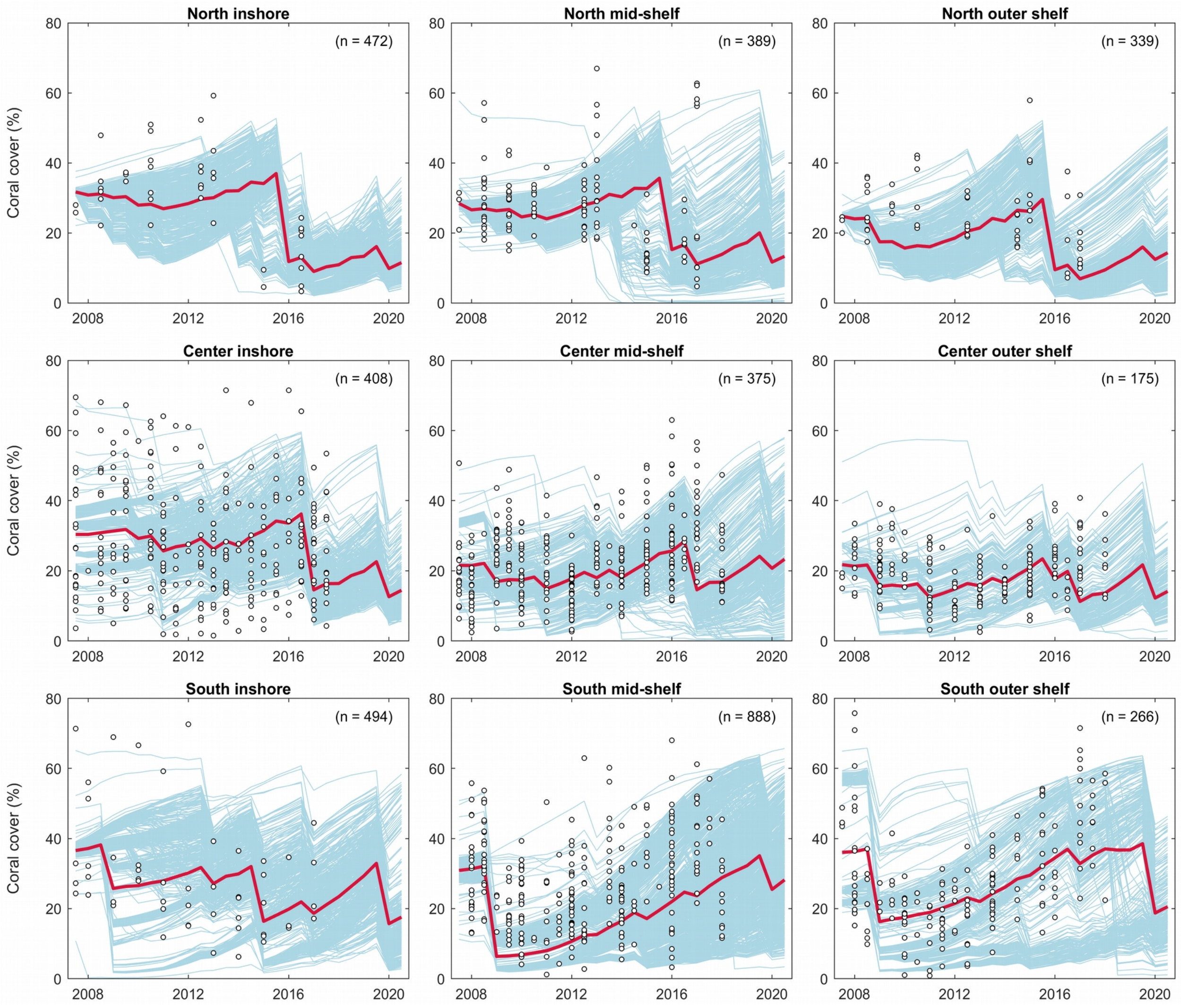


**Fig. S15.** 2008-2020 reef individual trajectories (blue lines) averaged over 40 replicate simulations for each shelf position (inner-shelf, mid-shelf and outer-shelf) in the Northern, Central and Southern GBR. Data points indicates observations of coral coverage from AIMS monitoring (transects and manta tow surveys combined). The red lines correspond to the regional average trajectory for each shelf position weighted by reef area.

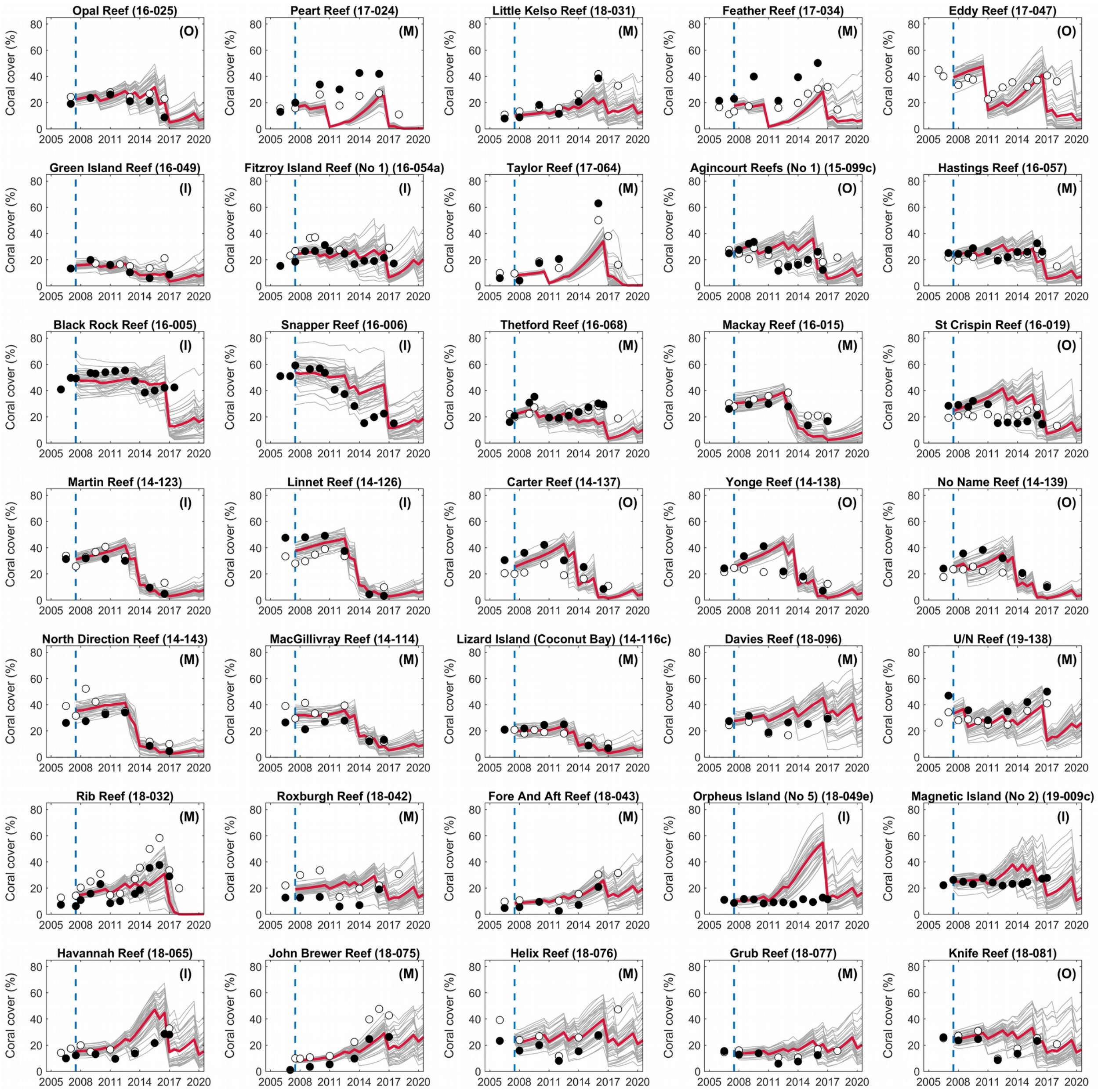


**Fig. S16A.** Validation of the reconstructed trajectories of coral cover with field observations from the AIMS LTMP (filled circles: video-transects; open circles: standardized manta tows). Reefs selected for validation (n=67) gather at least 12 surveys during 2009–2020.

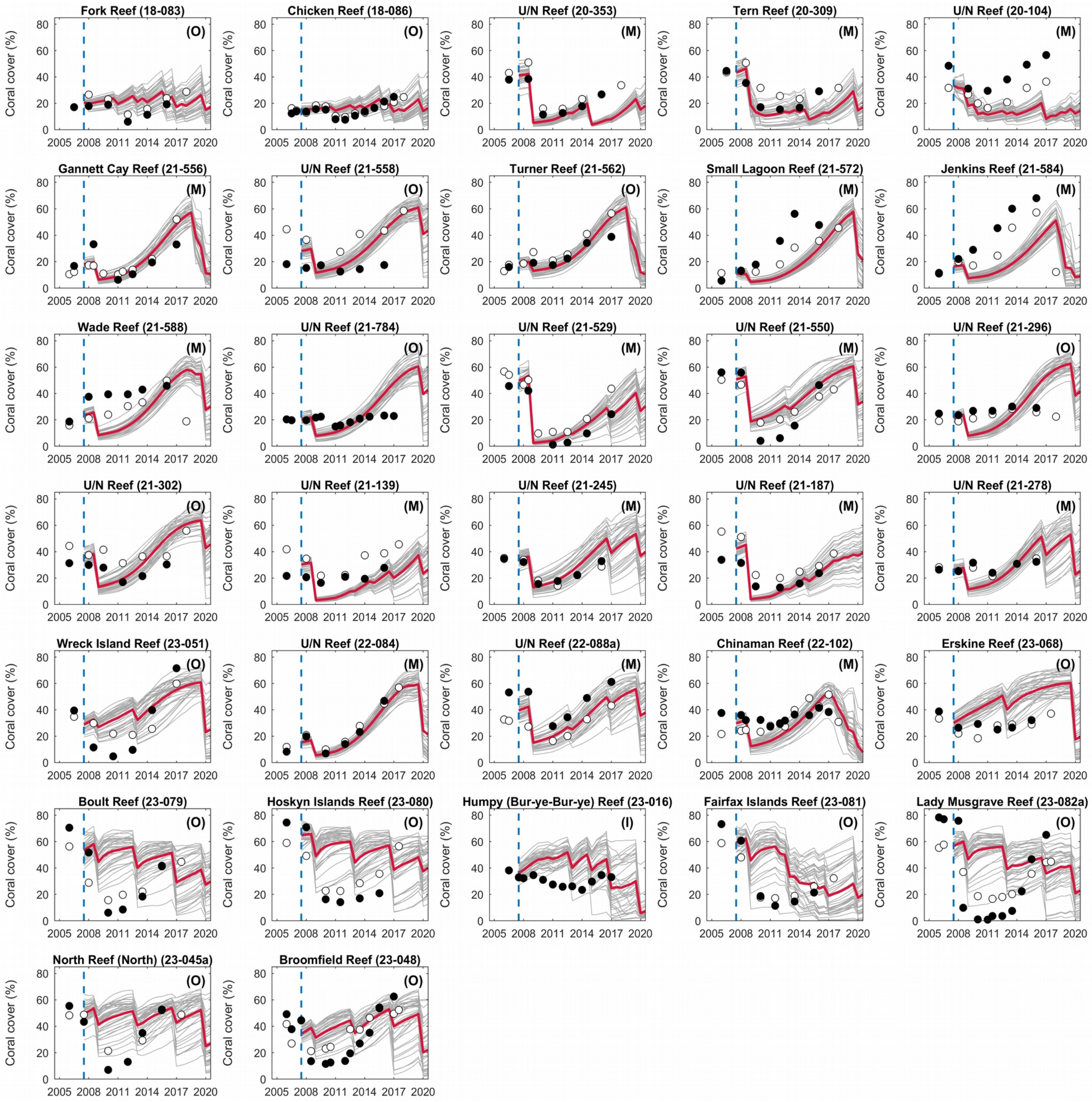


**Fig. S16B.** Validation of the reconstructed trajectories of coral cover with field observations from the AIMS LTMP (filled circles: video-transects; open circles: standardized manta tows). Reefs selected for validation (n=67) gather at least 12 surveys during 2009–2020.

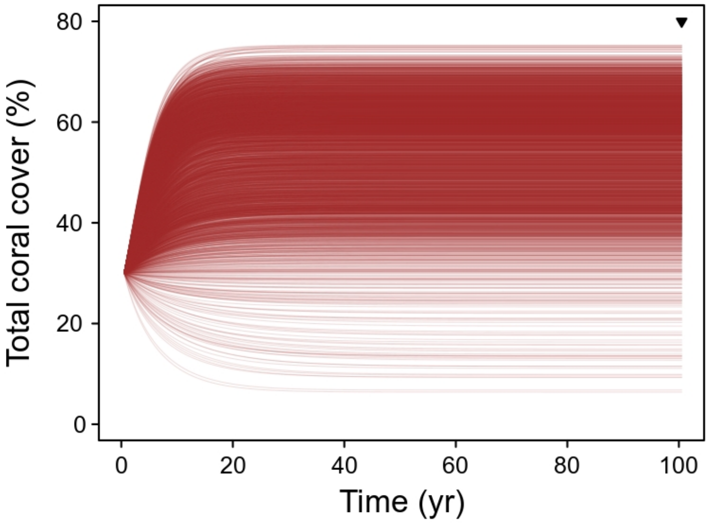


**Fig. S17.** Deterministic simulation of percentage coral cover for every reef (*n*=3,806) using a dynamic model of growth and mortality (equation 15). Growth is the increment in percent cover (*g*) predicted for a given coral cover value and reef-specific patterns of water quality and larval connectivity (sand and rubble cover fixed to 30% and 11%, respectively). Mortality represents the average annual mortality (converted to 6 months in simulations) due to all acute disturbances (cyclones, bleaching and CoTS) for the represented regimes (cyclones: 1970–2011; bleaching: 1998–2020; CoTS: 2008–2020). The cursor indicates values retained for approximating the equilibrial covers. Each individual reef has a single equilibrium (i.e., independent of starting cover) which reflects the adverse effects of coral growth and mortality determined by the local environment.
